# A glyoxal sensing *Pseudomonas aeruginosa* transcription factor enables lung infection

**DOI:** 10.64898/2026.09.20.752901

**Authors:** Christopher J. Corcoran, David G. Glanville, Bonnie Cuthbert, Rodger de Miranda, Jocelin Martinez, Johnathan D. Keith, Peter Prevelige, Susan E. Birket, Celia W. Goulding, Andrew T. Ulijasz

## Abstract

Aldehydes are a class of normally unwanted toxic electrophilic compounds that mainly arise from oxidation of glucose, lipids or DNA. However, it has recently come to light that they can also be weaponized by professional phagocytes to kill engulfed bacteria. How microbes subvert these assaults remains largely enigmatic. Here we describe the function, atomic structure and mechanism of the first bacterial transcription factor able to directly sense the dicarbonyl glyoxal (GO), which we aptly named the <u>Glyo</u>xal <u>R</u>egulator (GloR, from *Pseudomonas aeruginosa*PAO1), We show that GloR directly senses GO through a reversible cysteine modification that results in its binding to a conserved DNA regulatory motif (a *glo box*), which then triggers a transcriptional activation of a defined set of genes to help counter GO toxicity and enable acute lung infection. Despite substantial evolutionary divergence, when unmodified *gloR* and a *glo box*-regulated reporter were transferred into *E. coli*, a strikingly tight GO-specific regulation was maintained, suggesting this system could be readily transferred between unrelated microbial species. As homologs of GloR were identified in diverse bacterial species we anticipate its use to be widespread in both pathogens and environmental bacteria. Taken together, we present the first *bona fide* bacterial aldehyde regulator which senses host GO to enable survival during infection.

**Significance Statement:** Aldehydes are weaponized by the immune system to eliminate bacterial pathogens, yet little is understood about how pathogens sense and respond to these toxic metabolites. Here we describe GloR from the Gram-negative pathogen *Pseudomonas aeruginosa*, the archetype for a novel family of LysR-type transcription factors which respond specifically to the aldehyde glyoxal (GO). Through a reversible modification mechanism, GloR specifically senses GO using a conserved cysteine (C191) ligand binding pocket residue to enable the GO response. We show GloR and C191 are required for *P. aeruginosa* infection in a rat lung model of pneumonia. GloR homologs were identified in diverse bacterial species, suggesting its relevance in nature is widespread. This work paves the way to identify other bacterial sensory systems that responded to environmental aldehydes.

---

**Keywords:** Aldehydes, transcription factor, infection, *Pseudomonas aeruginosa*, regulation.

## Introduction

Reactive electrophilic species (RES) are naturally occurring toxic byproducts of metabolism which are understudied compared to reactive nitrogen species (RNS) or reactive oxygen species (ROS), yet are far more stable and can be just as damaging to cells (1). Aldehydes are a chemically diverse group of RES that are of biological relevance in all forms of life. One important class of aldehydes are the α-oxoaldehydes glyoxal (GO) and methylglyoxal (MGO), which are dicarbonyls produced by the oxidative degradation of glucose, lipids and DNA. In general, aldehyde accumulation by both prokaryotes and eukaryotes results in significant cellular malfunction and/or cell death. In humans, aldehydes have been implicated in diseases related to aging such as diabetes, cancer, and an array of neurodegenerative conditions including Parkinson’s and Alzheimer’s disease (2–7). However, recent research indicates that human cells also utilize aldehydes in the innate immune response (8, 9). In order to eliminate phagocytosed pathogens, activated macrophages endure temporary aldehyde increases through downregulation of aldehyde remediation enzymes, such as glyoxalases, and metabolic reprogramming (8–10). To counter these assaults, pathogens such as *P. aeruginosa* have evolved both detection and remediation systems (11).

The bacterial response to aldehyde exposures has been mainly limited to studies in non-pathogenic *E. coli* (12). From these studies, enzymes were discovered dedicated to MGO and GO detoxification, called ‘glyoxalases’ or ‘Glo enzymes’, which are also functionally conserved in human GLO homologs (13). More recent studies have determined that Glo enzymes can indeed protect pathogens from phagocytic killing and facilitate infection (9, 14, 15). Further, our own research has shown a link between GO and quorum sensing in *P. aeruginosa*, where GO addition initiates synthesis of an operon encompassing the major glyoxalase GloA2 and ArqI, an antibiotic monooxygenase domain that directly regulates biosynthesis of *P. aeruginosa* quorum sensing molecules (11). However, the receptor(s) that directly respond to host aldehyde assaults in *P. aeruginosa* and other bacteria remains largely unknown.

Here we describe the first *bona fide* family of aldehyde regulators in bacteria, the **<u>Gl</u>**<u>y**o**</u>xal **<u>R</u>**egulator (**GloR**) family, and its unique and highly selective mechanism of sensing and responding specifically to GO in *P. aeruginosa* to enable host survival.

## Results

### GloR (PA0709) is induced by GO and required for GO resistance

In a previous study we found that GO treatment of *P. aeruginosa* (and not any other additives or aldehydes, including MGO) resulted in induction of the *arqI-gloA2* operon and other downstream genes (PA0711-PA0714), but the regulator responsible was unknown (11). We noticed that a predicted LysR-type transcription factor, PAO1 locus number PA0708, was divergently transcribed to *arqI-gloA2* (**Fig. 1A**). To assess if PA0708 was indeed the GO-responsive transcription factor (here on referred to as <u>Glyo</u>xalase <u>R</u>egulator or ‘GloR’) we examined WT MPAO1 (16), a *ΔgloR* mutant, and *gloR* complement strain (*ΔgloR*::*gloR*) for GO versus MGO susceptibility using an efficiency of plating assay (11). Importantly, MGO differs from GO by only an additional methyl group. Data in **Fig. 1B** (above) shows a marked effect on GO susceptibility in the Δ*gloR* mutant, but not in the complement nor with addition of MGO. We next tested if the *gloR* deletion alone exhibited similar GO-mediated killing to the *arqI-gloA2* operon observed previously (11). **Fig. 1B** (below) indeed shared parallel killing between *ΔgloR* and *ΔarqI-gloA2* strains, and therefore suggested that GloR is epistatic to the ArqI-GloA2 GO resistance signaling and remediation cascade.

**Figure 1.**
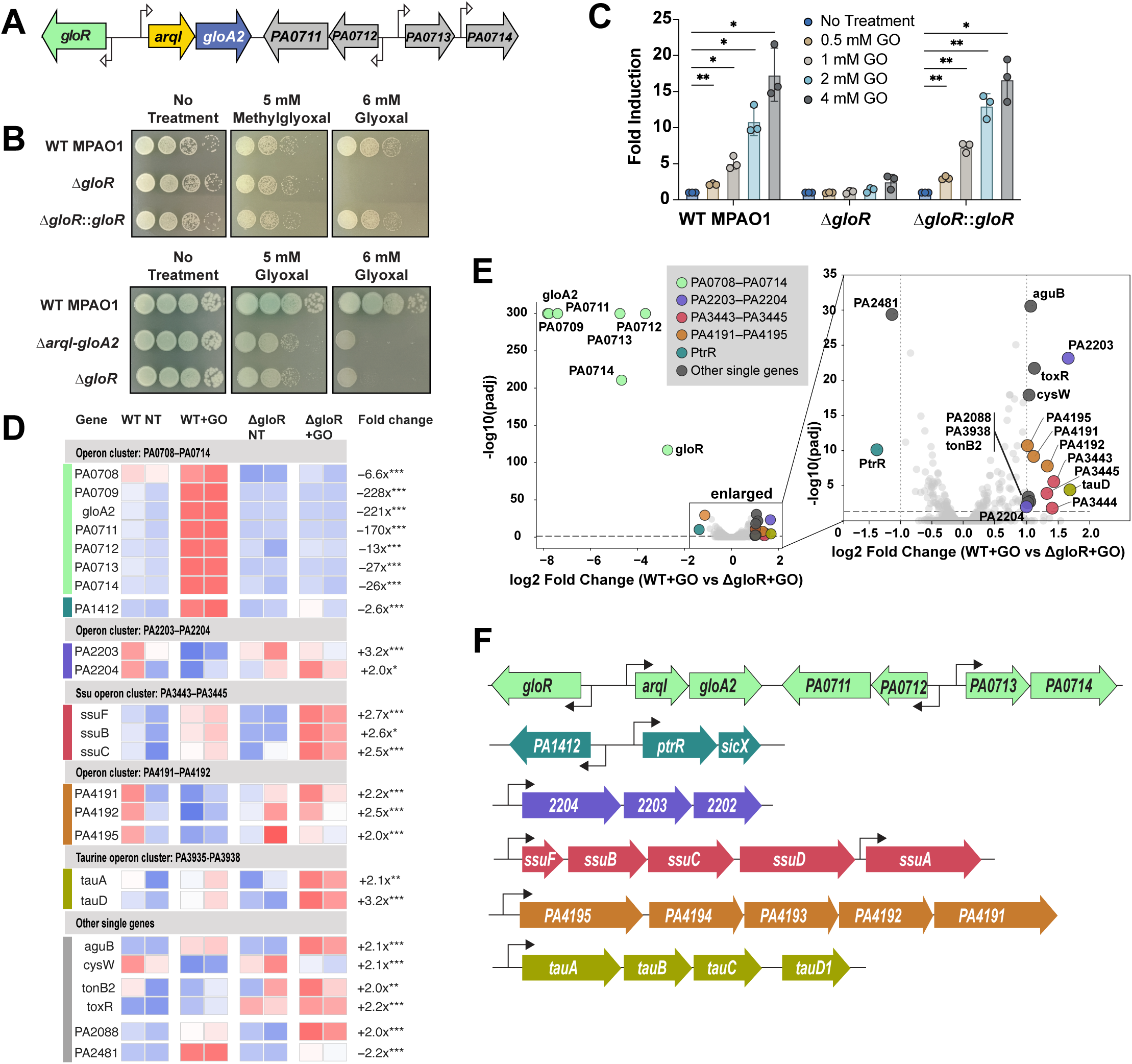
GloR regulation and its role in GO resistance. (**A**) Schematic diagram of the *P. aeruginosa* genomic region containing *gloR*, the *arqI-gloA2* operon and other nearby GO-regulated genes. (**B**) Spotted MPAO1 cells after GO, MGO, or no treatment with WT MPAO1, *ΔgloR*, and *gloR* complement *(ΔgloR::gloR*) expressed *in trans* as a FLAG-tagged version from replicating plasmid pMMB67EH (pMMB67EH-GloR-FLAG, which is IPTG inducible (57)) (above), or with the *ΔarqI-gloA2* operon deletion strain (10) (below) using 1:10 dilutions. Figure is representative of 3 biological repeats. (**C**) Promoter activity after addition of increasing amounts of GO measured by using a dual fluorescent reporter system (17) with the experimental mScarlet-I signal being driven by the *arqI-gloA2* promoter in a WT MPAO1, *ΔgloR*, and pMMB67EH-GloR-FLAG complement. Statistics were performed using a one-sided two-sample t-test against untreated control. *, p < 0.05; **, p < 0.01. Experiments were done on three separate days. (**D**) Volcano plot of RNA-seq results (average of n = 2 experiments performed on separate days) comparing WT MPAO1 or Δ*gloR* strains after GO addition for 15 minutes (+GO) or not treated (NT). Plot on the right is an expansion of the boxed points on the left. Dotted line indicates the limit of plotting for the adjusted p-value (Padj). (**E**) Heatmap of individual experiments depicted in *D*. The eight leftmost columns show per-replicate row z-scores of log2 (normalized counts + 1) across all four conditions (WT NT, WT +GO, Δ*gloR* NT, Δ*gloR* +GO), colored on a diverging blue (low), white (middle) and red (high) scale. Colored bars on the left match the colors in *D* of the operons in question. The rightmost numeric column reports the fold-change and significance of the WT vs Δ*gloR* comparison under GO treatment for each gene. Asterisks indicate DESeq2 Wald-test adjusted p-value: *p < 0.05, **p < 0.01, ***p < 0.001. (**F**) Schematic representation (not to scale) of genome sections of *P. aeruginosa* determined by RNA-seq to be influenced by GloR. Arrows indicate predicted promoters that were identified using PromoterHunter (19).

We next tested if GloR could induce its regulon in a GO-specific manner. We first tested the GloR’s ability to induce the *arqI-gloA2* promoter (P*_arqI_*) by exploiting our dual fluorescent reporter system (pCG-VmS; (17)) to monitor P*_arqI_* activity by creating plasmid pCG-V_P*arqI-gloA2-*_ mS. pCG-V_P*arqI-gloA2-*_mS was then transformed into wild-type, *ΔgloR* and *ΔgloR*::*gloR* MPAO1 strain backgrounds. In rich broth, P*_arqI_* activity increased in a dose-dependent manner when GO was added to the WT reporter strain, whereas in the *ΔgloR* mutant GO responsiveness was negligible, and could be restored after complementation (**Fig. 1C**). When taken together, these results indicated that GloR was a primary regulator of the GO resistance response in *P. aeruginosa*.

### Mapping the GloR regulon

After establishing that GloR induced the *arqI-gloA2* operon we wanted to examine if it had a more expansive role in gene regulation. To do this we performed RNA-seq and compared gene expression profiles of WT MPAO1 versus Δ*gloR* MPAO1 cell with or without a 15-minute incubation with 2 mM GO (a concentration we previously determined was below the MIC (11)). Results indicated little significant difference between WT and *ΔgloR* samples in the absence of GO, suggesting that under these *in vitro* conditions GloR does not have a regulatory role in the absence of GO (**Dataset S1**). In contrast, in the presence of GO samples demonstrated a defined set of genes that were GO-GloR dependent, where the strongest effect was seen in the *arqI-gloA2* operon and downstream operons/genes of unknown function *PA0711-PA0714* (**Fig. 1D-F; Table S1; Dataset S1**).

Other genes that were modestly affected in the presence of GO and were GloR dependent included sulfur metabolism genes/operons *tauD* and *tauA*, *ssuC and ssuD*, and *cysW*, as well as two operons that encode predicted transporters (*PA2203*-*4*, *PA4191*-*5*; **Fig. 1E**; **Table S1; Dataset S1**). Another transporter, *PA1412*, was shown to be positively regulated by GloR. *PA1412* is divergently transcribed from an adjacent two-gene operon containing a GloR homolog (PtrR, see below; **Fig. 1F**) and the small regulatory RNA *sicX* that controls the oxygen-dependent switch between chronic and acute infection (18) and was previously shown by our us to be repressed 14-fold after GO addition (11), suggesting that GO signaling might be epistatic to *sicX* and thus involved in virulence transformation. Once again using our dual reporter (17), RNA-seq data was further confirmed by assessing GloR-dependent expression of promoters controlling the *PA1412*, *PA2202-04*, and *PA4191-95* genes/operons (**Fig. 1F**) in both the presence and absence of *gloR* and increasing concentrations of GO. Data shown in **Fig. S1A** allude to an activating role for GloR in regulating both *PA2202-04* and *PA4191-95* operons, and a repressive role for regulation of PA1412. Taken together, these data validate the RNA-seq targets and confirm that GloR activates and represses transcription of multiple genes throughout the *P. aeruginosa* genome in response to GO.

### Identification of a GloR binding motif (*glo box*)

To identify a GloR consensus binding sequence we first used PromoterHunter (19) to tabulate the upstream regulatory regions in genes that were suspected of direct transcriptional regulation by GloR after examining our RNA-seq data. Sequences from a select number of GloR dependent regulatory regions (from **Fig. 1**) were then applied to MEME (20) to search for common motif(s). The reverse palindrome consensus sequence of TAATT-N_7_-AATTA was identified (**Fig. 2A**), which follows the general LysR-type transcriptional regulator family recognition sequence of T-N_11_-A (21). We named the GloR consensus recognition sequence a ‘*glo box*’. The predicted GloR binding sequence was found to lie just upstream of the predicted −35 RNA polymerase (RNAP) binding sequence within the *arqI-gloA2* and *PA1412* promoters (**Fig. 2B**), which is typical of transcriptional activators and supports GloR’s ability to activate transcription of these promoters (22). In contrast, the *glo box* in the *PA2204* and *PA4195* promoters was found to either overlap with or was located just downstream of the predicted −10 and −35 sequences (**Fig. 2B**). This placement profile strongly suggests RNAP inhibition of binding and/or processivity (23) and was therefore consistent with the RNA-seq data. We then confirmed the direct binding of GloR to these *glo box*-containing promoter regions (P*_arqI_*, P*_PA2204_*, P*_PA4195_*, and P*_PA1412_)* using electrophoretic mobility shift assays (EMSAs). Purified GloR (determined to be a dimer in higher salt by size exclusion chromatography (SEC)); **Fig. S1B, C**) was addended in increasing concentrations to fluorescently labeled promoter regions. Results indicated that GloR bound to all probes at nanomolar to low micromolar concentrations and were deemed specific by unlabeled competitor DNA addition (**Fig. S1D**).

**Figure 2.**
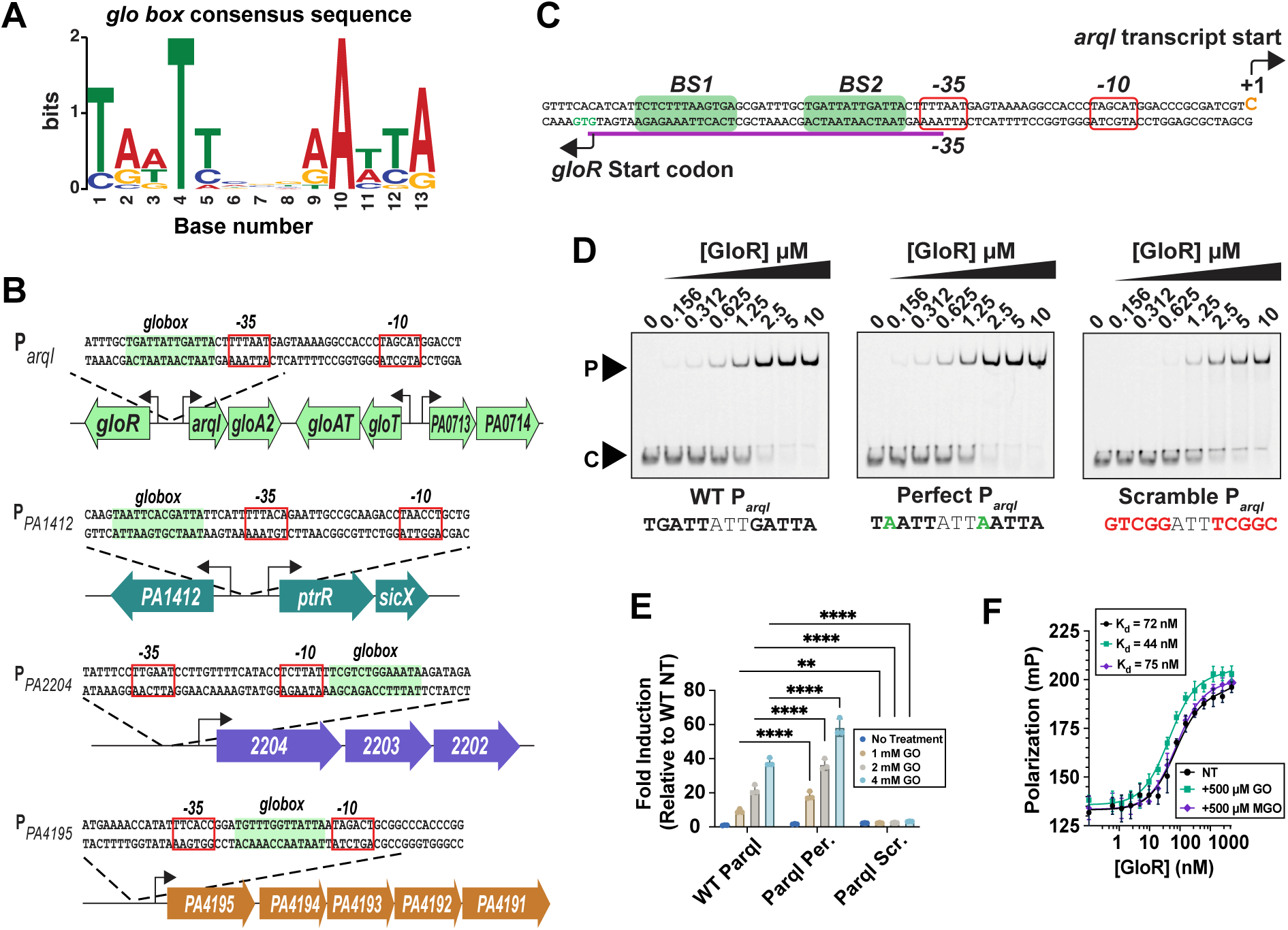
GloR regulates the GO response through its *glo box* DNA element. (**A**) HMM Logo depicting the GloR (*glo box*) consensus recognition sequence identified by MEME motif search (58). Search parameters were: one occurrence per sequence, palindromes only, given strand only, and 13-20 bps motif. (**B**) Schematic representation of *arqI-gloA2*, *PA1412*, *PA2204*, and *PA4195* operons and their partial promoters encompassing 55 bps centered around the GloR and *-10* and *-35* RNAP recognition sequences. Predicted *glo box* motifs are boxed in light green. Predicted *-10* and *-35* sequences are boxed in red. (**C***) ArqI-gloA2* upstream region showing the two predicted GloR binding sites (*BS1*, *BS2*; in green). The purple line indicates the oligo derived from the *arqI-gloA2* promoter used in EMSAs in *E* (below). (**D**) EMSAs of increasing amounts of purified GloR interacting with a fixed amount (0.5 µM) of fluorescently labeled DNA probe (WT, ‘perfect’ inverted repeat sequence, or scrambled *BS2* site). Colored nucleotides indicate where changes were made in the WT sequence. (**E**) GO induction of P*_arqI-gloA2_* WT, ‘perfect’ inverted repeat sequence, or scrambled *BS2* site dual reporter constructs in *P. aeruginosa* cells. ***, p < 0.001; ****, p < 0.0001. Statistics: One-sample t-test comparing untreated with treated samples. (**F**) Fluorescence polarization (FP) experiments presenting the binding of GloR to a 39-mer oligo containing *BS1*.

Within the P*_arqI_* region we found two potential GloR binding sites (BS1 and BS2), which were both predicted upstream of the identified −35 sequence (**Fig. 2C**). We then synthesized a 45-mer DNA double stranded oligo with an unaltered BS1 site and a scrambled BS2 to determine if GloR could still bind the oligo. In addition, a ‘perfect’ inverted palindrome sequence was used to replace BS2 in hopes of enhancing GloR interaction. EMSA analyses showed no observable difference in binding between the WT and ‘perfect’ BS2 sequence oligos (**Fig. 2D**). However, the 45 bp oligo containing the WT BS1 sequence and the scrambled BS2 resulted in a marked reduction in affinity (**Fig. 2D**), demonstrating that both BS1 and BS1 contribute to the interaction. However, when these same WT, ‘perfect’ and scrambles BS2 sequences were interrogated in the bacterial cell using a P*_arq_* reporter construct the differences were far starker, where the ‘perfect’ sequence enabled a higher induction by GO at lower concentrations and the scrambled BS2 sequence resulted in no detectable induction (**Fig. 2E**). Together, these results demonstrated that our proposed inverted palindrome is the GloR site of interaction, and that the BS2 site, which is closer to the core promoter, is crucial to *arqI-gloA2* promoter activation but only fully observable *in vivo*. This result comes as no surprise in that when multiple transcription factor binding sites are involved the site closest to RNAP usually serves as the higher affinity site, which when occupied first will then subsequently allow for a cooperative binding effect for the others to be recruited (24, 25). Finally, these results also suggest that other factors are involved in GloR regulation of target promoter sites *in vivo*.

### GO, but not MGO, increases GloR binding affinity to the *glo box*

GO and MGO differ only by a single methyl group, yet MGO does not appreciably affect GloR activity, nor does GloR confer resistance to MGO treatment (**Fig. 1**) (11). To determine if the GloR mechanism of action was due to direct binding of GO to alter its position/affinity for the *glo box*, we used an in-solution binding assay (fluorescence polarization, FP) with GloR and its labeled wild-type DNA recognition sequences encompassing both BS1 and BS2 binding sites from the *arqI-gloA2* promotor (**Fig. 2C**). Upon addition of GO, GloR affinity for its DNA increased from 72 to 44 nM Kd, whereas addition of MGO had no appreciable effect (75 nM Kd; **Fig. 2F**). These results support the hypothesis that GO increases GloR affinity for the *glo box* regulatory sequence. The small shift in affinity could be reflective of how LysR transcription factors have been shown to respond to ligand binding by simply repositioning themselves on the DNA to alter DNA bending which, in turn, changes RNAP recruitment, rather than canonical regulation where the ligand confers more outright DNA interaction (21, 26). This is further supported by the fact that GloR is able to bind the *arqI-gloA2* promoter at relatively high affinity in the absence of GO (**Fig. 2D**).

### The GloR-*glo box* signaling axis is functionally transferable to a heterologous system

Although we demonstrated that GloR could induce gene expression in *P. aeruginosa*, it was unclear whether GloR required an accessory *P. aeruginosa*-specific factor to initiate transcription other than RNAP. To answer this question, we used an *E. coli* dual plasmid system where the *gloR* open reading frame was cloned into one expression plasmid and placed under the control of an IPTG-inducible promoter. The second plasmid was the pCG-P*_arqI-gloA2_*-mS reporter (17) (**Fig. 3A**). Surprisingly, signal was only detectable when GO was added, which revealed a dose-dependent transcriptional activation (**Fig. 3B**). These results show that the GloR-*glo box* signaling axis is all that is required to transfer tightly controlled promoter induction to a genetically divergent bacterial species.

**Figure 3.**
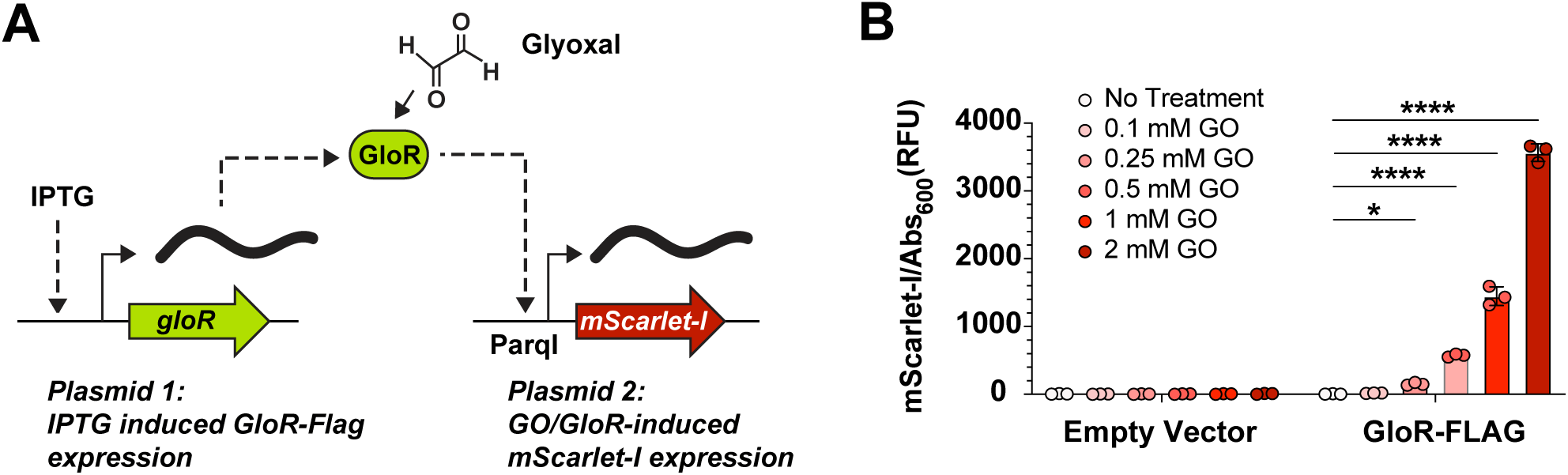
GloR regulation is transferable into a heterologous system. (**A**) Schematic diagram of the two-plasmid heterologous *E. coli* reporter system. GloR-FLAG is induced from the first plasmid (pMMB67EH) with IPTG. GloR then binds to P*_arqI_*, driving mScarlet-I expression on the second plasmid pCG-V_P*arqI-*_mS (17) upon GO induction. (**B**) *E. coli* DH5α cells harboring the pCG-V_P*arqI-*_mS mScarlet-I reporter and pMMB67EH-GloR-FLAG or pMMB67EH empty vector control were treated with increasing concentrations of GO. Expression of GloR-FLAG resulted in increased reporter signal upon GO exposure in a dose-dependent manner. Statistics were calculated using a two-way ANOVA test. *, p < 0.05; **, p < 0.01; ****, p < 0.0001.

### The atomic structures of the GloR ligand binding domain (GloR_LBD_)

To understand potential mechanism(s) of the GloR response to GO we initiated structural studies. The ‘native’ GloR_LBD_ was purified to near homogeneity (**Fig. S2A**) and its oligomeric state was determined to be a dimer by size exclusion chromatography (SEC; **Fig. S2B**). Selenomethionine-labelled (SeMet) GloR_LBD_ crystals were obtained, and the structure of GloR_LBD_ was determined to 2.5 Å resolution (statistics are given in **Table S2**). The GloR_LBD_ asymmetric unit (ASU) contained four subunits (**Fig. 4A**; **Fig. S2C**) in a ‘dimer of dimers’ tetramer oligomeric state which is common for LysR proteins (21, 26). Each subunit contained two globular domains encompassing a 5-stranded mix β-sheet and four α-helices (**Fig. S2D**). The dimerization interface, between subunits A&C or subunits B&D, comprised ∼1000-1200 Å^2^ on the surface of each subunit (∼11-12% of subunit surface) (27), and appeared to reflect the physiological dimeric interface of GloR as a ‘head-to-tail’ dimer that is also typical of many LysR family transcription factors (21).

**Figure 4.**
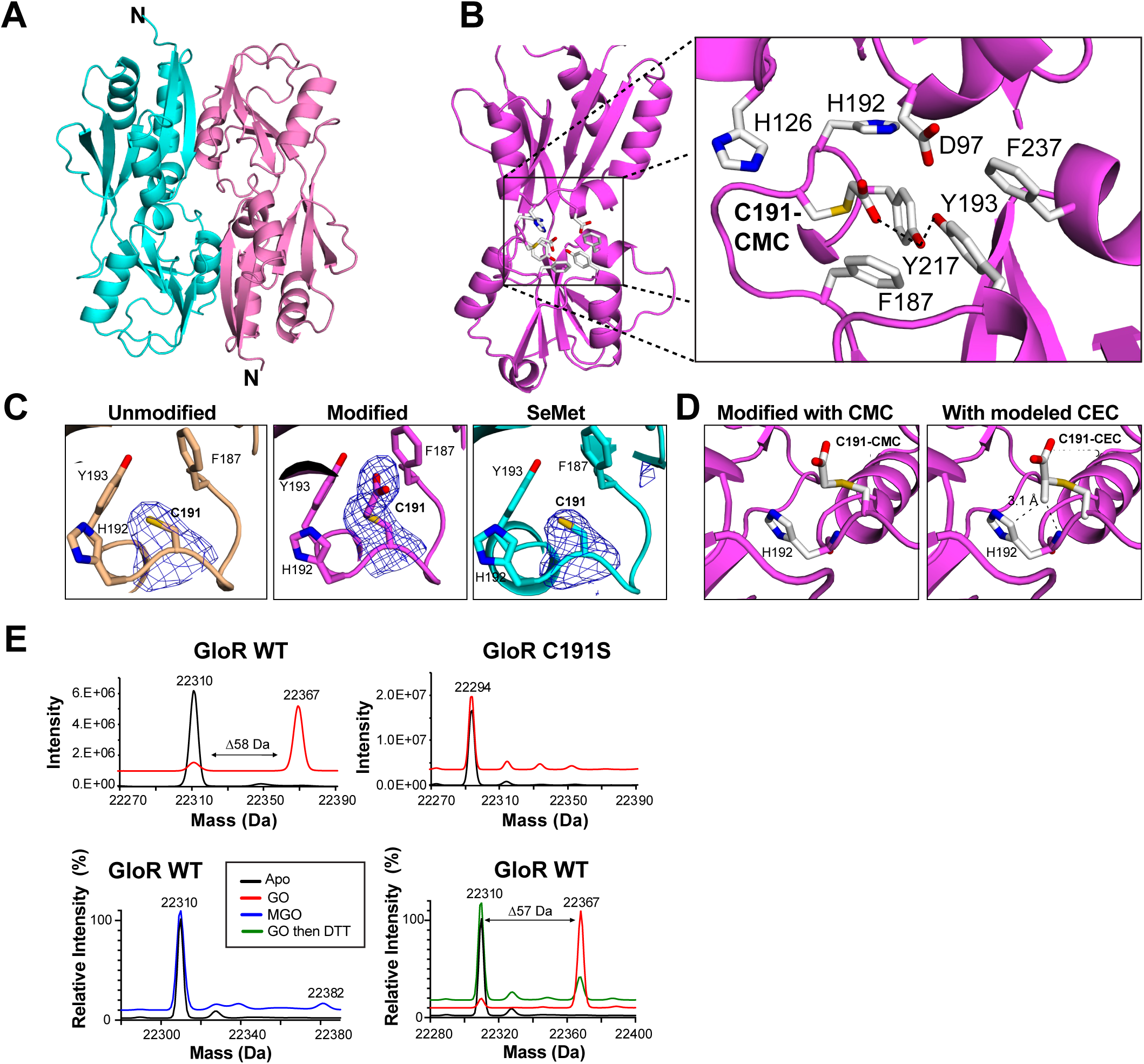
GloR_LBD_ atomic structure and pocket residue contribution to function. (**A**) Cartoon of the unmodified GloR_LBD_ dimer showing the head-to-tail packing common to the LysR transcription factor family. Monomers are shown as pink and cyan structures. (**B**) Closeup of the GloR GO-modified structure highlighting conserved ligand binding pocket residues (white sticks). Dotted lines denote predicted hydrogen bonding. (**C**) Comparative omit maps of C191 for the unmodified (brown), modified (pink), and SeMet (cyan) structures. (**D**) Modified structures with GO CMC modification and a modeled MGO-derived CEC modification on C191. Dotten lines show the distance from the MGO methyl group is 3.1 Å away from the His192 backbone nitrogen and from the histidine imidazole ring Cd2 carbon atom. (**E**) Intact mass spectrometry spectra of purified WT GloR_LBD_ sample with GO, MGO, GO and then DTT added, or no addition run after a 20-minute incubation. Spectra are also displayed for the GloR_LBD_ C191S mutant after GO or no GO addition. Numbers given above peaks are calculated masses. Abbreviations: GO, glyoxal; MGO, methylglyoxal; DTT, dithiothreitol; WT, wild-type; CMC, carboxymethylcysteine. CEC; carboxyethylcysteine.

We next co-crystallized GloR_LBD_ in the presence and absence of GO and solved the structures to 2.1 Å and 2.6 Å resolution, respectively (**Table S2**). Surprisingly, when we compared the native structure to the structure in the presence of GO, we observed ‘extra’ electron density extending from the sulfur atom of Cys191 in the GO treated (modified) structure, which was not present in the untreated (unmodified) structure (**Fig. 4B**; **Fig. S2E, F**). Significantly, modelling a S-carboxymethyl cysteine (CMC; (28)) covalent PTM fulfilled the electron density (**Fig. 4C**). The unmodified and modified GloR_LBD_ crystals formed a nearly identical unit cell with two subunits in the ASU, aligning with a RMSD of 0.6 Å across 392 residues (**Fig. S2G**). The SeMet structure was nearly identical to the modified and unmodified structures (1.1 Å across the dimers), except across the loop bearing residues 241-247 (**Fig. S2G** denoted by the arrow). In the SeMet structure these residues were unstructured in 2 of the 4 subunits, as evidenced by an absence of electron density.

The modified Cys191 (CMC) helps form the canonical LysR ligand binding pocket (21), comprising residues Asp97, His126, Phe187, Cys191, His192, Tyr193, and Tyr217 (**Fig. 4B**). In the SeMet structure the ligand binding pocket was occupied by an acetate molecule from the crystallization solution, which formed hydrogen bonds to one of two conformations of His126 mapped in this structure (**Fig. S2H)**, and in the unmodified structure Cys191 makes direct contact with His126. However, in the modified structure the acetate is replaced by two water molecules, which form hydrogen bonds with neighboring Tyr193 and Tyr217, and to accommodate the CMC PTM on Cys191 the His192 imidazole ring was rotated to prevent a steric clash (**Fig. S2I**). To understand how this pocket might exclude the highly related MGO molecule, we modeled a MGO modification (a carboxyethyl cysteine (CEC) adduct) in place of the GO modification. Although a *bona fide* steric clash was not observed with MGO, its methyl group was ∼3 Å away from Cδ2 and the mainchain nitrogen of His192, suggesting that this residue might play a role in aldehyde selectivity (**Fig. 4D**).

### Mass spectrometry (MS) verification of the GloR_LBD_ PTM

To verify the CMC-191 PTM we performed liquid chromatography mass spectrometry (LC-MS) on both GO and MGO treated GloR_LBD_. A GO treated sample showed the expected change in mass for a CMC modification of +58 Daltons (Da), whereas a C191S mutant or addition of MGO mutant did not (**Fig. 4E; Fig. S3A**). Interestingly, the mass shift observed in the WT could be reversed by addition of the reducing agent dithiothreitol (DTT; **Fig. 4E; Fig. S3A**). Because a CMC modification is irreversible these data suggested that the adduct was likely a hemithioacetal; a GO intermediate seen in glyoxalases that is easily reversed in the presence of a reductant like DTT, but that can subsequently rearrange to form the irreversible CMC adduct seen in the GloR crystal (29). Importantly, both hemithioacetal and CMC adducts maintain the same mass shift (Δ58 amu), making them virtually indistinguishable by MS.

These data showed that modification of GloR is specific to GO, and that the PTM is likely, initially, a reversible cysteine hemithioacetal adduct. The hemithioacetal adduct reflects glyoxalase *in vivo* chemistry (30) as an irreversible modification would result in suicide-type inactivation. Modelling hemithioacetal adducts for both GO (CMC) and MGO (CEC) into the modified structure revealed that both adducts were easily accommodated in the active site, as modelling did not give any steric clashes (**Fig. S3B**). Together these findings suggest a complex *in vivo* mechanism of GloR signaling mediated by an initial unstable modification of C191 to a hemithioacetal adduct, enabling downstream signaling to bind DNA.

### Contribution of GloR ligand binding site residues to its activation

After solving the structure of the GloR_LBD_ we wanted to interrogate the contribution of GloR ligand binding pocket residues to activation of the *arqI-gloA2* operon and resistance to GO. To accomplish this we first deleted *gloR* in its native chromosomal location (*ΔgloR*) and replaced it with FLAG-tagged versions of wild-type *gloR*, and D97A, H126A, F187A, H192A, Y193A, Y217A, C191A, C191S, and C191D variants to create *in cis* native-locus complementation strains. Importantly, an anti-FLAG western blotting of GloR-FLAG WT and variants showed similar expression levels (**Fig. S4A**). The strains were then transformed with the pCG-P*_arqI-gloA2_*-mS reporter plasmid and activity was measured by mScarlet-I signal normalized to constitutive sfGFP after GO or MGO addition, or no treatment control.

Results indicated that all residues were critical for GO induction of P*_arqI-gloA2_*, with the notable exceptions of the Y193A mutant that was not significantly different from WT, and also the H192A mutant that resulted in a reduction in P*_arqI-gloA2_* activity rather than complete abolition (**Fig. 5A**). Interestingly, the Y217A and Y193A mutants showed altered selectivity. Both mutants result in statistically significant increased MGO-dependent activity compared to WT, where Y193A could now be induced by both GO and MGO, and the Y217A could only be induced by MGO (albeit induction by MGO did not reach the levels of GO induction; **Fig. 5A**). These data support our structural modeling and indicate that the two tyrosine residues, Y217A and Y193A, as well as His192, could be involved in excluding other aldehydes.

**Figure 5.**
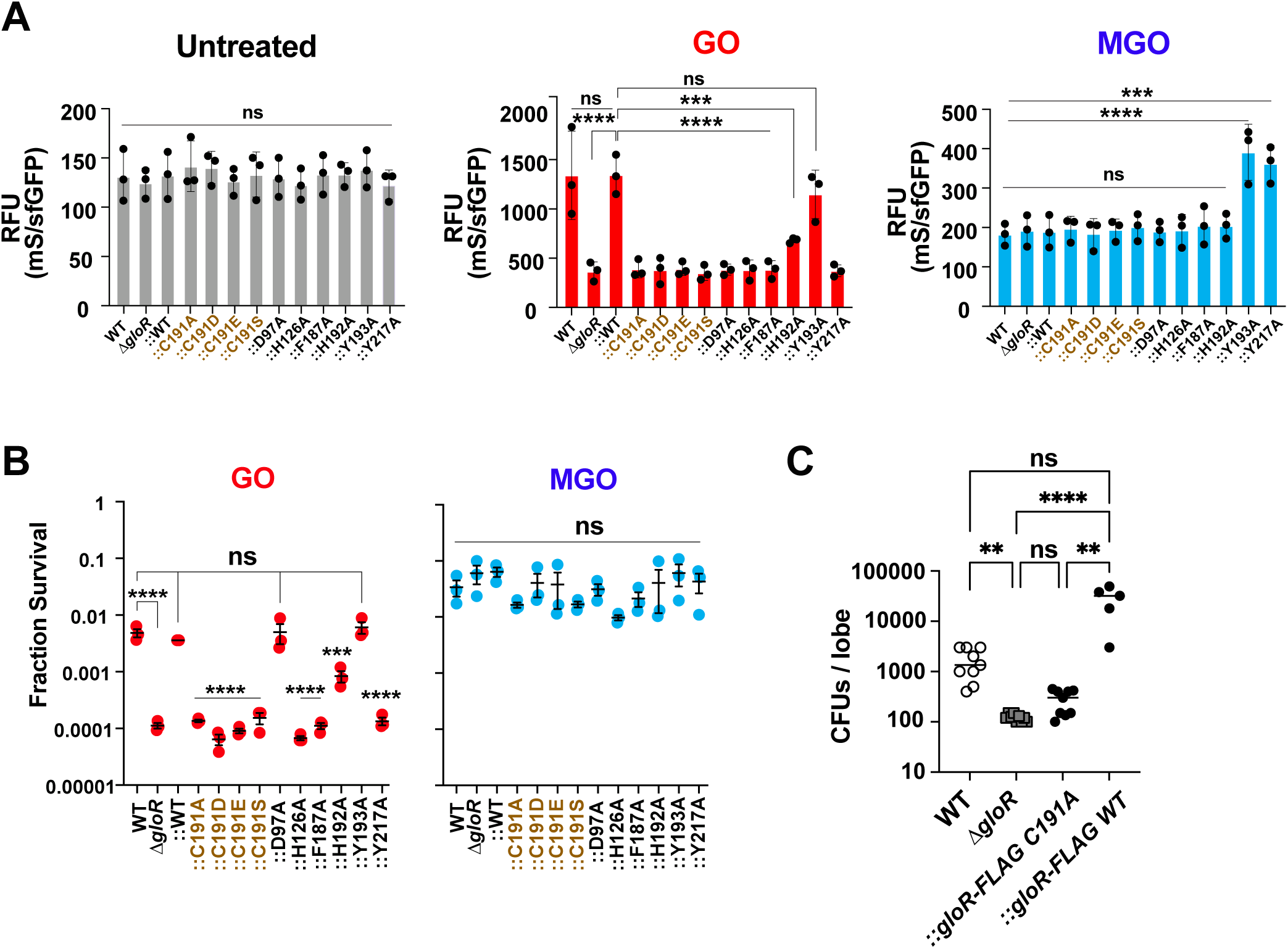
*In vivo* analyses of GloR pocket residues and infectivity. (**A**) Examination of GloR isogenic mutants and their effect on P*_arqI-gloA2_* induction by GO and MGO using our dual reporter system (17). (**B**) Aldehyde toxicity assays of WT, *ΔgloR*, and *in cis* complemented WT *gloR* and *gloR* variants. Assays were run in triplicate (3 technical repeats) on three separate days (biological repeats). Bars indicate SEM. Statistics were calculated using a one-way ANOVA test followed by a Dunnett’s multiple comparisons text. ***, p <0.001; ****, p < 0.0001. ns, not significant. (**C**) Acute infection of rat lung. 3 x 10^6^ MPAO1 *P. aeruginosa* cells were used to inoculate rat tracheas by direct intratracheal installation. Infections were allowed to proceed for 72 hours before animals were euthanized and lung homogenates plated to determine remaining bacterial CFUs. For statistics, a one-way ANOVA with Kruskal-Wallis test was applied. **, p < 0.01; ****, p < 0.0001; ns, not significant. The bar denotes the median. Each dot represents a single animal. “::” denotes a knock-in/markerless replacement of the WT or variant *gloR* versions at the native *P. aeruginosa* chromosomal location (*gloR* gene locus number in the PA01 genome is *PA0708*).

Subsequently, we performed GO versus MGO killing activity assays with the GloR ligand-binding site variants. Here the diminished CFUs generally paralleled the P*_arqI-gloA2_* reporter results (**Fig. 5B**, **Fig. S4B**), where GO addition to C191, H127, F187, and Y127 mutants exhibited similar CFUs in comparison to the Δ*gloR* strain. The H192A mutant showed partial resistance to GO, and the Y193 mutant reflected WT CFU counts (**Fig. 5B**, right panel). Interestingly the D97A GloR variant diverged from P*_arqI-gloA2_* reporter experiments, where this point mutant displayed WT CFU levels (**Fig. 5B**, right panel), yet was incapable of inducing promoter activity (**Fig. 5A**). The reason for this discrepancy is not obvious but could be due to the D97A GloR constitutively inducing another unknown GO resistance locus. Notably, MGO addition exhibited no statistical differences in killing between mutant and WT strains (**Fig. 5B**, left panel), which is in line with our previous report that the *arqI-gloA2* operon is not involved in MGO resistance (11, 17). These results further support the importance of the ligand binding site residues surrounding the C191 PTM for GloR function.

### GloR and its C191 sensory residue are critical for lung infection

*P. aeruginosa* is a well-known lung pathogen, especially in the context of chronic infection of the CF lung (31), and acute, non-CF hospital-acquired and ventilator-associated pneumonia (32). To test the importance of GloR and GO resistance in *P. aeruginosa* infection we used an acute rat model of pneumonia with WT, *ΔgloR*, and both WT and C191A *in cis* native-locus (FLAG-tagged) complementation strains of *P. aeruginosa*. Both the *ΔgloR* and *gloR* C191A mutant complement strains exhibited approximately an order of magnitude reduction in CFUs compared to WT MPAO1, whereas, as expected, the WT *gloR* complement control was not statistically different from WT MPAO1 cells (**Fig. 5C**). These results clearly indicate that GloR and its conserved GO-modifiable C191 residue are required for *P. aeruginosa* acute infection of the lung, and in more general terms emphasize the importance of GO sensing and remediation to enable pathogen infection (11).

### Evolution and conservation of GloR

GloR belongs to the extensive, yet largely functionally unassigned LysR family of bacterial transcription factors (21). To better understand the evolution of GloR and how it clusters within the LysR family, we used BLAST with full-length GloR to obtain its closest homologs. A list of 198 non-redundant sequences was obtained, all of which contained the conserved GloR C191 (**Dataset S2**). A phylogenetic tree generated from a sequence alignment (Clustal Omega (33); **Fig. S6**) suggested that GloR evolved from the LysR regulator PtrR, which has recently been described as regulating succinate semialdehyde dehydrogenase expression, and has also been shown to function as a putrescine stress response regulator (34). Interestingly, here we show that GloR binds within the regulatory region of the *ptrR-sicX* operon (**Fig. 2**), suggesting these two closely related LysR transcriptional factors are not only homologous, could also be functionally connected.

To examine differences in key ligand binding pocket amino acids between GloR and PtrR, the *P. aeruginosa* PtrR homolog (PA1413) was aligned with that of GloR using both a primary sequence alignment (**Fig. S6A**) and a superimposition of the modified GloR_LBD_ structure and AlphaFold (35) predicted PtrR_LBD_ structures (**Fig. S6B**). In PtrR, the GloR H126 and H192 residues have been replaced with a proline and serine, respectively, and D97 with a tyrosine, whereas all other important ligand binding pocket residues described above for GloR were maintained (**Fig. S6**). From these observations, and the fact that we can switch a single ligand binding pocket residue to enable partial MGO induction (either Y217 or Y193; **Fig. 5**), we can conclude that minor changes within the LysR family pocket can easily enable the evolution of divergent ligand accommodation, whilst maintaining the overall signaling process.

## Discussion

Here we describe GloR, its mechanism and regulatory activities (summarized in **Fig. 6**); an archetype for the first prokaryotic transcription factor family that has been mechanistically proven to directly sense an aldehyde. Whereas other bacterial transcription factors have been associated with such a response, *e.g.* NemR, YqhC, NsrR, and Fnr from *E. coli* (36), AldR from *Bacillus subtilis* (37), and CmrA from *Pseudomonas aeruginosa* (38), these examples have been more tethered to general stress responses and/or indirect aldehyde sensing, and therefore the precise ligand and mechanism have remained elusive. For example, *E. coli* NemR utilizes a cysteine-mediated redox-sensing mechanism to respond to a wide range of RES (*e.g.* GO, MGO and quinolones), with the formation of intermolecular disulfide bonds leading to its activation (39–41). Like GloR, YqhC and CmrA were shown to respond to GO, but also many other chemically diverse compounds (36, 42, 43). While these studies have implicated certain transcription factors within the greater RES/aldehyde general ‘toxicity response umbrella’, a *bona fide* receptor which directly responds to an aldehyde and the mechanism by which signaling occurs through atomistic analyses has not been demonstrated. Interestingly RAGE, the eukaryotic aldehyde-sensing receptor is activated by direct MGO modification, but also accommodates many other ligands (44), therefore making GloR’s specific sensitivity unique even by standards set though the better studied, analogous human receptor systems.

**Figure 6.**
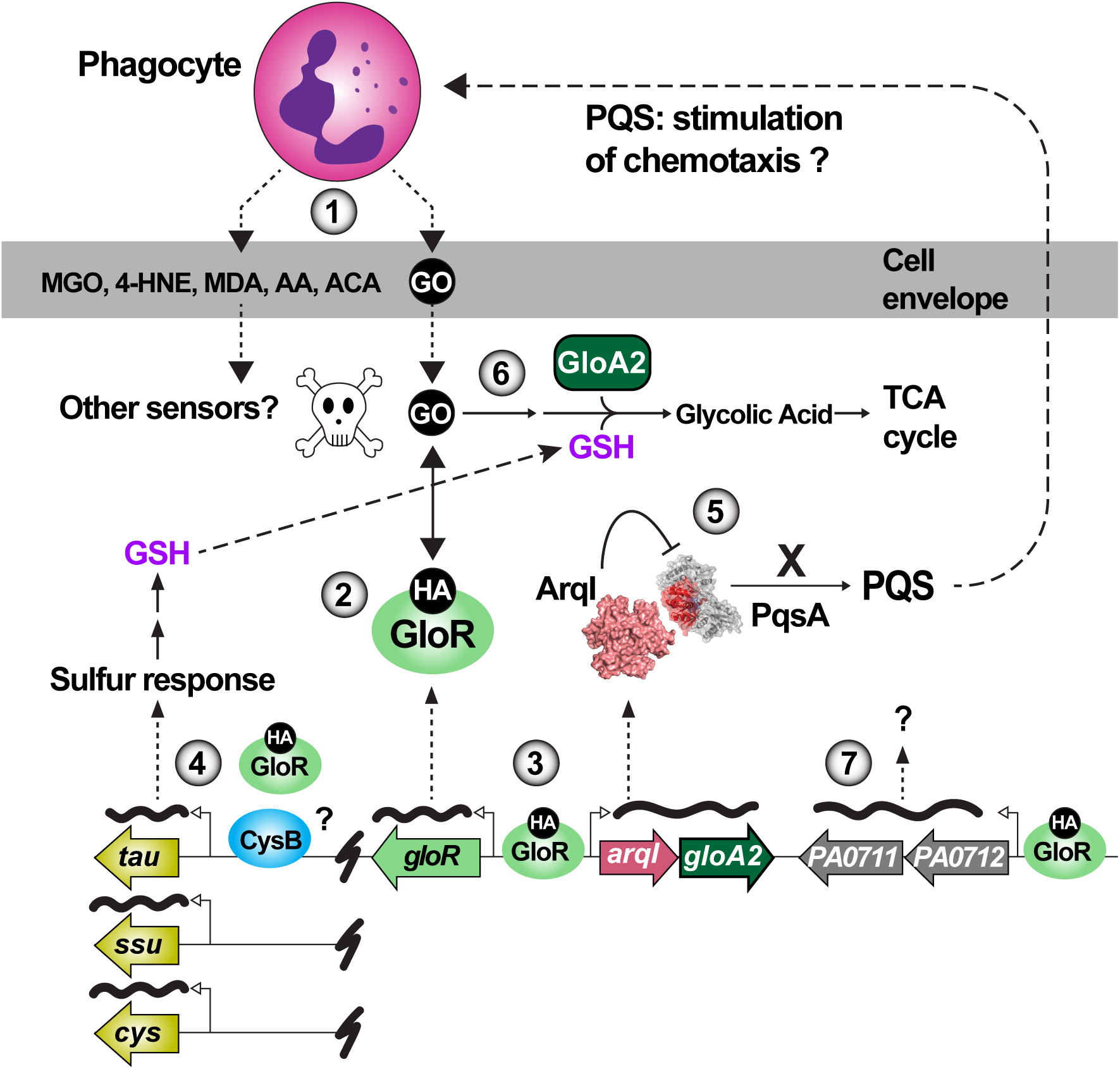
**Schematic diagram of proposed GloR function**. (1) GO is produced by phagocytes through increased glycolytic flux, lipid peroxidation, and DNA oxidation. Infected macrophages often downregulate detoxification systems like GLO1 and DJ-1 to ensure these compounds accumulate and damage bacterial proteins and DNA. Many other aldehydes are also produced which currently have no known bacteria sensing systems. 4-HNE, 4-hydroxy-2-nonenal; MDA, malondialdehyde; ACA, acetaldehyde; AA, arachidonic acid derived aldehydes. (2) GO diffuses into the bacterial cell and reacts with Cys191 on GloR to form a hemithioacetal (HA) adduct which enables (3) binding to the *arqI-gloA2* operon promoter to initiate RNA synthesis. Conversely, we propose that a (4) relief of GloR binding allows an as-of-yet undescribed factor to then initiate RNA synthesis (*e.g.* CysB) and induction of sulfur response genes (*tau*, *ssu* and *cys* operons). Sulfur response proteins enable glutathione (GSH) synthesis which is required to detoxify GO by glyoxalases (*e.g.* GloA2). (5) ArqI then directly binds PqsA, the first enzyme in PQS biosynthesis (11), to block PQS signaling (a quorum sensing molecule that has been shown to stimulate phagocyte chemotaxis (70)). (6) The GloA2 glyoxalase detoxifies GO using GSH as a substrate to create the less toxic glycolic acid, which can then enter the TCA cycle as a carbon source. (7) GloR also upregulates the adjacent operon *PA0711-PA0712* which is a predicted toxin-antitoxin (TA) system of unknown function. The roles of PA0713 and PA0714 are also currently unknown.

One of the most intriguing findings in this report is the PTM on C191, which is required for GloR activation and virulence. Recent reports have indicated that many proteins might undergo reversible cysteine-specific GO/MGO modification which can have real biological consequences (45), and that many of the proteins that harbor such GO/MGO modified cysteines have glyoxalase activity which has gone unrecognized (46). Some explanations for these shortcomings could be the instability of the hemithioacetal, its identical mass to CMC and inevitable conversion to CMC with time which creates a difficult problem when assessing its presence biochemically (e.g. harsh mass spectrometry pretreatments) and with structural biology procedures. Indeed, atomic structures exhibiting CMC modifications have often been presumed artifacts (*e.g.* see ref. (47)). Some of these PDB entries include: (i) the regulatory CAAX box cysteine on some small G-proteins in the RAS superfamily (48), (ii) the catalytically important cysteine residues on glyceradehyde-3-phosphate dehydrogenase (GAPDH) (49), and (iii) Park-7 (also referred to as DJ-1). Although these CMC modifications have been presumed to be artifacts of mass spectrometry (50) or crystallographic processes, our data here and a recently published article (45) support the alternative hypothesis that these cysteines might, in fact, exhibit previously unmapped and biologically relevant glyoxalase activity. In the specific example of the DJ-1/Park7 cysteine 106 residue that is essential for Parkinson’s disease progression, this residue is now accepted as having redox sensing capabilities which ultimately translate to a role in the maintenance of mitochondrial function and neuronal homeostasis (51). Subsequent studies show that both bacterial and vertebrate DJ-1 (Park7) homologs act as glyoxalases or ‘de-glycating’ enzymes whose roles are to reverse both GO and MGO damaging glycation of proteins (52, 53). It is therefore likely that DJ-1/Park7 shares a similar hemithioacetal intermediate with GloR and other proteins with glyoxalase enzymatic activity.

Also paralleling our GloR studies here (**Fig. 5**), a recent report has connected host DJ-1/Park7 activity to host immunity, showing that its presence hinders macrophage-driven bacterial clearance in mice (54). Although this result might at first sound counterintuitive, activated M1 macrophages have now been shown to require a temporary increase in aldehyde concentrations (such as MGO) to kill phagocytosed bacteria (8, 9), which would be greatly assisted by the absence of the vertebrate DJ-1 glyoxalase. Indeed, our work has shown that, from the perspective of the invading pathogen *P. aeruginosa*, aldehyde exposure strongly upregulates the *P. aeruginosa* DJ-1 homolog (over 25-fold; (11)), presumably to eliminate excess cellular aldehydes like GO/MGO originating from phagocytes. Although we did not see DJ-1 differentially regulated in our RNA-seq studies with GloR, one could hypothesize that DJ-1 and other enzymes besides GloA2 are upregulated by GloR during infection in retaliation to the host weaponizing aldehydes as antimicrobials. Taken together, one can envision a host-pathogen battle during phagocyte capture whereupon the host is trying to increase aldehyde production while the pathogen is, in turn, attempting to blunt these assaults. Given the absolute requirement for the presence of GloR C191 for infection, and by inference its CMC PTM for signaling, it is interesting to speculate that many more bacterial and host proteins which respond similarly assaults by other phagocytic-produced aldehydes, especially ones produced from oxidation such as 4-hydroxy-2-nonenal (4-HNE), malondialdehyde (MDA), and acetaldehyde (**Fig. 6**). Indeed, a recent publication has implicated aldehydes and other toxic byproducts from the oxidation of arachidonic acid in the phagocyte-mediated killing of *Staphylococcus aureus* (55).

Modeling GO (CMC), MGO (CEC) and their corresponding precursory hemithioacetal adducts on C191 suggested a mechanism of how GloR might discriminate between GO and MGO in the ligand binding pocket, and strongly implicated some residues in the signaling process. With GO addition the C191 CMC PTM results in a rotation of H192, which would put a methyl group in the same position close enough to the mainchain nitrogen of H192 to possibly exclude MGO in the reaction (**Fig. 4D**). This observation suggests that H192 might play a role selectivity of GO over MGO. In further support of this hypothesis, Y193 and Y217 coordinate the PTM (**Fig. 4B**, **Fig. S2I**), and are the only residues that when replaced with alanine lead to statistically significant GloR activation with MGO addition (**Fig. 5A**). Even more interesting is the fact that a Y193A mutant enables activation from both GO and MGO (*i.e.* yields enhanced promiscuity), whereas the Y217A mutant notably switches its preference to MGO (GO no longer induces this mutant). These results seem suggestive of the H192-Y193-Y217 triad of ligand binding pocket residues playing important roles in forming a favored recognition site for GO adducts, whilst working to exclude other would-be reactive aldehydes (*e.g.* the closely related aldehyde, MGO). On the other hand, the fact that Y193 and Y217 are conserved among GloR and PtrR homologs alike, and conversely that H192 seems conserved only in GloR (**Fig. S6**), suggests a more complex picture which would include other pocket residues in dictating precise substrate specificity. With only a single residue required for substrate recognition change, one could conceive such slight changes within the GloR ligand binding pocket might have evolved to naturally accommodate other aldehydes in GloR versions from different bacterial species.

Recent publications have demonstrated the dicarbonyls MGO and GO, and likely other aldehydes produced by phagocytes (*e.g.* 4-hydroxynoneanal, malondialdehyde, and acetaldehyde), are important for bacterial killing by macrophages (9–11, 55). Here and in our previous work (11) we show that bacteria might have developed specific aldehyde sensory and remediation systems to counter these assaults. Indeed, in the important pathogen *P. aeruginosa* GloR appears to be central to this effort, as it is required for acute lung infection (**Fig. 5C**). We have previously shown that the *arqI-gloA2* operon and downstream operons/genes of unknown function are highly upregulated in the presence of GO, as are genes involved both the phosphate and sulfur starvation response, *e.g. tauABCD*, *ssuFBCD* and *cysTWA* (10, 11), whose role might be to replenish the GSH pool – a substrate for glyoxalases (**Fig. 6**). Here we show that indeed, in response to GO, GloR binds upstream of and controls the positive expression of *arqI-gloA2*. However, in contrast, here we found that GloR (slightly) *represses* the sulfur starvation response (**Table S1**), where we know from our previous studies GO *activates* the sulfur starvation response (11). One explanation for this discrepancy could be that GloR acts to compete with another unidentified *positive* regulator to limit the expression of the sulfur response in the presence of GO. A likely candidate could be another LysR regulator, CysB, which positively controls these operons in response to sulfur starvation in *P. pudia* and *E. coli* (56) (**Fig. 6**).

Given the large discrepancy of gene regulation differences between *in vitro* versus *in vivo* growth conditions, it will be of future interest to explore GloR regulation in the context of more relevant host-pathogen assays. Since we were able to transfer GloR into a heterologous background where it was fully functional (**Fig. 3**), and that many homologs we found in other species have high sequence similarity to *P. aeruginosa* GloR, it is quite possible that the GloR signaling axis could be laterally transferrable to diverse Gram-negative bacteria and enable similar GO responsive defenses. Future studies of GloR in other bacterial pathogens (**Fig. 6**) will be necessary to properly answer this question.

## Methods

### Bacterial Strains and Culture Conditions

Bacterial strains used in this study can be found in **Dataset S3**. *P. aeruginosa* and *E. coli* cultures were grown in Lysogeny Broth media (LB, Difco; Beckton Dickinson) for standard growth and maintenance. Liquid cultures were grown at 37 °C and 230 rpm on an orbital shaker unless otherwise stated. Culture density was monitored using a Genesys 150 UV-Vis spectrophotomer (Thermo Fisher Scientific) at a wavelength of 600 nm. Solid media was solidified with 1.5% agar (bacteriological; VWR). Superoptimal broth with catabolite repression (SOC) was made with 20 g/L tryptone, 5 g/L yeast extract, 0.5 g/L NaCl, 10 mM MgCl_2_, 10 mM MgSO_4_, 2.5 mM KCl, and 10 mM glucose. M9 minimal media (6.78 g/L Na_2_HPO_4_, 3 g/L KH_2_PO_4_, 0.5 g/L NaCl 1 g/L NH_4_Cl, pH 7.4) was supplemented with 0.2% glycerol 100 μM CaCl_2_, 100 μM FeCl_3_, and 2 mM MgSO_4_ after autoclaving. Pseudomonas Isolation Agar (PIA; Sigma Aldrich) was supplemented with 20 mL/L glycerol for selection of *P. aeruginosa* exconjugants. Sucrose containing plates were made by the addition of 5% sucrose to LB agar after autoclaving. Antibiotic concentrations for *E. coli* were as follows: 15μg/mL Gentamicin (Gm), 50 μg/mL carbenicillin (Cb), and 50 μg/mL kanamycin (Kan). Antibiotic concentrations for *P. aeruginosa* were as follows: 30 μg/mL Gm, and 250 μg/mL Cb. For inducible constructs, isopropyl-ß-D-1-thiogalactopyranoside (IPTG; GoldBio) was supplemented to cultures and plates at a concentration of 0.1-1 mM and arabinose to a concentration of 0.02-0.2% w/v.

### Construction of *gloR* deletion, *in cis* native-locus complements, *P. aeruginosa* and *E. coli* reporter strains

Generation of the *gloR* deletion mutant in *P. aeruginosa* were carried out as per reference (11) using pEX18Gm and primers *gloR-Up-F*/*R* and *gloR-Down-F*/*R* (**Dataset S3**). Similarly, *in cis* native-locus complements were created using pEX18Gm, but in the MAPO1 *ΔgloR* background. pEX18Gm-gloR-FLAG WT was created using primers *gloR-Up-F*/*gloR-FLAG-R* and *gloR-FLAG-F/gloR-Down-R* (**Dataset S3)**. Site directed mutagenesis was carried out as per reference (REF) using pEX18Gm-gloR-FLAG WT as template and primers *gloR-D97A-F/gloR-D97A-R, gloR-H126A-F/gloR-H126A-R*, *gloR-F187A-F/gloR-F187A-R*, *gloR-C191A-F/gloR-C191A-R, gloR-C191S-F/gloR-C191S-R, gloR-C191D-F/gloR-C191D-R, gloR-H192A-F/gloR-H192A-R, gloR-Y193A-F/gloR-Y193A-R,* and *gloR-Y217A-F/gloR-Y217A-R* (**Dataset S3)**. For plasmid-borne complementation and expression in *E. coli*, the GloR coding sequence, omitting the native stop codon, was then amplified from the MPAO1 chromosome using primer pairs SphI-GloR-F1/BamHI-GloR-R1 and SphI-GloR-F2/BamHI-GloR-R2. The product was then ligated into SphI/BamHI digested pMMB67EH-FLAG (57) to yield pMMB67EH-GloR-FLAG.

### Reporter strain assays

Reporter assays using *P. aeruginosa* strains were carried out using pCG-P*_arqI-gloA2_*-mS by culturing cells to mid-log phase and inducing with increasing concentrations of GO. Assays were conducted as per ref. (11) using a BioTek H1 Synergy multimode plate reader. Reporter assays using the *E. coli* strains harboring pMMB67EH-FLAG and pCG-VmS derivatives were conducted as follows, overnight cultures were diluted 1:100 into fresh LB media containing 100 µM IPTG to induce GloR expression. Cultures were incubated for two hours before GO addition and incubated an additional 3 hours before fluorescence and OD_600_ were measured.

### Western blot analysis

To assess GloR-FLAG protein levels, cells were cultured as described in “Reporter strain assays”. Cells were pelleted then resuspended in 1X SDS loading buffer according to the formula cell culture volume x OD_600_ x 100, to normalize the cell concentration. After boiling, samples centrifuged at 16,000 x g for 10 minutes before being separated by SDS-PAGE and transferred to PVDF membranes (EMD Millipore). Membranes were blocked for 1 hour at RT with WB Blocking Buffer (PBS + 0.05% v/v Tween-20 (PBST-T) + 5% milk). Membranes were probed with Mouse anti-FLAG M2 antibody (1:10,000, F1804, Sigma Aldrich) in WB Blocking Buffer for 1 hr at room temperature, then washed 4 times with PBS-T for 5 minutes each before probing with HRP-conjugated goat anti-mouse (1:10,000, Jackson ImmunoResearch, UK) in WB blocking buffer, again for 1 hr at room temperature. Finally, membranes were washed 4 times with PBS-T for 5 minutes each before developing using Pierce ECL Western Blotting substrate (Thermo Scientific) and imaging on an Azure 400 imaging system (Azure Biosystems, Dublin, CA). Three biological replicates were performed.

### Aldehyde toxicity assays

Aldehyde toxicity tests were carried out as per ref. (11). Briefly, overnight cultures of *P. aeruginosa* MPAO1 were subcultured 1:100 in fresh LB media and incubated at 37 °C with 230 rpm shaking for 2 hours until cells reached mid-logarithmic phase. Cells were then placed on ice for 10 minutes to stop growth before normalizing cell density to an OD_600_ of 1.0 in LB medium. Samples were then serially diluted before spotting 10 μL of each dilution onto LB agar plates containing the appropriate antibiotic and aldehyde.

### RNA-seq

WT and Δ*gloR P. aeruginosa* MPAO1 were cultured to mid-log phase before being treated with 2 mM GO, and cells harvested at 15 minutes post treatment for RNA isolation and sequencing as per ref. (11). For isolation of RNA for sequencing utilized RNAprotect Bacteria Reagent (Qiagen) and RNeasy Mini RNA isolation kit (Qiagen) were used according to the manufacturer’s protocols. Sequencing and downstream analysis was performed by Genewiz (Azenta Life Sciences, Chelmsford, MA). rRNA depletion was implemented and strand-specific RNA sequencing for 5-10 million reads per sample. The experiment was done in duplicate (biological repeats.

### RNA-seq differential expression analysis

Differential expression between P. aeruginosa genotypes and treatment conditions was determined using DESeq2 (Love, Huber & Anders, 2014, Genome Biology). Five pairwise comparisons were performed across the four experimental conditions (wild-type and ΔgloR, each with and without 2 mM glyoxal treatment), each fit independently with its own dispersion estimation and Benjamini-Hochberg multiple-testing correction. Genes were considered differentially expressed at an adjusted p-value (padj) < 0.05 and |log2 fold-change| > 1, a threshold applied consistently across all five comparisons. Raw count matrices from all five comparisons were cross-checked for internal consistency: samples shared across multiple comparison files (*e.g.*, wild-type-untreated samples appearing in more than one pairwise contrast) were confirmed to have identical raw counts in every file, verifying a single, consistent alignment/quantification pipeline underlying all reported comparisons.

### Data visualization

Heatmaps and volcano plots were generated using custom Python scripts (make_heatmap.py, make_volcano.py, and a shared drawing module svg_helpers.py; Python 3.12, pandas, NumPy, and matplotlib for the volcano plots; adjustText for non-overlapping point labels). Heatmap cell colors represent row-standardized z-scores of log2 (normalized count + 1) computed per gene across all samples in all four conditions (n = 2 biological replicates per condition); fold-change and significance for the comparison of interest are reported in an adjacent column using the same padj < 0.05 and |log2FC| > 1 threshold described above. Volcano plots display log2 fold-change against −log10(padj) for all tested genes, with genes passing the significance threshold colored by chromosomal cluster; a full-range panel and a zoomed inset panel (limited to a smaller fold-change window) are shown together so that genes with very large and comparatively modest effect sizes can both be inspected at readable scale within the same figure. Figure generation scripts were developed with the assistance of Claude (Anthropic), an AI coding assistant, through an iterative process in which all statistical thresholds, gene curation decisions (including locus-tag resolution and cluster/operon assignments), and final figure outputs were reviewed by the authors against the underlying DESeq2 output before inclusion in the manuscript.

### Bioinformatic identification of the *glo box cis* acting element

145 bp segments of DNA upstream of the *arqI-gloA2* and *PA1412* promoters was a GloR consensus recognition sequence by inputting into MEME Suite motif search engine using Gapped Local Alignment of Motifs (GLAM2) (58). Search parameters were: one occurrence per sequence, palindromes only, given strand only, and 13-20 bps motif.

### Construction of GloR full-length and GloR_LBD_ expression plasmids

To express full-length GloR we used pE-SUMO (Life Sensors) to construct the pE-SUMO-GloR expression plasmid. All DNA for restriction cloning was generated using a method which does not require restriction enzyme digest of the insert, or the ‘Melt and Reanneal Method’, originally invented in the Weisblum lab (59). The GloR ORF together with an N-terminal Ser-Gly-Gly-Gly linker was PCR amplified from *P. aeruginosa* MPAO1 genomic DNA using two primer pairs to generate BsaI complementary overhangs using primers *BsaI-gloR-F1/BsaI-gloR-R2* and *BsaI-gloR-F2/BsaIgloR-R1* (**Dataset S3**). The PCR products were ligated into BsaI digested pE-SUMO to enable selection of the pE-SUMO-GloR expression plasmid. The GloR ligand-binding domain was expressed using plasmid pSX2-SUMO2 which is derived from pSX2 (Scarab Genomics, Madison, WI). To create pSX2-SUMO2, the 6xHis-SUMO ORF preceded by Met-Val-Lys-Phe-Ser - to increase translation efficiency (60) - was amplified from pE-SUMO using primers *NdeI-KFS-6xHis-SUMO-F1/SacI-KFS-6xHis-SUMO-R2* and *NdeI-KFS-6xHis-SUMO-F2/SacI-KFS-6xHis-SUMO-R1* (**Dataset S3**). The PCR products were ligated into NdeI and SacI digested pSX2 to enable selection of the pSX2-SUMO2 plasmid. To create pSX2-SUMO2-GloR_LBD_, the GloR_LBD_ (residues 87-284) was amplified from *P. aeruginosa* MPAO1 genomic DNA using primers *BsaI-gloR-LBD-F1/BsaI-gloR-LBD-R2* and *BsaI-gloR-LBD-F2/BsaI-gloR-LBD-R1* (**Dataset S3**). The PCR products were ligated into BsaI digest pSX2-SUMO2 to yield the pSX2-SUMO2-GloR_LBD_ expression plasmid for crystallography. The C191S (C110 in the GloR_LBD_ construct) mutant was generated as described in “Construction of *gloR* deletion, *in cis* native-locus complements, *P. aeruginosa* and *E. coli* reporter strains” above using pSX2-SUMO2-GloR_LBD_ as template and primers *gloR-C191S-F/gloR-C191S-R* (**Dataset S3**).

### Expression and purification of full-length GloR

pE-SUMO-GloR was transformed into T7 Express *lysY^I/q^ E*. *coli* (New England Biolabs, Ipswich, MA). Overnight cultures were subcultured 1:100 into fresh LB medium supplemented with 50 µg/mL kanamycin and grown at 37 °C, with 230 rpm shaking until an OD_600_ of ∼0.8 was reached. Protein expression was induced with addition of 1 mM IPTG and the cultures were incubated at 18 °C with 150 rpm shaking overnight before harvesting by centrifuging cultures at 5,000*g* for 10 minutes. Pellets were stored at −80 °C before purification. GloR was first purified by IMAC followed by SEC using an S75 column. Fractions were pooled and stored in the −80 before EMSAs were performed.

### Electrophoretic Mobility Shift Assays (EMSAs)

6-carboxyfluorescein (6-FAM)-labelled oligonucleotides were used for all EMSA experiments. Promoter sequences were PCR amplified using the primer pairs indicated in (**Table S1**). All other EMSAs probes were synthesized as complementary oligonucleotides from IDT and annealed with 10% excess of the unlabeled-reverse complement oligonucleotide to ensure that all labeled strands were ultimately double stranded oligos. EMSAs were performed as in ref. (61) with some modifications. Briefly, purified GloR protein was rapidly thawed from −80 °C in a 37 °C water bath followed by centrifugation at 16,000 x *g* for 1 minute to remove any insoluble material. 10 μL binding reactions were set up as follows: 20 mM HEPES pH 7.2, 50 mM NaCl, 5 mM MgCl_2_, 1 mM CaCl_2_, 0.1 mM EDTA, 12% glycerol, 10 mM DTT, 0.5 μM oligonucleotide, 400 ng Poly(dI-dC). Competing oligonucleotides were added to 10X excess of the labeled oligonucleotide (5 μM). *Listeria monocytogenes* P*_pflBA_* was used as a nonspecific control (**Table S1**). GloR concentrations ranged between 0 and 10 μM. Reactions were allowed to equilibrate on ice for 10 minutes before being resolved on 8% non-denaturing 1X Tris-Glycine EDTA (TGE) gels for 40 minutes at 120 V. Gels were visualized by imaging on an Azure 400 imaging system (Azure Biosystems, Dublin, CA) using the Cy2 filter (ex: 472 nm, em: 513 nm).

### Fluorescence Polarization (FP)

FP was performed as described in ref. (61). A 6-FAM labeled oligonucleotide was utilized as found in **Dataset S1** (5’-ATCAT<u>TCTCTTTAAGTGA</u>GCGATTTGC<u>TGATTATTGATT</u> – 3’). Briefly, 150 μL binding reactions containing 20 mM HEPES pH 7.2, 50 mM NaCl, 5 mM MgCl_2_, 1mM CaCl_2_, 0.1 mM EDTA, 2% glycerol, 10 nM 6-FAM probe, and purified GloR were prepared. Protein concentrations ranged from 0 to 10 μM. Additives were supplemented as indicated. Reactions were allowed to equilibrate for one hour at room temperature before dispensing in triplicate 45 μL aliquots into black 384-well plates (781900; Greiner Bio-One). Polarization was read on a Biotek Synergy H1 multimode plate reader equipped with a green FP filter set (8040561; excitation 485/20 nm, emission 528/2 nm, dichroic mirror 510 nm). Polarization values (in mP) were calculated by the Biotek Gen5 software. To calculate equilibrium dissociation constants, GraphPad Prism software version 10.1.1 for Mac (GraphPad Software) was utilized with a one-site binding model (total binding).

### Expression and purification of GloR_LBD_ for crystallography

pSX2-SUMO2-GloR_LBD_ was transformed into *E*. *coli* Clean Genome (Scarab Genomics, Madison, WI) and grown in LB media supplemented with 50 μg/mL kanamycin to an OD_600_ of 0.6 before protein expression was induced by the addition of 0.25 mM IPTG. Cell growth was continued overnight at 18 °C and 225 rpm shaking before the cells were harvested by centrifugation and stored at −80 °C. Cell pellets were resuspended in Buffer A (50 mM Tris, pH 7.4, 150 mM NaCl, 50 mM imidazole) containing hen egg white lysozyme (10 mg per 10 mL of Buffer A) and 1 mM Phenylmethylsulfonyl fluoride (PMSF), and the cells were lysed by sonication on ice (45% amplitude, 15 seconds on 45 seconds off for 10 cycles). The lysate was centrifuged for 60 minutes at 14,000 rpm and filtered using a 0.2 μm filter (MiniSart, Sartorius). The clarified supernatant was incubated with Ni-NTA resin (HisPur, Thermo Scientific) in a gravity column at room temperature for 1 hour, washed extensively with wash buffer (Buffer A), and the bound protein was eluted with a stepwise gradient of imidazole going from 50 mM to 300 mM final imidazole concentration. Eluted protein purity was assessed by SDS-PAGE and fractions containing pure protein were pooled and dialyzed against 50 mM Tris pH 7.4, 150 mM NaCl, and 1 mM DTT in the presence of ULP-1 protease (1 mg:10 mg ULP1:GloR_LBD_ ratio) overnight at 4 °C to remove the His-SUMO-tag. The cleaved protein sample was incubated with Ni-NTA resin (HisPur, Thermo Scientific) for one hour at room temperature. The flow-through was collected and verified to contain cleaved GloR_LBD_ by SDS-PAGE. The resulting cleaved protein was then concentrated to 10 mg/mL for crystallization trials.

### GloR_LBD_ protein crystallization and structure determination

Crystallization screening was performed with a Mosquito robot (SPT Labtech, Cambridge, MA) and commercially available crystallization screens by the hanging drop vapor diffusion method. In order to provide phase information for the native crystals, selenomethionine-substituted (SeMet) GloR_LBD_ was generated as previously described (62), purified, concentrated to 10 mg/mL, and screened for crystallization. For data collection, crystals of SeMet GloR_LBD_ were grown in PEG/Ion 2, condition 9: 4% Tacsimate pH 7.0, 12% PEG3350, and cryo cooled directly from the crystallization condition. A multi-wavelength anomalous diffraction (MAD) dataset was collected at SSRL from cryo-cooled crystals.

The MAD data were collected on a single crystal at energies: 0.9796 Å (E1), 0.9794 Å (E2) and 0.9116 Å (E3) (**Table S2**). The datasets were processed in XDS (63) and an initial solution was determined in Crank2 to 2.5 Å resolution (63). The initial model was refined manually in Coot and in Phenix.refine (64, 65). Following refinement, a GloR_LBD_ dimer was used to search for 2 copies of a GloR_LBD_ dimer in the E1 dataset in Phaser (66). The molecular replacement (MR) solution was improved by molecular replacement single-wavelength anomalous diffraction (MR-SAD) in AutoSol (67). Subsequently, the solution was run through AutoBuild (68). The resulting model was then refined as described above in Coot and Phenix.refine to a final R_work_/R_free_ of 21.8/25.7 % (**Table S2**). Refinement in Phenix.refine utilized Translation-Libration-Screw-rotation (TLS) parameters.

Diffraction-quality native GloR_LBD_ crystals were grown in 4% Tacsimate pH 8.0. 20% PEG3350 with a reservoir to protein drop ratio of 1:1. Data were collected at SSRL, and processed in XDS to 2.6 Å resolution (63) (**Table S2**). Using a subunit from the SeMet GloR_LBD_ structure, we were able to solve the phases by MR in phaser (66). Interestingly, these crystals were in a different space group to the SeMet protein crystals, with only two subunits in the ASU. The structure solution was refined as described above in Phenix.refine and Coot, and refined to a final R_work_/R_free_ of 23.3/30.0 % (**Table S2**).

In order to study glyoxal interactions with GloR, GloR_LBD_ at 10 mg/mL was incubated with 4 mM glyoxal and then used for crystallization trays. Diffraction-quality crystals were grown in 0.1 M sodium nitrate, 0.1 M Bis-Tris Propane pH 8.5, and 22.5% PEG 3350, and cryocooled in the crystallization condition for collection at SSRL. These crystals were processed, solved by MR, and refined as per the native crystals described above. The resulting structure has resolution to 2.1 Å RMSD and refined to a final R_work_/R_free_ of 21.9/24.8 % (**Table S2**).

### Mass spectrometry (MS) of purified GloR_LBD_

Purified GloR_LBD_ WT and C110S (C191 in full-length GloR) mutant proteins (1.8 µM) were incubated with either MGO or GO (3.6 µM) for 30 minutes at 25 °C in 50 mM Tris-HCl (pH 8.0) and 150 mM NaCl prior to analysis. In a separate reaction, WT GloR_LBD_ (1.8 µM) was incubated with GO (3.6 µM) for 30 minutes, followed by the addition of DTT (3.6 µM) and an additional 20-minute incubation before analysis. Samples were analyzed by MS using LC-MS/MS (ACQUITY UPLC H-class system, Xevo G2-XS QTof, Waters) adapted from a previous methodology (69). To remove buffer salts, samples were subjected to reverse-phase chromatography at 45 °C using a C4 column (Protein BEH C4 Column, 300 Å, 1.7 μm, 2.1 mm X 50 mm, Waters) and a 5-minute gradient of Buffer C (0.1% formic acid in water), Buffer D (100% acetonitrile from 0% Buffer D to 90% Buffer D with a flow rate of 0.3 mL/min). The Xevo Z-spray source was run in positive ion mode with a capillary voltage of 300 V, and a cone voltage of 40 V (NaCsI calibration, Leu-enkephalin lock-mass). Nitrogen was used as the desolvation gas at 350 °C and a flow rate of 800 L/hour and data acquisition was done at alternating collision energy with low energy at 0 V and high collision energy ramp from 15-45 V. Data was acquired in continuum with 0.5 second scans across a mass range of 400-4000 Da. Total average mass spectra were reconstructed from the charge state ion series using the MaxEnt1 algorithm from Waters MassLynx software V4.1 SCN949 according to the manufacturer’s instructions. To obtain the ion series described, the major peak of the chromatogram was selected for integration before further analysis.

### Rat lung infections

Rat lung infections were carried out using an intra-tracheal administration procedure. Solutions containing bacterial inoculums were directly instilled into the trachea. Recipient rats were anesthetized with isofluorane. After the anesthesia takes effect (as judged by a toe pinch), rats were placed on an angled stage, secured by a band around the abdomen and a wire held across the incisors. The tongue is withdrawn from the mouth and a rigid luer stub inserted into the trachea. The syringe delivers 300 μL of solution. Rats are held in place for 2-5 seconds to allow complete inhalation of the dose, and then placed back into the cage for recovery from the anesthetic. Animals undergoing this procedure typically return to normal activity and feeding patterns within 90 minutes of instillation. *P. aeruginosa*, MPAO1, or mutations derived thereof, were grown in 30 mL of LB broth for 12 hours. Bacteria are encapsulated in agarose beads, essential for the induction of prolonged infection. A 1 mL suspension of the agarose beads is then collected, washed in PBS and resuspended in 1 mL of PBS. Lethal dose at 50% (LD50) experiments have determined the appropriate inoculum, which is 3 x 10^6^ CFUs. 8 rats per strain were calculated to give sufficient statistical power for these studies. CFUs were counted after rats are euthanized by harvesting lungs, homogenizing them, and plating cells on Pseudomonas isolation agar (PSA) plates.

### Statistical analysis

Statistical analyses were calculated using GraphPad Prism or with the assistance of Claude (Anthropic). Specific tests are described in the figure legends.

## Data availability

Mass spectrometry data has been deposited in MassIVE under MSV000101505_reviewer password GloR. Atomic structural data has been deposited in the RCSB Protein Data Bank under accession numbers 9NLI (native GloR_LBD_ structure, unmodified), 9NRQ (GO modified GloR_LBD_ structure), and 9NLM (SeMet unmodified GloR_LBD_ structure). Other raw data are provided either as supplementary information/data files or in the Source Data file.

## Code availability

Scripts used to generate the heatmaps and volcano plots in this manuscript will be deposited in a public GitHub repository under the MIT license upon acceptance for publication.

## Author Contributions

C.J.C. and A.T.U. conceived the project. C.J.C., A.T.U., D.G.G, B.C., S.E.B., P.P., C.W.G. analyzed the data. C.J.C., D.G.G., B.C., R.D., J.M., P.P., and J.D.K. performed the experiments. C.W.G. and A.T.U. wrote the manuscript.

## Supporting information

Corcoran et al. Suppl Figures and Tables

Dataset S1 GloR RNA-seq data

Dataset 2 GloR homologs

Dataset 3 Strains, plasmids and primers

## Acknowledgements

This work was funded by grants NIH grants R01AI135060 and R01GM141230 to A.T.U.

