## Supplementary material for "A glyoxal sensing *Pseudomonas aeruginosa* transcription factor enables lung infection": Corcoran et al. Suppl Figures and Tables

### Figure S1

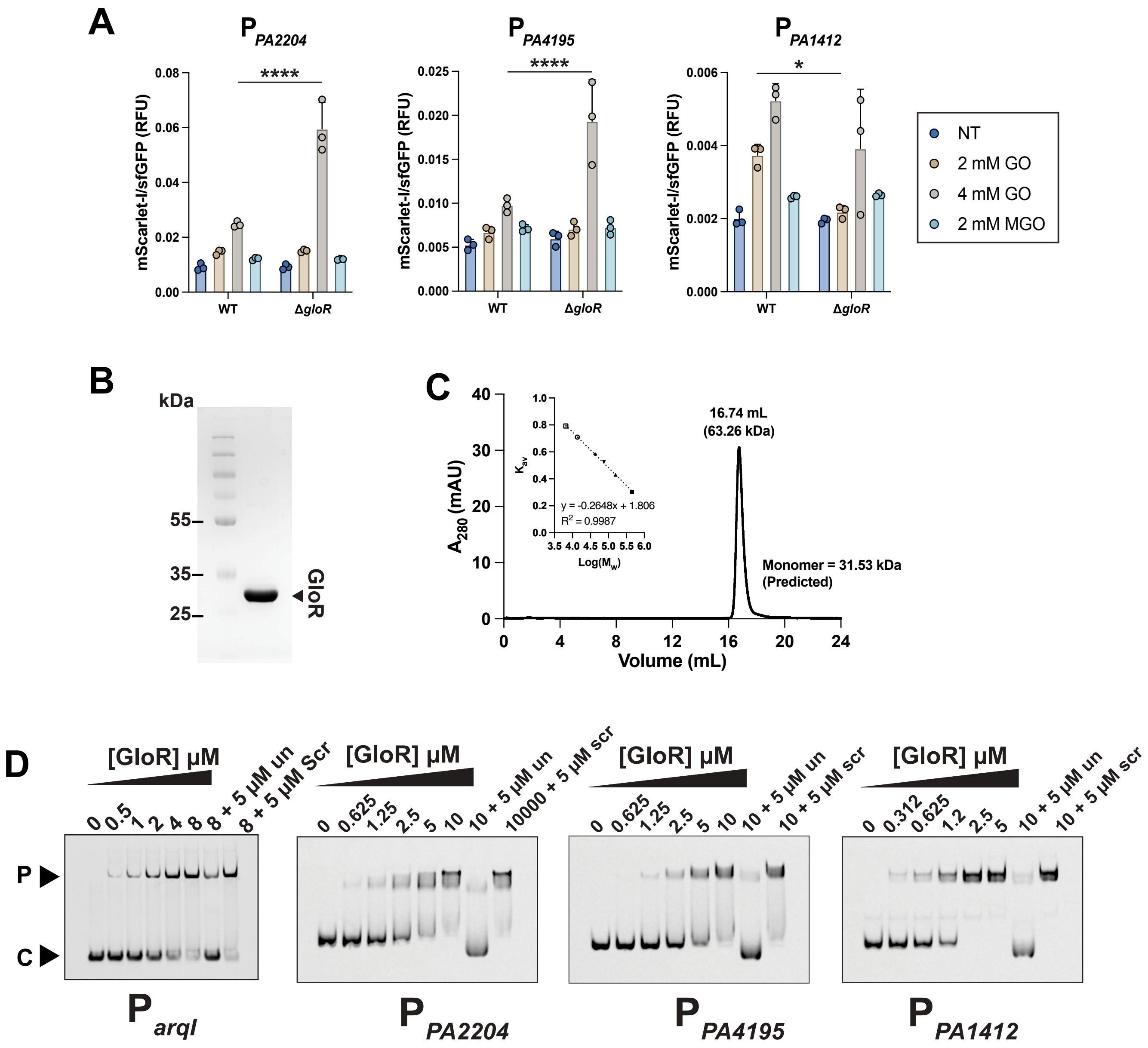

A
P PA2204
P PA4195
P PA1412
****
*
mScarlet-I/sfGFP (RFU)
WT
ΔgloR
NT
2 mM GO
4 mM GO
2 mM MGO
B
kDa
55
35
25
GloR
C
16.74 mL
(63.26 kDa)
Monomer = 31.53 kDa
(Predicted)
A280 (mAU)
Volume (mL)
Kav
y = -0.2648x + 1.806
R2 = 0.9987
Log(Mw)
D
[GloR] µM
0 0.5 1 2 4 8 8 + 5 µM un 8 + 5 µM Scr
0 0.625 1.25 2.5 5 10 10 + 5 µM un 10000 + 5 µM scr
0 0.625 1.25 2.5 5 10 10 + 5 µM un 10 + 5 µM scr
0 0.312 0.625 1.2 2.5 5 10 + 5 µM un 10 + 5 µM scr
P
C
P arqI
P PA2204
P PA4195
P PA1412

### Figure S2

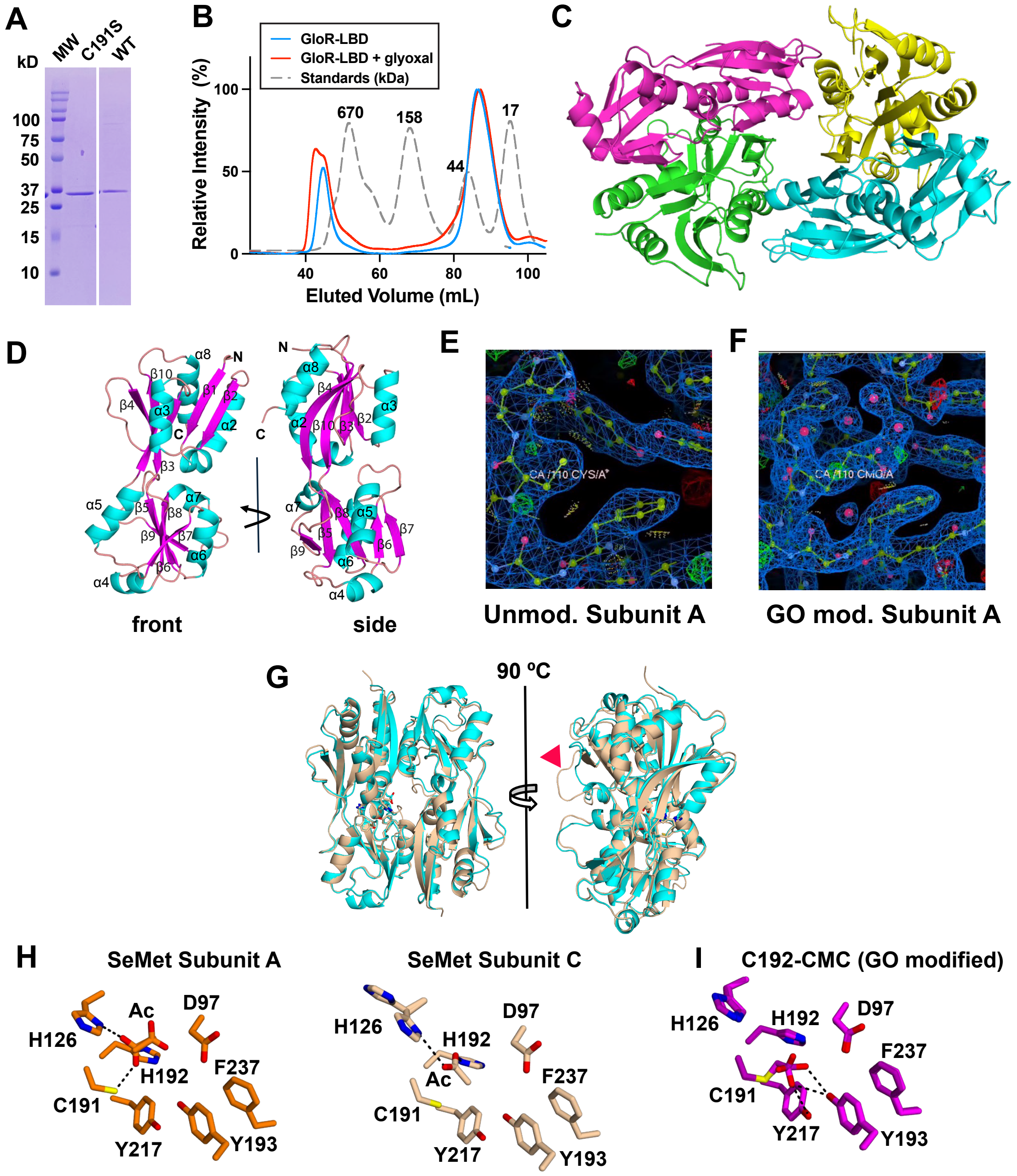
A
kD
MW
C191S
WT
100
75
50
37
25
15
10
B
GloR-LBD
GloR-LBD + glyoxal
Standards (kDa)
Relative Intensity (%)
100
50
0
670
158
44
17
40
60
80
100
Eluted Volume (mL)
C
D
front
side
E
Unmod. Subunit A
F
GO mod. Subunit A
G
90 ºC
H
SeMet Subunit A
H126
Ac
D97
H192
F237
C191
Y217
Y193
SeMet Subunit C
H126
H192
D97
Ac
F237
C191
Y217
Y193
I
C192-CMC (GO modified)
H126
H192
D97
F237
C191
Y217
Y193

### Figure S3

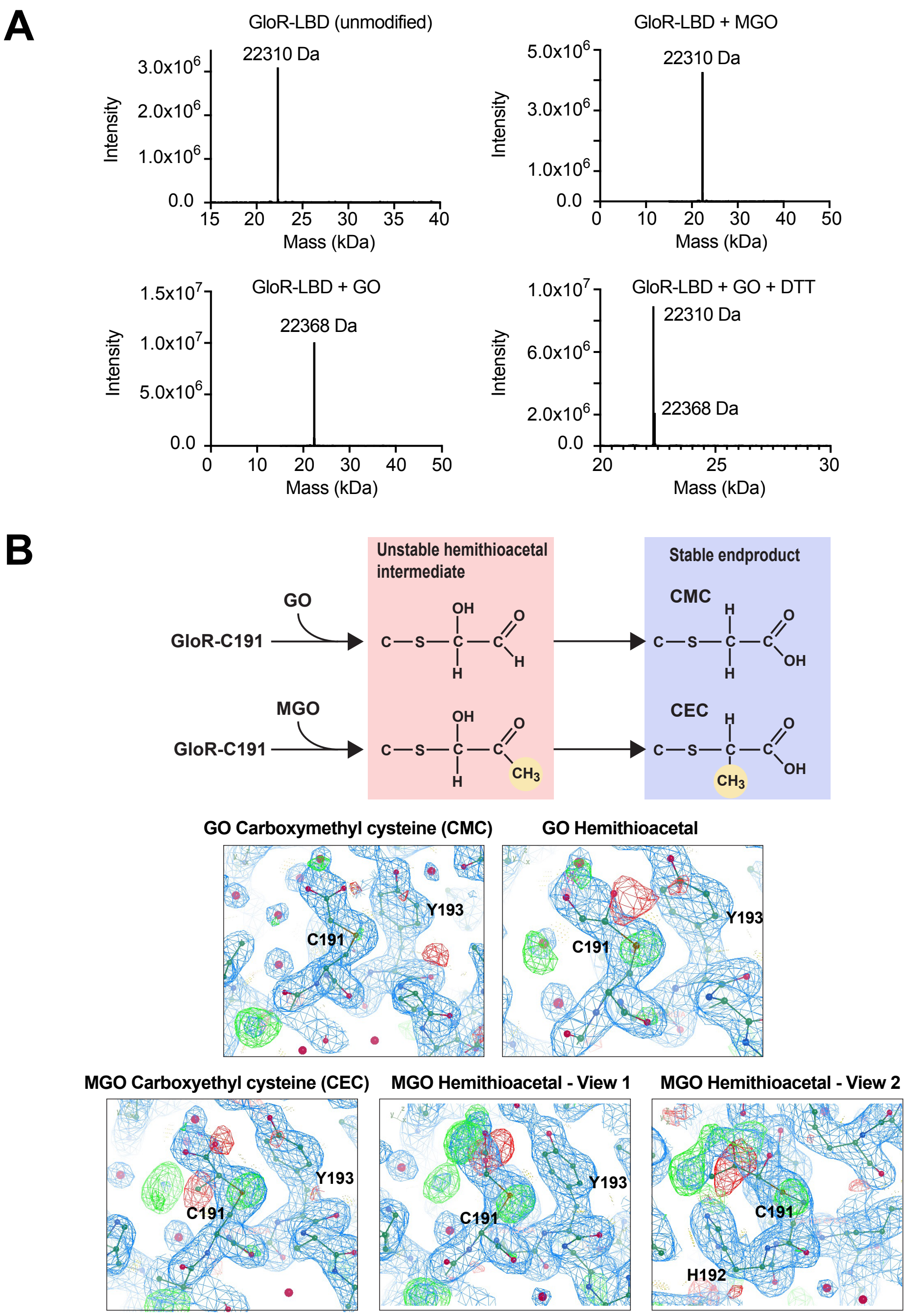
A
GloR-LBD (unmodified)
22310 Da
Intensity
Mass (kDa)
GloR-LBD + MGO
22310 Da
GloR-LBD + GO
22368 Da
GloR-LBD + GO + DTT
22310 Da
22368 Da
B
Unstable hemithioacetal intermediate
Stable endproduct
GO
MGO
GloR-C191
CMC
CEC
GO Carboxymethyl cysteine (CMC)
GO Hemithioacetal
MGO Carboxyethyl cysteine (CEC)
MGO Hemithioacetal - View 1
MGO Hemithioacetal - View 2
C191
Y193
H192

### Figure S4

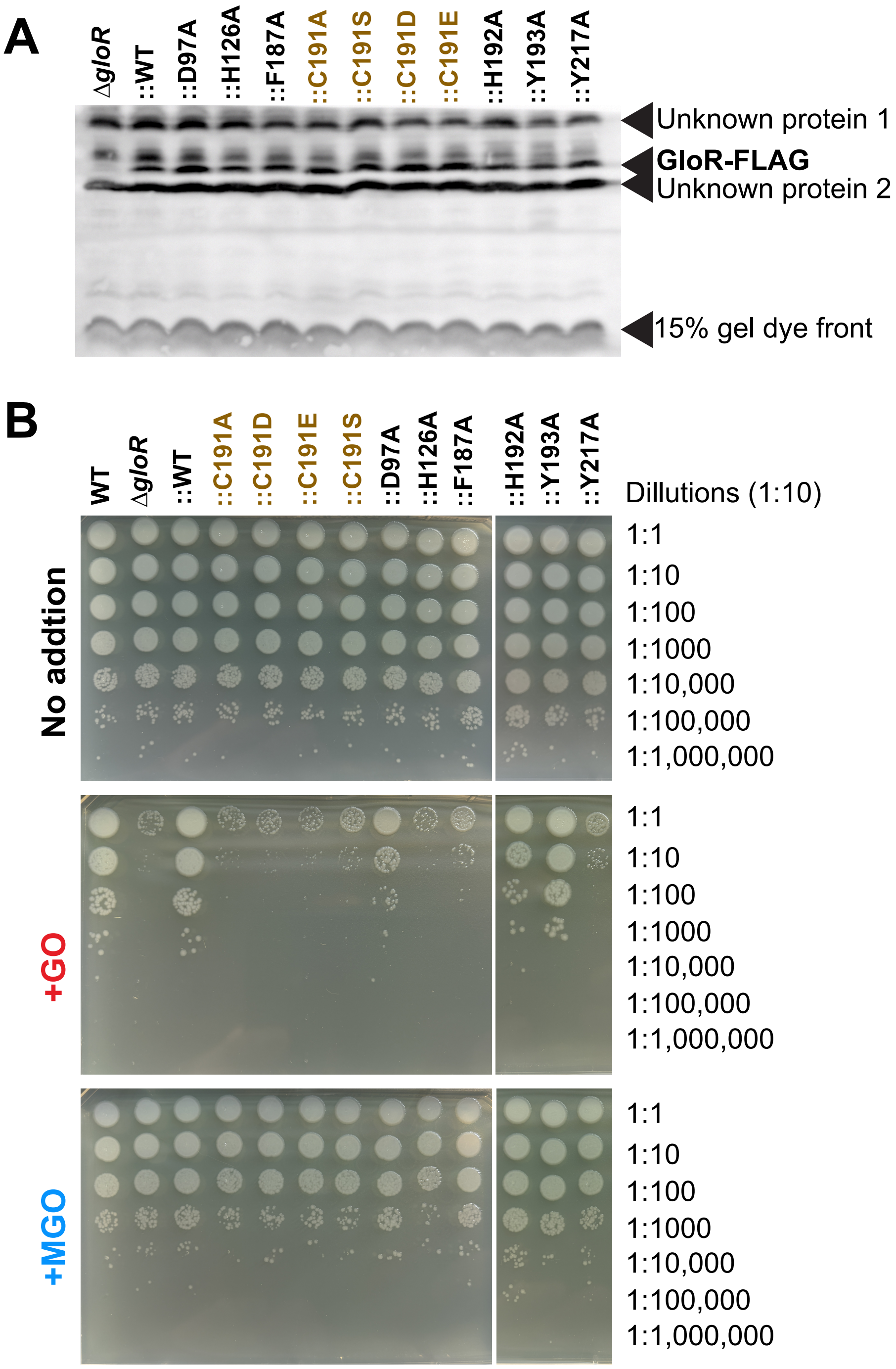
A
ΔgloR
::WT
::D97A
::H126A
::F187A
::C191A
::C191S
::C191D
::C191E
::H192A
::Y193A
::Y217A
Unknown protein 1
GloR-FLAG
Unknown protein 2
15% gel dye front
B
WT
ΔgloR
::WT
::C191A
::C191D
::C191E
::C191S
::D97A
::H126A
::F187A
::H192A
::Y193A
::Y217A
Dillutions (1:10)
No addtion
+GO
+MGO
1:1
1:10
1:100
1:1000
1:10,000
1:100,000
1:1,000,000

**Figure S5**

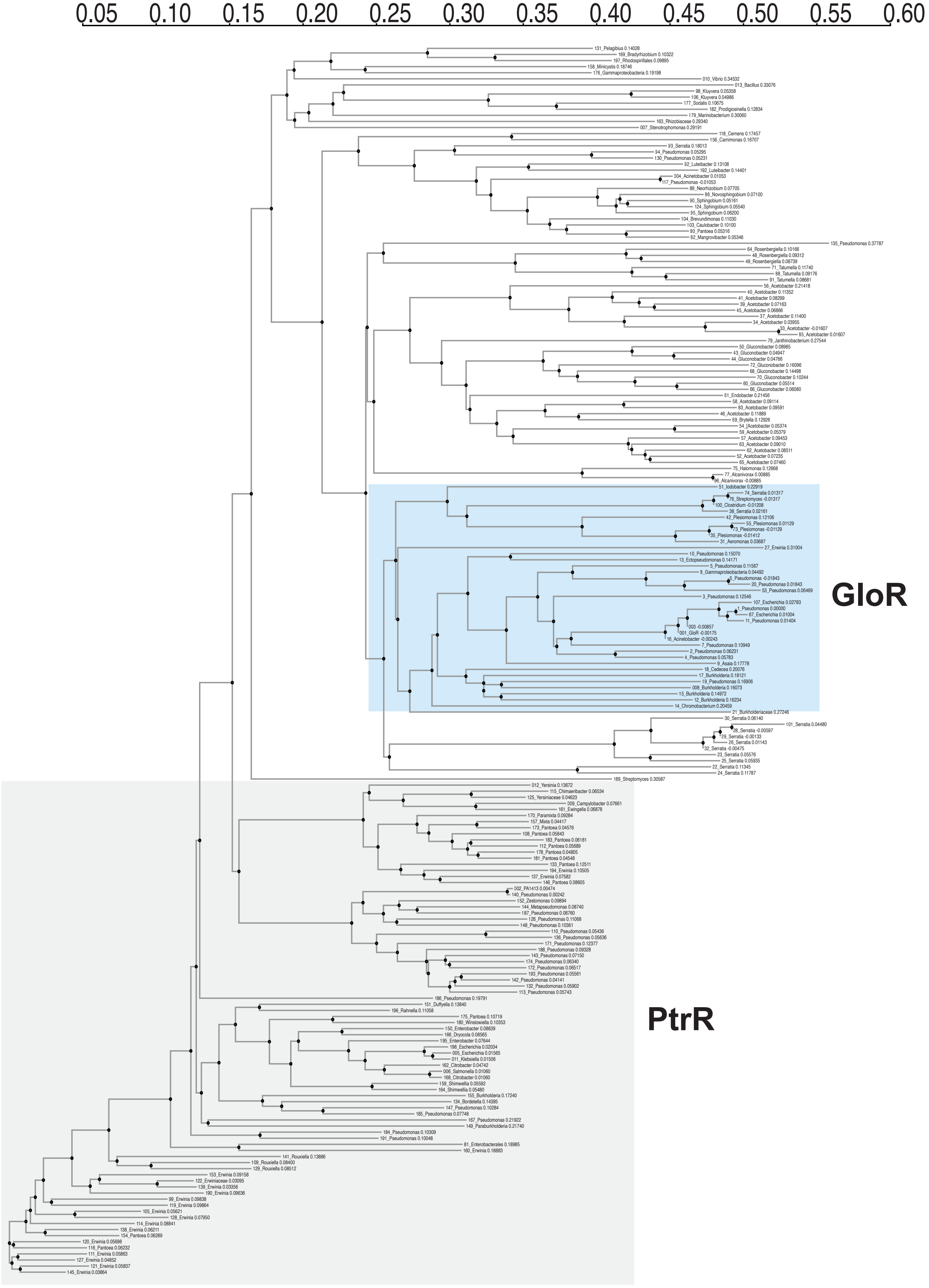

#### Figure S6

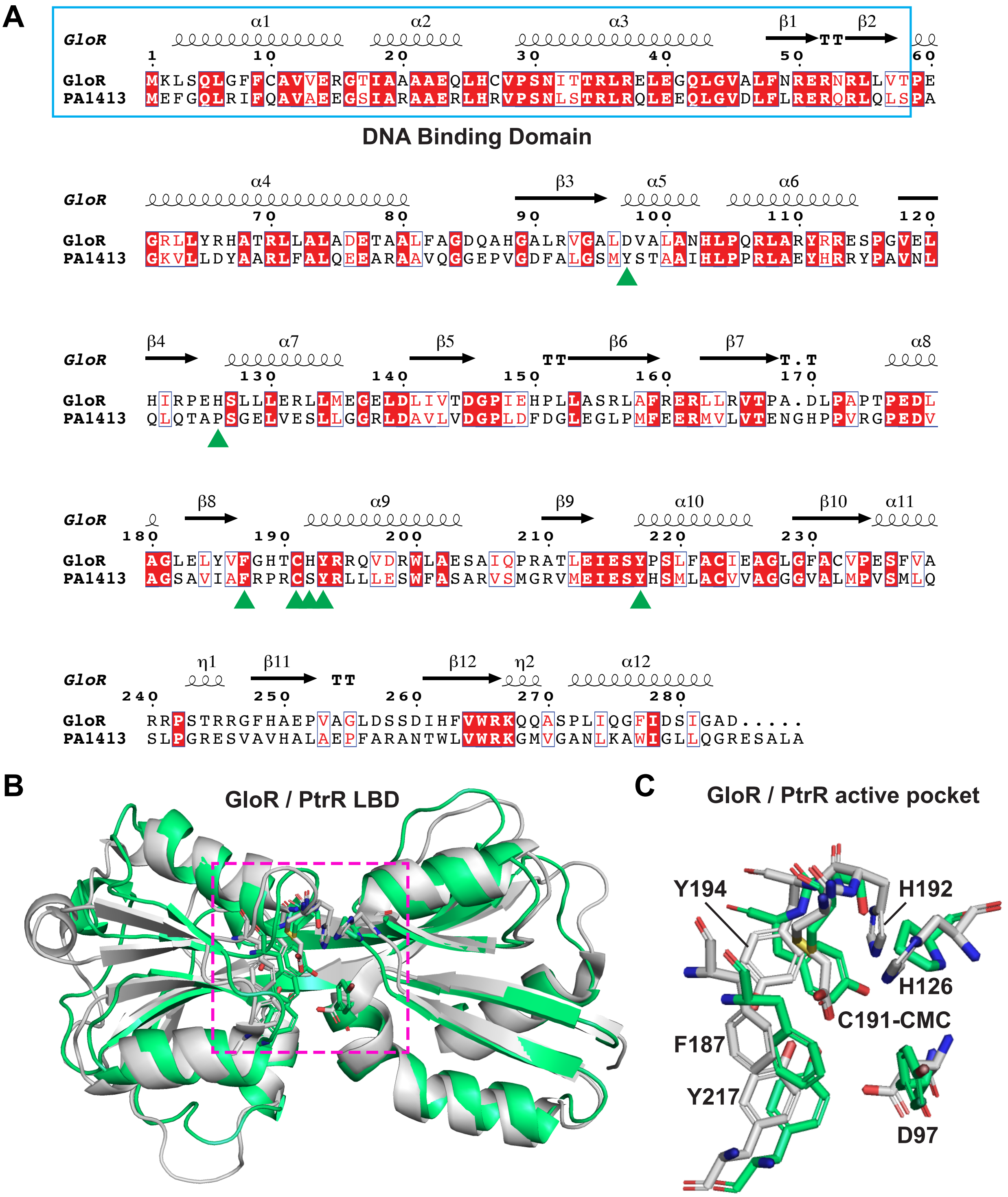

A
GloR
GloR MKLSQLGFFCAVVERGTIAAAAEQLHCVPSNITTRLRELEGQLGVALFNRERNRLLVTPE
PA1413 MEFGQLRIFQAVAEEGSIARAAERLHRVPSNLSTRLRQLEEQLGVDLFLRERQRLQLSPA
DNA Binding Domain
GloR GRLLYRHATRLLALADETAALFAGDQAHGALRVGALDVALANHLPQRLARYRRESPGVEL
PA1413 GKVLLDYAARLFALQEEARAAVQGGEPVGDFALGSMYSTAAIHLPPRLAEYHRRYPAVNL
GloR HIRPEHSLLLERLLMEGELDLIVTDGPIEHPLLASRLAFRERLLRVTPA.DLPAPTPEDL
PA1413 QLQTAPSGELVESLLGGRLDAVLVDGPLDFDGLEGLPMFEERMVLVTENGHPPVRGPEDV
GloR AGLELYVFGHTCHYRRQVDRWLAESAIQPRATLEIESYPSLFACIEAGLGFACVPESFVA
PA1413 AGSAVIAFRPRCSYRLLLESWFASARVSMGRVMEIESYHSMLACVVAGGGVALMPVSMLQ
GloR RRPSTRRGFHAEPVAGLDSSDIHFVWRKQQASPLIQGFIDSIGAD.....
PA1413 SLPGRESVAVHALAEPFARANTWLVWRKGMVGANLKAWIGLLQGRESALA
B
GloR / PtrR LBD
C
GloR / PtrR active pocket
Y194
H192
H126
C191-CMC
F187
Y217
D97

**Table S1.** Significantly differentially regulated WT vs. Δ*gloR* transcripts

| **Δ*gloR* 2 mM GO vs WT 2 mM GO (15-minute incubation)** | | | |
|---|---|---|---|
| **Gene** | **$Log_2$ Fold Change** | **Padj** | **Annotation / Prediction** |
| ***tauD1*** | 1.687 | 3.93E-05 | Taurine dioxygenase |
| ***PA2203*** | 1.659 | 7.11E-24 | Possible arginine/ornithine transporter |
| ***ssuC*** | 1.430 | 2.64E-06 | Aliphatic sulfonate permease |
| ***ssuD*** | 1.405 | 0.01540846 | Alkanesulfonate monooxygenase |
| ***PA4192*** | 1.327 | 1.58E-08 | Polar amino acid transport (ArtP) |
| ***PA3445*** | 1.320 | 0.00012566 | Hypothetical |
| ***toxR*** | 1.122 | 1.93E-22 | Transcriptional regulator |
| ***PA4191*** | 1.111 | 6.77E-10 | Isopenicillin-N synthase/ascorbate iron reductase |
| ***aguB*** | 1.065 | 2.77E-31 | N-carbamoylputrescine amidohydrolase |
| ***tauA*** | 1.049 | 0.00181819 | Periplasmic taurine-binding protein |
| ***cysW*** | 1.037 | 1.26E-18 | Sulfate transport |
| ***PA2088*** | 1.027 | 0.00041108 | Hypothetical |
| ***tonB2*** | 1.017 | 0.00263937 | TonB transporter |
| ***PA4195*** | 1.012 | 1.93E-11 | Polar amino acid transporter |
| ***PA2204*** | 1.000 | 0.0101233 | Possible arginine/ornithine transporter |
| ***PA2481*** | -1.135 | 4.18E-30 | Possible cytochrome c |
| ***PA1412*** | -1.376 | 7.64E-11 | MFS general transporter |
| ***gloR*** | -2.721 | 1.45E-117 | LysR-type transcription factor |
| ***PA0712*** | -3.663 | 0 | Possible toxin/antitoxin pair |
| ***PA0714*** | -4.681 | 2.24E-211 | Hypothetical |
| ***PA0713*** | -4.749 | 0 | Hypothetical |
| ***PA0711*** | -7.407 | 0 | Possible toxin/antitoxin pair |
| ***gloA2*** | -7.785 | 0 | Lactoylglutathione lyase |
| ***arqI*** | -7.834 | 0 | Regulates PQS |

[1]Differentially regulated genes with a $log_2$Fold Change greater than 2 or less than -2 are displayed.
[2]A Padj equal to zero is rounded down due to the actual value being below the displayable value.

**Table S2.** Data collection, phasing and refinement statistics for GloR datasets.

| | Native | Glyoxal | MAD dataset | | |
|---|---|---|---|---|---|
| **Data collection** | | | | | |
| Space group | P $2_1$ $2_1$ 2 | P $2_1$ $2_1$ 2 | P $6_1$ | | |
| Cell dimensions | | | | | |
| *a*, *b*, *c* (Å) | 99.9, 117.3, 34.3 | 100.5, 119.1, 34.2 | 149.1, 149.1, 75.8 | | |
| (°) | 90, 90, 90 | 90, 90, 90 | 90, 90, 120 | | |
| | | | *Peak* | *Inflection* | *Remote* |
| Wavelength | 1.0 | 1.0 | 0.97961 | 0.97944 | 0.91162 |
| Resolution (Å)[a] | 39.1-2.6 (2.72-2.6) | 38.4-2.09 (2.15-2.09) | 37.4-2.5 (2.6-2.5) | 37.4-2.5 (2.6-2.5) | 37.4-2.5 (2.6-2.5) |
| $R_{merge}$[b] | 0.122 (1.132) | 0.146 (1.284) | 0.078 (1.239) | 0.202 (4.063) | 0.168 (3.320) |
| *I* / □*I* | 8.2 (1.8) | 6.2 (1.5) | 16 (2.1) | 17.4 (15.7) | 10.1 (1.7) |
| Completeness (%) | 99.9 (100) | 98.9 (91.5) | 99.5 (97.6) | 99.0 (92.4) | 99.4 (96.0) |
| Redundancy | 6.9 (7.2) | 9.2 (8.8) | 12.9 (11.7) | 17.4 (15.7) | 17.5 (17.2) |
| **Refinement** | | | | | |
| Resolution (Å) | 38.0-2.6 (2.7-2.6) | 36.9-2.09 (2.17-2.09) | 37.4-2.5 (2.56-2.50) | | |
| No. reflections | 12998 (1393) | 24735 (1202) | 33367 (1301) | | |
| $R_{work}$ / $R_{free}$[c] | 23.3/30.0 (38.6/51.1) | 21.9/24.8 (32.1/36.0) | 21.8/25.7 (35.9/41.4) | | |
| Number of subunits | 2 | 2 | 4 | | |
| Number of Residues | 399 | 395 | 788 | | |
| No. atoms | 3120 | 3242 | 6014 | | |
| Protein | 3120 | 3110 | 5973 | | |
| Ligand/ion | 0 | 0 | 34 | | |
| Water | 0 | 132 | 7 | | |
| *B*-factors | 64.4 | 44.6 | 91.1 | | |
| Protein | 64.4 | 44.6 | 91.0 | | |
| Ligand/ion | N/A | N/A | 94.2 | | |
| Water | N/A | 46.3 | 114.2 | | |
| Ramachandran Favored | 96.2 | 97.4 | 96.1 | | |
| Ramachandran Outliers | 0 | 0 | 0 | | |
| R.m.s deviations | | | | | |
| Bond lengths(Å) | 0.008 | 0.009 | 0.009 | | |
| Bond angles (°) | 1.06 | 1.05 | 1.14 | | |
| PDB Deposition ID | 9NLI | 9NRQ | 9NLM | | |

[a]. Values within parentheses refer to the highest shell.

[b]. $R_{merge} = \sum\sum|I_{hkl} — I_{hkl}(j)| / \sum I_{hkl}$, where $I_{hkl}(j)$ is observed intensity and $I_{hkl}$ is the final average value of intensity. $R_{pim} = \sum\{1/[N_{hkl}-1]\}^{1/2} \times \sum|I_{hkl}(j) — I_{hkl}| / \sum\sum I_{hkl}(j)$. $R_{meas} = \sum\{N_{hkl}/[N_{hkl}-1]\}^{1/2} \times \sum|I_{hkl}(j) — I_{hkl}| / \sum\sum I_{hkl}(j)$. $CC = \sum (x-(x))(y-(y)) / [\sum(x-(x))^2 \sum(y-(y))^2]^{1/2}$.

#### Supplementary Figure Legends

**Figure S1. Additional EMSAs and reporter assays to assess GloR regulation.** (**A**) WT MPAO1 and Δ*gloR* MPAO1 strains harboring the dual reporter system to analyze the activity of GloR regulated promoters were cultured and either not treated (NT) or treated with 2 mM GO, 4 mM GO or 4 mM MGO and reporter activity read after 3 hours incubation. Statistical analysis: *, $p < 0.05$; ****, $p < 0.0001$ by Two-way ANOVA and Šídák's multiple comparisons test. (**B**) SDS-PAGE of purified full-length GloR protein. (**C**) Size exclusion chromatography (SEC) chromatogram showing that GloR migrates as a predicted dimer. 50 µl of GloR at 100 µM (monomer) concentration was injected onto a Superose 6 increase10/300 GL column equilibrated with buffer (50 mM Tris-Cl pH 8.0, 500 mM NaCl, 1 mM TCEP). Protein was resolved at a flow rate of 0.4 mL/minute. (**D**) EMSAs of promoter regions of *arqI-gloA2*, *PA2204*, *PA4195*, and *PA1412*. Increasing amounts of purified GloR were added to a fixed amount (0.5 µM) of fluorescently labeled dsDNA probe. Un, unlabeled probe (competitive). Scr, scrambled *glo box* binding site sequence to prevent GloR from binding (non-competitive).

**Figure S2. Purification and structure insights into the $\mathbf{GloR_{LBD}}$.** (**A**) SDS-PAGE gel of WT $GloR_{LBD}$ and the $GloR_{LBD}$ C191S mutant purified protein. (**B**) SEC analyses of $GloR_{LBD}$ with GO (red spectrum) and without GO (blue spectrum) using a HiLoad 16/600 column. Standards are shown in grey (dotted line) with the peaks labeled in Daltons. (**C**) Cartoon representation of $GloR_{LBD}$ tetramer. The four monomer subunits are shown as magenta, cyan, green, and yellow. (**D**) 180° rotation of the $GloR_{LBD}$ monomer with labeled α-helices (cyan), β-strands (magenta) and loop regions (wheat). (**E**) Omit map of subunit A of the unmodified structure and (**F**) modified structure with C191-CMC. Here C191 is labeled as C110. (**G**) Front and side views of the SeMet (wheat) and unmodified (cyan) structures superimposed showing the loop region which differs between them (highlighted by the red arrow). (**H**) Ligand binding pocket residues from SeMet subunits A (orange) and C (wheat) showing the acetate hydrogen bonding versus (**I**) the same pocket in the C191-CMC modified structure (magenta). Dotted lines indicate hydrogen bond interactions.

**Figure S3**. **Mass spectrometry and structural modeling to interrogate C191 adducts.** (**A**) MS of $GloR_{LBD}$. Purified WT $GloR_{LBD}$ was subjected to either GO, MGO, GO for 10 minutes followed by DTT treatment, or no DTT addition control, and subjected to LC-MS. Calculated masses are shown above peaks. The theoretical mass for $GloR_{LBD}$ is 22,309.16 Daltons (Da), and $GloR_{LBD}$ modified with GO to form C191-CMC observed a mass shift of 56.84 Da (calculated). (**B**) Chemical formation (above) and atomic modeling (below) on C191 of GO CMC and MGO CEC adducts versus their corresponding precursory, unstable hemithioacetal PTMs. Hemithioacetal GO and MGO cysteine modification adducts were modelled into the density of the stable GO CMC adduct (carboxymethyl) cysteine at position 191 from the crystal structure, and then refined in phenix.refine (71) to generate an electron density map. The resulting model and map are shown for the stable CMC and CEC adducts, and more unstable hemithioacetal GO and MGO versions. Images were generated in Coot (72). Carbons are shown in green, the 1 Å $2F_o$-$F_c$ map in blue, and the 3 Å $F_o$-$F_c$ is shown in green (positive density) and red (negative density).

**Figure S4. Western blot of GloR variant expression and representative plates for aldehyde killing assays.** (**A**) Western blot showing GloR-FLAG levels of protein in knock-in *P. aeruginosa* strains. *P. aeruginosa* expressing GloR-FLAG variants from the native chromosome location were cultured in LB to mid-log phase, pelleted and run as whole cell lysates on an SDS-PAGE before membrane transfer and probing with an M2 anti-FLAG monoclonal antibody. GloR-FLAG is indicated by arrows and the C191 variants labeled in brown. (**B**) Photos of plates from aldehyde killing assays where 1:10 dilutions of *P. aeruginosa* were spotted to determine CFUs. These plates are representative of 3 biological repeats.

**Figure S5. Phylogenetic tree of GloR related sequences.** The GloR sequence from *P. aeruginosa* PAO1 strain was used in a NCBI Blast search to identify homologs. Sequences were uploaded into Clustal Omega (33) to generate an alignment, which was curated for sequences that harbored a C191 equivalent residue before a phylogenetic tree was finally generated (also using Clustal Omega). GloR and PtrR homologs are colored blue and grey, respectively, based on their amino acid adjacent to the conserved C191 (Histidine for GloR and Serine or other residues for PtrR). Branch lengths indicate divergence, with shorter branches signifying greater similarity, typically representing substitutions per site (*e.g.*, 0.1 = 10% difference). Numbers on branches represent distance, with lower scores indicating closer relationships. Nodes (junctions) represent common ancestors. Groups of sequences branching from the same node are more closely related. Branch ends are labeled with the corresponding number to Dataset S2, the distance number and the genus of the species. Full species names and NCBI numbers which are unique to the sequences are given in Dataset S2.

**Figure S6. GloR and PtrR primary and tertiary alignments.** (**A**) *P. aeruginosa* GloR and PtrR alignment. The GloR sequence from *P. aeruginosa* PAO1 (PA0708) was aligned to the PtrR homolog from PAO1 (PA1413) using Clustal Omega and alignment generated using ESPript version 3 (73). GloR secondary structure information based on the deposited crystal structure (PDB code 9NLI) is given above the alignment. Key pocket residues mentioned in the text are indicated with the green arrows. A blue box highlights the DNA binding domain. (**B**) Superimposition of $GloR_{LBD}$ modified structure (grey) and a $PrtR_{LBD}$ model generated from the

*P. aeruginosa* sequence (locus number PA1413) with AlphaFold (green) (35). The pink dotted line box indicates the ligand binding pocket(s). (**C**) Closeup of ligand binding pocket(s) from the $GloR_{LBD}$/$PtrR_{LBD}$ overlay in *B*. CMC, carboxymethyl cysteine.
