## Supplementary material for "A glyoxal sensing *Pseudomonas aeruginosa* transcription factor enables lung infection": Dataset 2 GloR homologs

>001_GloR
MKLSQLGFFCAVVERGTIAAAAEQLHCVPSNITTRLRELEGQLGVALFNRERNRLLVTP
EGRLLYRHATRLLALADETAALFAGDQAHGALRVGALDVALANHLPQRLARYRRESPG
VELHIRPEHSLLLERLLMEGELDLIVTDGPIEHPLLASRLAFRERLLRVTPADLPAPTPED
LAGLELYVFGHTCHYRRQVDRWLAESAIQPRATLEIESYPSLFACIEAGLGFACVPESF
VARRPSTRRGFHAEPVAGLDSSDIHFVWRKQQASPLIQGFIDSIGAD
>002_PA1413
MEFGQLRIFQAVAEEGSIARAAERLHRVPSNLSTRLRQLEEQLGVDLFLRERQRLQLS
PAGKVLLDYAARLFALQEEARAAVQGGEPVGDFALGSMYSTAAIHLPPRLAEYHRRYP
AVNLQLQTAPSGELVESLLGGRLDAVLVDGPLDFDGLEGLPMFEERMVLVTENGHPP
VRGPEDVAGSAVIAFRPRCSYRLLLESWFASARVSMGRVMEIESYHSMLACVVAGGG
VALMPVSMLQSLPGRESVAVHALAEPFARANTWLVWRKGMVGANLKAWIGLLQGRE
SALA
>004_Acinetobacter baumannii 2
MQLSQLRAFCAVAHTGSINAAAKALNRVPSSLSVRIRQLEEDLGCQLFLREHQRLRLS
PDGRRLLEHARRILDLSDSTRAMMRNEESGGRLVVGALDVVLVAFMPCLIGRFRQRH
RAIELDIRSEASEALVQQVSDGVLDLALSDGPVRSQTLESSLAFVDEMVLVTELDHPPIT
SPRDLRCAELYGFRHDCSFRFRMDRWLEEADCLQQLPVLEIESYHTMLACVSAGMGA
AWVLRSVLATLPGNHQVRAHPLRDYGTTEIHFIWHSGHLTPNAQRLMA
VHQDALVERRD
>005_Escherichia coli EPEC
MDLTQLEMFNAVAEAGSITQAAAIVHRVPSNLTTRLRQLETELGVDLFIRENQRLRLSP
AGHNFLRYSQQILALVDEARSVVAGDEPQGLFSLGSLESTAAVRIPATLAEFNHRYPKI
QFSLSTGPSGTMLEGVLEGKLNAAFIDGPINHTAIDGIPVYREELMIVTPQGHAPVIRAS
QVNGSNIYAFRANCSYRRHFESWFHADGAAPGTIHEMESYHGMLACVVAGAGIALIPR
SMLESMPGHHQVEAWPLAEQWRWLTTWLVWRRGAKTRPLEAFIQLLDVSDSAKQSY
Q
>006_Salmonella enterica
MDLTQLEMFNAVAETGSITQAAAKVHRVPSNLTTRIRQLEADLGVDLFIRENQRLRLSP
SGHNFLRYSQRILALVEEARMVVAGDEPQGLFSLGSLESTAAVRIPATLAEYNQRYPKI
QFALATGPSGTMLDGVLDGTLNAAFVDGPVAHPGLEGIPVYREEMMIVTPHGHPPIQR
ASDVNGCSIYAFRANCSYRRHFESWFHADRAMPGTIHEMESYHGMLACVIAGAGIALI
PRSMLESMPGHHQVNAWPLSENWRWLNTWLVWRRGAMTRQLEAFIEVLDPQLSQD
TD
>007_Stenotrophomonas maltophilia
MDDTSLDVFRTVAEELSITRAAQRLGRAPSNVTTRVQQLEAELGAELFVRSGKRLSLS
PQGERFLGYAQRLLALTEEARQSLQPDALSGLLRIGSMESTAASRLPQPLARLHARWP
QVRLEVSTGPSAQLLERLRTQAIDCALLALPPSQEAADLAPSGIELQPVFNEQLELLLPA
GHPPVSGPADLQVTTLAAFAAGCSYRAVAEHWLAAAGHRLDVQVVGSYHAMLACVA
AGHSACLMPLSVRALSPALDLRSVPLFDLPTQLAWRSGYLTPALDALRSSLLQP
>008_Burkholderia multivorans
MKLSHLGFFCAIVEHGTVAAAADKLNCVPSNVTIRIRELEEQLGVALFSRERNRLFVTP
EGRLLYDKARTLLGLASEAEQLFTGERPVGTLKIGALDVVLSNHLPPRLPAYRSRMPNV
ELHLHPGHSLVLERKLVDGEFDLIVSDGPISHPLLSSSLAFRENLKLITPKTVRTLSPKSL

AAFEFYVFGKDCHYRQTVDNWLASTGLAPRALLEIESYPAMFACVAAGHGIACVPDSF
LATYRSAYPVNAHALPATGSAEIHFIWRKYQVSTLVTDFIEIISGKAAA
>009_Campylobacter jejuni
MDLTQLRMFCCVAETGSLARAAEQLHRVPSNLTTRLRQLELELGTDLFIREKQRIRLSA
MGHNFLNYAERILALSDEAMSITHAGEPAGNFALGSMESTAATRLPSLLAAYHQKFPQ
VALSLVTATSGETIDSVRAGRLAAALVDGPIDFDDINGCISFRERMVVITPPGESPQEVP
AGRSGHTVFAFRPSCSYRQRLQGWIKHMSIPVANTMEIQSYHSMMACVASGAGIAMV
PRSVLEQLPGHERVSIHEIPDEFADTATWLIWRRDAFSPNVRALKELIIEQNDGVIPLLK
HPLAGEHDPALHPTY
>010_Vibrio cholerae
MNKIFLFSELMDSSDLKIFVAVVEQGGIVNASAFLHKVPSAISTRIKQLESNLDTQLFTRE
HRKLFLTAEGKILYEYALDILSLTERAERHVKHRSPGGKFKLGAMDSMASSRLPEPLSN
LYIRYPSIQLELITGISQFLSDAIKNNSLDATFIADFPTDDRFERVTVFKEELVVVAPSNHP
PINSPKDIECDTLLVFKEGCSYRSRLFDWYRDFQVQPTRVATMSSYPVILASVASGMGI
GIVPLSLIEDFKYSNSIRIHKIAEIEPITTDLIWKKGHYSLNIESLLEVLNVSSFQE
>011_Klebsiella pneumoniae YneJ
MDLTQLEMFNAVAEAGSITQAAAKVHRVPSNLTTRLRQLETELGVDLFIRENQRLRLSP
AGHNFLRYSQQILALVDEARSVVAGDEPQGLFSLGSLESTAAVRIPATLAEFNRRYPKI
QFSLSTGPSGTMLDGVLEGKLNAAFIDGPINHTAIDGIPVYREELMIVTPQGYAPVTRAS
QVNGSNIYAFRANCSYRRHFESWFHADGAAPGTIHEMESYHGMLACVIAGAGIALIPR
SMLESMPGHHQVEAWPLAEQWRWLTTWLVWRRGAKTRPLEAFIQLLDVPDSARQGY
Q
>012_Yersinia pestis
MDLTQLRMFCCVAETGSVARAAEQMHRVPSNLTTRLRQLELELGADLFIREKQRLRLS
PMGHNFLCYANRILALSEEAMSITHAGEPAGNFPVGSMESTAATRLPNLLAAYHQRYP
KVSLSLITGTSGEIIEQVRAGTLATALVDGPVQHDELHGCRSFDEQLVIISCLERPPILQA
RDAVDETLFAFRPSCSYRLRLESWFRQVGVLPGHIMEIQSYHAMLACVASGAGLALIPL
SVLTLLPGHERVQVHPLPPDIAHTATWLIWRKDAFSPNVRALKELIIENVEIP
>013_Bacillus cereus
MDIKDLTVFKKVAEEGSISKAAKSLSYVQSNVTMKIKQLEDELGVPLFYRNGKGVTLNS
NGEILLTYTRKILDLTEQSIRLVQSNQSKPLGTLKIGSTNSTVAVRLPPLLKNYCEKYPEV
ELVLETHKSTELTQLVLERKLEGAFIVEDVHHPDLTSFLFCQEELTIISYQPLHTLQELDK
VNMLAFGNGCHYRNRVDKWLKEAGIYPKRVLEFGTIEAIIGCVKAGMGIAVMVKSILKD
HEQSLTMTDLPEKYSKVPTYFIMRKDVLFSAALQRFVELIKGKTM
>1_Pseudomonas aeruginosa_GloR
MKLSQLGFFCAVVERGTIAAAAEQLHCVMGASLPACANWKQQLGVALFNRERNRLLV
TPEGRLLYRHATRLLALADEPDH
CSPATRRTAHYGCLTLDVALANHLPQRLARYRRESPGVELHIRPEHSLLLERLLMEGEL
DLIVTDGPIEHPLLASRLAFR
ERLLRVTPANLPAPTPEDLASLELYVFGHTCHYRRQVDRWLAESGIQPRATLEIESYPS
LFACIEAGLGFACVPESFVAR
MPSTRRGFHAEPVAGLDSSDIHFVWRKQQASPLIQGFIDSIGAD
>2_Pseudomonas sp. MF6751

MKLSQLGFFCAVVERGTVAAAAAQLHCVPSNVTTRLRELEDQLGVALFNREKNRLLVT
PEGRLLYRNAKQLLALADKTRS
LFTDDDVRGVLRVGALDVALDNHLPQRLARYRGEAPLVELHIRPEHSLMLERLLMEGE
LDLILTDGPIVHPLLVSRLAFR
ERLVRVTPINLTTPTIDDLARLELYVFGHTCHYRRQIDHWLEMSGIEPQVTLEMESYPSI
FACIEAGLGFACVPESVVAQ
VSLTKKKVHAEAVASLDSSDIHFVWRKNQASPLIQSFIDSISS
>3_Pseudomonas sp. CMR5c
MKLSQLSFFCAVVEHGTIAAAAAQLHCVPSNITTRLRELEGQLDVVLFNREKNRLLVTP
EGRLLYRNAKQILAMTADTQS
LFSSEKPHGVLRVGALDVALANHLPPRLSRYRRENPTVELHIRPEHSMLLERLLMEGE
MDLILTDGPIEHPLLASHLAFR
ERLLLVTPTKLSAPSTEELEHLELYVFGHTCHYRRQVDHWLETCDIKPRVALEIESYPNI
FACIEAGLGFACVPESVVNQ
FSLTHKGFYAQHIQGLGSSDIHFVWRKQQASPLIQGFIESI
>4_Pseudomonas fluorescens
MKLSQLGFFCAVVELGTVAAAAAQLHCVPSNITTRLRELEEQLGVVLFNREKNRLLVTP
EGRLLYRNAKQLLVMAADTRS
LFTGDEVQGVLRVGALDVALANHLPQRLARYRREAPFVELHIRPEHSLMLERLLLEGEL
DLILSDGPIVHPLLVSRLAFC
ERLVRVTPINLTTPTADDLAKLELYVFGHTCHYRRQVDRWLEESGIQPQATLEIESYPSI
FACIEAGLGFACVPESFVAQ
ISLTQKKVHAEAVAGLDSSDIHFVWRKHQASPLIHRFIESIGSS
>5_Pseudomonas sp. S31
MKLSHLGYFCAVVEQGTIAAAATQLHCVPSNITTRLRELENQLGVALFNREKNRLMTTP
EGRLLYRNAKKMVDLAAETKS
LFSSECNHGVLRVGALDVALANHLPTGLVRYRREAPGVELHIRPEHSLLLERLLMDGEL
DLILTDGPVEHPLLESRLAFR
EKLLAVTPNKLNTITPENLSNLELYVFGQTCHYRQQVDRWLAISGILPRAILEIESYPSIF
ACINEGLGFACIPESYVLK
FSQQKYDFKSELAHGLDSSDIYFVWRKNQQSPLIEDFIRIITS
>6_Pseudomonas abietaniphila
MKLSQLGFFCAVVDHGTIAAAAASLHCVPSNITARLRELEIQLGVVLFNRENNRLLVTPE
GRLVYRKAKQLIDLAAETRS
LFSNDSMQGVLRVGALDVALANHLPERLVRYRLEAPGVELHIRPEHSLFLERLLMDGE
LDLILTDGPIQHPLLDSRLAFR
ERLIRVVPKALSSPTPQDLSGLELYVFGRTCHYRQQVDNWLESSGIQPRAILEIESYPSI
FACIAQGIGFACIPESYVDR
FASQAHTFQADQVPSLDSSDIYFVWRKNQQSPLIRNFIEVIDS
>7_Pseudomonas fluorescens
MVEHGTIAAAAAQLHCVPSNITTRLRELEEQLGVVLFDREKNRLLVTPEGRLLYRHARQ
LLDMANHTRALFAAEEAHGVL
RVGALDVALANHLPHRLARYRRHAPGVELHIRPEHSLLLERLLMDGELDLILTDGPIEHP
LLTGQLAFRERLLRVTPWHL

KAPTADDLAALELYVHGHTCHYRRLVDSWLAHSGITPRMTLEIESYPAIFACIEQGLGF
ACVPESFLEAIAPAERRFHAE
AVAELASSDIHFVWRKQQASPLIQGFIDSVAA
>8_Gammaproteobacteria
MKLSQLGFFCAVVECGTIAAAAVQLHCVPSNITTRLRELENQLGVVLFNREKNRLLVTP
EGRLVYRNAKLLVDLAAETRS
LFSTDSINGVLRVGALDVALANHLPQRLVRYRLEAPGVELHIRPEHSLRLERLLIDGELD
LILTDGPIEHPLLESRLAFR
ERLLRVVPKAFSSPTLENLAGLELYVFGRTCNYRLQVDNWLESNGIHPRAILEIESYPSI
FACIAEGLGFACVPESYVDR
YASQADTFQADYVPGLDSSDIYFVWRKSQQSPLIKNFIEII
>9_Asaia siamensis
MKLSQLAFFCAVVEQGTVAAAAAQLNCVSSNITTRVRELEDLLGVSLFSREKNRLLVTP
EGRLFYQKAKQVLAMAQEARA
LFKDDAERGVIRVGALDVALTNHLPERIARYRANTPHVELHIRPEPSLMLEQLLMDGEL
DLILTDGPIIHPLLVSRLAFR
ERLVLVTPKALSAPTVDDLSRMELYVFGQTCHYRRQVDGYLETSKIKPRSVMEIESFSS
IFACIEAGVGFSCLPESVVEQ
AVTRHGKIHFERTDSLESSDIYYVWRKNYTSPLIHRLIESL
>10_Pseudomonas sp. 5P_3.1_Bac2
MKLSQLKFFCAVVEQGTVAAAAEQLHCVPSNITTRLKELEEGLATTLFNREKNRLLITP
QGRLFYQHAKQLLSLAQQTEG
LFRDDSQPQGVLRIGALDVALASHLPQHIVSYRSAYPSVELHIRPEHSFVLERLLMEGE
LDLIVTDGPIEHPLLASSPAF
HEQLLLITPPGVHADSSKLAELELYVFGKSCYYRHQVDQWIDQRIQPRITLEIESYPSILA
CVAAGLGFACIPESIFKAS
GLAIQQVQAQVVEELSASDIYFVWRKQQHSPLIEKFIE
>11_Pseudomonas sp. AF1
QRLARYRRESPGVELHIRPEHSLLLERLLMEGELDLIVTDGPIEHPLLASRLAFRERLLR
VTPADLPAPTPEDLAGLELY
VFGHTCHYRRQVDRWLAESAIQPRATLEIESYPSLFACIEAGLGFACVPESFVARRPST
RRGFHAEPVAGLDSSDIHFVW
RKQQASPLIQGFIDSIGAD
>12_Burkholderia alba
MKLSQLGFFCAVVEQRTIAAAAEQLHCVPSNITTRLRELEALLGVTLFSRERNRLLVTP
EGRLLYDRAKALLGAADDTRQ
LFAGERRAGALRLGALDVALGSHLPKRIAEYRRAMPGVELHVHPGHSFALERRLIDGE
LDLIVSDGPIEHPLLASSLAFR
ETLKLVTPRALRTPSPARLAEFEMYVFGTDCYYRQQVDEWLARTGAAPRALLEVESYP
VMFACVAAGHGIACAPESFIRA
SVDAAGLRAHALDAIGESHLYFIWRKQQRSTLIAEFIDIVAA
>13_Ectopseudomonas mendocina
MKLSQLNFFCTVVEHGTIAAAAERLHCVPSNITTRIKELEEELGASLFSREKNRLLITPQ
GRLLYSRATQLLQLAQETKE

LFVHDAQPQGVLRMGALDVALTQHLPPHIVRYRRTHPKVELHIRPGHSFQLERLLVEG
ELDLIVTDGPIEHPLLASSLAF
NERLVLITPKEVRETADELNTLELYVFGKNCYYRHQVDQWISERIKPRAMLEIESYPTIL
ACVSAGLGFACIPESILNSA
PDVHAHLNSRALDELPSSDIYYVWRKQHTSPLVENFM
>14_Chromobacterium haemolyticum
MKLSQLNFFCAVVEQGSIAAAAEQLHRVPSNLTARLQELEAELGVKLFSRENRRLLTTP
EGHLLYRHARGLLQQADEVRG
LFAGAHARGLLRVGALDGALATHLPARIARYRRQHAEVELHLRAGHSLQLERQLQDGE
LDLILSDGPVEHPLLTSRLAFR
EKLMLIAPLDIDAASAQALAGLELYTFGAACHYRRLVDQWLQASGVRPRAVLEIESYPAI
FACVAAGAGFSCVPESLCRP
QDGYRGFALPDLAISDIHFIWRKQQESALVDGFVQSVVAN
>15_Burkholderia ambifaria
MKLSQLSFFCAVVEHGTIASAAEHLHCVPSNITTRLRELEEQLGIALFSRERNRLFVTPE
GRLLYDRAKTLLALADDANR
LFSNTLPTGLLNVGALDAVLSDHLPARIAEYRRQRPGVELNLYPGHSFVLENRLSEGEL
DVIISDGPIDHPLLESKFAFR
ETLTLISPESVRTPSPARLAAFELYVFGKDCYYRQLVDHWLAETGIAPRATLEIESYPIM
FACVAAGQGIACVPESLALP
LKRRYAVRSHALDGIGPSDTYFVWRKHQASALIADFVSIVSTD
>16_Acinetobacter baumannii
PSNLTTRLRELEGQLGVALFNRERNRLLVTPEGRLLYRHARRLLALADETAALFAGDQ
AHGALRVGALDVALANHLPQRL
ARYRRESPGVELHIRPEHSLLLERLLMEGELDLIVTDGPIEHPLLASRLAFRERLLRVTP
ADLPAPTPEDLAGLELYVFG
HTCHYRRQVDRWLA
>17_Burkholderia stabilis
MKLSQIKFFCAVVEHGTVAAAAKALNCVPSNITIRVRELESQLNVALFIRERNRLFVTPE
GRLLYEKAGTLLALASETQQ
LFEGNQPSGSLNVGALDAVLSNHLPPHIANYRRARPRVRLNIHPGHSFALERRLVDGE
LDVIVSDGPIDHPLLASSIAFR
ESLMLITPKAIRRVSAEWYAEHELYVFGTSCYYRQLVDAWISRNQVTPRAVLEIESYPI
MFACVAAGHGFAFVPASFFAT
YQRGYSVRAHQLDDIGSADTYFVWRKHQASALVPDFIDNCLPD
>18_Cedecea sp.
MKLSQLRFFCAVVEQGTIAAAADMWYCVPSNITMRIHELERQLGVALFSREKNRLMVT
PEGRLLYRKAKNMLEMATEAEQ
LFYNVSKTGLLNVGALDVALSAHLPQRIADYLLREPGVELNLHCGQSFTLERQLIEGEL
DLIISDGPVEHPLLTSALAFC
ECLKLITPMEVSNLSAMQLARYDLYVFGEKCYYRHQVNRWLALSGISPRAVLEIESYPV
MFACVASGIGCACVPESFIAD
TPTFRVHSLDLIGSCNMYFVWRKHLKSPVVVEFI
>19_Pseudomonas chlororaphis

MKLSQLSFFCAVVEHGTIAAAAEQLHCVPSNITIRIRELERLLGTKLFSRERNRLYVTPE
GRLLYTKAKVLLSQALEIQH
LFEGSARAGMLNVGALEGVLNYQLPPFIARYRSMMPNIELNLHSGHTFSLERMLTDGE
LDLIISDGPIEHPLLSSSLAFR
ESLRLITPQDVMAPSAKNISALEFYVFGKECNYRQVVDNWLAGLGIAPRAILEIESYPVM
FACVAQGHGMACVPESSLSM
YREAYSVNSHRLDNVGAVDTYFIWRKYQVSRLTTDFIDIVSG
>20_Pseudomonas graminis
QGVLRVGALDVALANHLPERLVRYRLEAPGVELHIRPEHSLFLERLLMDGELDLILTDG
PIQHPLLDSRLAFRERLIRVV
PKALSSPTPQDLSGLELYVFGRTCHYRQQVDNWLESSGIQPRAILEIESYPSIFACIAQG
IGFACIPESYVDRFASQAHT
FQADQVPSLDSSDIYFVWRKNQQSPLIRNFIEVIDS
>21_Burkholderiaceae bacterium
LKLTPLKFFCAVVEGGTVAHAAERLFCVPSNVTMRLRELEEQMNVTLFNRERNRLYVT
PEGRLLYEQAREILNRVEEVSG
LFTNRVRRGELRIGALDVALDAYLPQMIAHYRRQQSRVMTHLTQAPSLELERQLMGGE
LDLIFTDGPITHPLLKSCRAFD
VQLLCVTPITTTNLSPQALVGLELYVFRKTCVYRKMADLWLEKNQVKPISVIEIESYPVT
LACVEAGLGFAFVPETYMAN
TFSFYRNLTAHIVLPKTPVYAVWRRGDRSAVMRDFVSNVTA
>22_Serratia sp. NPDC087055
MKLHQLSLFCAIVENGTIAAAAKKLNCVPSNITIRLKQLEDQLDTKLFNREKNRLLITPQG
RILYRNAKDLLERSMDIKK
LFNKNAEVQGFFFIGALDGVLSAHMPSYIVNHRIKHPKVELNIRSDNSLSLERDLLDGAL
DLIICDGPIEHPLLSSSLAF
KESLVLIVPPSLPETITLSLLETLDLYTFRKSCSYRHQIENWLSSNHVTPNMVLEMESYD
VITACVSAGLGFSLVPESIL
EKQSLCKHDVQTIKIPELNTNDVFFVWRKSDDKSDLISDFI
>23_Serratia proteamaculans
MKLSQLRFFCAVVEYKTIAAAARELHCVPSNVTLRIKELEESLGGELFFRDKNRLYVTP
KGRLFYQQAVEILALTEQSQQ
LFAGRQQQGLLNFGALDFSLVSHLPGRIAQLRRSQPQMQVNVLSRDSLVLERMLIDSE
LDLAVTDGPIEHPLLAGRAAFD
ERLVLLMPASVPQLNAATLAQLEFYTFSRECSFRLKIDHWLASRQFKPRMTLEMESYT
AMVACVKAGCGVACIPGSLLPL
ILPTSELKVMEMGEEGVSNLYFVWRRHQLSDELQAAIE
>24_Serratia marcescens
MKLHQLSLFCAIVENGTIAAAAKKLNCVPSNITIRLKQLEEQLGAKLFNREKNRLLITPQG
RILYRSAKDLLLRAVNIEH
LFNRNTAANGVFLVGALEGVLSSHMPSYIVNHRMKYPNIELHIRSDNSLNLERELFEGA
LDLIITDGPIEHPLLSSTLAF
RERLMLIVPPFLPATIDLGFLETLDLYTFRKSCSYRHLIEYLLSSKKIRPNMILEMESYDVI
TACVSAGLGFSLVPESIL

EKCFPTQSKIQAMEIPELERNDLFFVWRKSDDASMLVSDFVSGV
>25_Serratia liquefaciens
MKLSQLRFFCAVVEYKTIAAAARELHCVPSNVTLRIKELEDSLGGELFFRDKNRLYVTP
KGRQFYQQAVDILTLADRSQQ
LFAGRQPQGLLNFGALDFSLISHLPSRIAHLRRSQPHMQVNVLSRDSLVLERMLIDSEI
DLALTDGPIEHPLLASSAAFD
ERLVLLMPASVPQLDAATLAQLEFYTFSRECSFRLKIDRWLASRQLKPRMTLEMESYT
AMAACVKAGCGVACVPGSLLPM
ILPEQALKVVGMGAEGISNLYFVWRRHQLSDELQTALD
>26_Serratia ureilytica
MKLSQLKFFCTVVEHKTIAAAARELHCVPSNVTLRLRELEESLGGELFFRDKNRLYVNP
KGRLFYQQARDIVAQAERSKQ
LFAGEQQHGLLNLGALDFSLVSHLPARIARLRRLQPHLHINVLSRDSLVLERMLIDSDLD
LAITDGPIEHPLLASQKAFD
ERLVLLMPADAGQPDAATLAPLEFYTFSRECSFRLKVDHWLASRGLKPRMTLEMESY
AAMAACVKDGCGVACVPGSLLPL
ILPAPGLKVVEMGEEGVSDLYFVWRRHQLSEELQTILE
>27_Erwinia psidii
LKISQLVYFCAVVEEKSIALAALRCHCVAANITIRIKELERLLDAELFYREKNRIFVTPEGR
LLYENAKKMIDLHDRTVA
IFSADAEKGTLNIGALDAALSNQLPGVIAMCREKMPGVAFNVRTGHSFDLESALLDGSL
DIIISDGPVQHPLMTSILAFK
ESLMLVTSSNVSGISQASFNERELYVFGDNCFYRNQVDNWLTANNLKPRIMMEFESYP
VMFSCIKKDLGCAFIPESFRDH
IKNAGDFTFHTGLIEETSDLYFIWRKHSRSKLTANFINHV
>28_Serratia sp. IR-2025
MKLSQLKFFCTVVEHKTIAAAARELHCVPSNVTLRLRELEESLGGELFFRDKNRLYVNP
KGRLFYQQARDIVAQAERSKQ
LFAGEHQHGLLNLGALDFSLVSHLPARIARLRRLQPHLHINVLSRDSLVLERMLIDSDLD
LAITDGPIEHPLLASQKAFD
ERLVLLMPADAGEPDAATLAPLEFYTFSRECSFRLKVDHWLASRGLKPRMTLEMESYA
AMAACVQAGCGVACVPGSLLPL
ILPAPGLKVVEMGEEGVSDLYFVWRRHQLSDELQTIL
>29_Serratia marcescens
MKLSQLKFFCTVVECKTIAAAARELHCVPSNVTLRLRELEESLGGELFFRDKNRLYVNP
KGRLFYQQARDIVAQAERSKQ
LFAGEHQHGLLNLGALDFSLVSHLPARIARLRRLQPHLHINVLSRDSLVLERMLIDSDLD
LAITDGPIEHPLLASQKAFD
ERLVLLMPADAATLAPLEFYTFSRECSFRLKVDHWLASRGLKPRMTLEMESYAAMAAC
VQAGCGVACVPGSLLPLIMPAP
GLKVVEMGEEGLSDLYFVWRRHQLSDELQTIL
>30_Serratia sp. root2
MKLSQLKFFCALVEHKTIAAAARELHCVPSNVTLRIRELETTLGGDLFFRDKNRLYVNP
RGRLFYQQARDIVMQAERSRQ

LFVGEQPQGLLNLGALDFSLVSHLPARIAQLRRLQPQLQVNVLSRDSLVLERMLIDSEL
DLAITDGPIEHPLLASQKAFD
ERLVLLMPAAVAQPDAATLSELEFYTFSRECSFRLKVDHWLASRNLKPRMTLEMESYT
AMAACVKAGCGVACIPGSLLPL
ILPAPELKVVEMGEEGVSDLYFVWRRHQLSAELQATLEILA
>31_Aeromonas bestiarum
MNNSQLAAFCKVVESESVTAAATQLFCVPSNITKKIKELESEFEVMLFTRDRNRLLLTP
EGRTFYQKAKQALQLEEQGRR
LFTRDQLRGILHIGALDIALSHYLPAQIAAFRTQHRDIQLNITPGYSLDLEYQLLKGERDII
FSDGPNQHPLLQSRLAFA
EELVLIGAESEPALLSQRDLYVFTRQCHYRHLIDRWISQHAVIPRAVLEIESYPVIFACVR
AGLGFALVPKSLAESEHLA
YSAQPDRIKCDIYAIWRSAHASPLLQDFIHSITA
>32_Serratia marcescens
MKLSQLKFFCTVVEHKTIAAAARELHCVPSNVTLRLRELEESLGGELFFRDKNRLYVNP
KGRLFYQQARDIVTQAERSKQ
LFAGEHQHGLLNLGALDFSLVSHLPARIARLRRLQPHLHINVLSRDSLVLERMLIDSDLD
LAITDGPIEHPLLASQKAFD
ERLVLLMPADAGEPDAATLAPLEFYTFSRECSFRLKVDHWLASRGLKPRMTLEMESYA
AMAACVQAGCGVA
>33_Acetobacter tropicalis
KLSQLRFFCAIVEGGTIAAAARKMNCVPSNITMRLQELEMDIGQDLFVRDRKKLLITPAG
RLFYREAREIVSRAERLPSL
WSKPKSRGILKVGALDVAFLTCLPKHVPQFIMDHPDVELNVLQHSSFVLERMLSEEEID
LAITDGPVENTLLEGSFAFHQ
SLYLVMPQHIKKITPEIMRSSDIYLFNRDCFYRRCAELWMDQKGYQPRSVLTMESYDLI
LSCLDVGIGLACMPESVVAQF
RTHGRKLNILRMDGIGPTDVYFVWRKHAKNNIIQAFLDYMRED
>34_Acetobacter senegalensis
KLSQLRFFCAIVEGGSIAAAARKMNCVPSNITMRLKELETDLGQDLFIRDRKKLLITPAG
RLFYREAREIVSRAERLPSL
WSRPKSRGILNVGALDVAFLTCLPKHVPQFMMDYPDVELNVLQHSSFVLERMLSEEKI
DLAITDGPVENTLLEGSFAFHQ
SLYLVMPEHIKKITPDVINRSDIYLFNKDCFYRRCAELWMEQKGYQPRSVLTMESYDLIL
SCLDMGIGLACMPESVVAQF
RVRGRKLNIFKMDDIGTTDVYYVWRKHGKNDIIQAFLDYMRED
>35_Plesiomonas sp.
MNNSQLAAFCKVVESESVTAAAAQLFCVPSNITKKIKELESEFEVMLFTRDRNRLLVTP
EGRIFYQNAKQALQLEEQGRQ
LFTRDQLRGILHIGALDIALSHYLPTQIAAFRAQHQDIQLNITPGYSLDLEYQLLKGERDII
FSDGPIQHPLLQSQLAFA
EELVLIGADFEPALLSQRDLYVFTRQCHYRHLIDRWIKQHAVMPRAVLEIESYPVIFACV
RAGLGFALVPKS
>37_Acetobacter indonesiensis

KLTHLRFFCAIVEEGSIASAAHRMNCVPSNITMRLQELETDIGQDLFVRERKKLLITPAG
RLFYREAKEIVSRAERLPLL
WSQPQSRGILKVGALDVAFLTCLAKNVPDFIMAHPDIELNVLQKPSFILEKMLSNEEIDL
AITDGPVENTVLEGEFAFSQ
SLYLVMPEHIKTLSPDSISRSDMYLFNRDCFYRRCAERWLERKNYTPRSILTIESYDLIM
ACLDTGTGLSCMPESVVAKY
RVQGRKINAVKLDGIGRTDVYFVWRKHAKSDIISKFVD
>38_Serratia marcescens
FKRAQLAAFCMVVESESLTAAAERLFCVPSNVTKKLKELEMACGVTLFERERNRLSLT
PEGRALYHKAKRFLAIEERARA
QLEQDPVQGELKLGALDIALSHHLPQKLAHYRAQNRRVQLSVVQGYSLDLEFQLQKG
ALDVVFSDGPIEHPQMQSRRAFQ
ERLVFIGGAYRPALLAKSDLYVYSRQCHYRHLVDAWLQKQQVIPAALLEVECYDVIFAC
VTAGLGYAFVPESLARQRKVA
YIECGNEMRSDIYAIWRKASDSRILHDFIDAV
>39_Acetobacter cerevisiae
KLSHLRFFCAIVEAGSIAAAARRMHCVASNLTMRLKELEADLGQDLFIRDRKKLLITPAG
RLFYKEAKDIVSRADRLPEL
WSQGQPRGILNVGALDVAFLSLLPHCVPRFMREHPGIQLNILKKPSFSLERMLMSDEID
LALTDGPVENTLLEGTFAFSE
TLHLVVPEHVKKITPELIEQSDIYLFNKDCFYRRRAEQWLDEKAYHPHSILTLESYDLILE
CLGTGAGLSCMPESVIRQF
RERGQSLKTFRLDGMGPTDVYFVWRKHAKSQIIETFIE
>40_Acetobacter farinalis
KLSHLRFFCAIVETGSIAAAARHMHCVASNLTMRLRELEADLGQELFLRERKKLLITPAG
RLFYEEAKEIVSRADRLPAL
WAQSRLRGILNVGALDVAFLNLLPRCVPRFMVQYPGIHLNVLQKPSFSLEKMLVEEEID
LAVTDGPVENTLLEGIFAFSE
TLHLVVPETISAITPELIEQSDMYLFDRDCFYRRCAEQWLERNGYQPRSILTLESYDLIT
QCLNTRKGLACMPESVIRQL
RARGQALKTFRLEGVGPTDVYFVWRKHAKNPVIEKFL
>41_Acetobacter persici
KLSHLRFFCAIVEAGSIAAAARQMHCVASNLTMRLKELEADLGQDLFVRDRKKLLITPA
GRLFYKEAKDIVCRAERLPKL
WSEGQPRGILTVGALDVAFLSLLPRCVPRFMRAYPGIQLNILQKPSFSLEDMLITEVIDL
ALTDGPVENTMLEGTFAFSE
KLHLVVPGHVKKITPDLIAQSDFYLFNKDCFYRRCTEQWLEDKGYQPRSVLTLESYDLI
LACLRAGEGLSCMPESVIRQF
RERGEALRTFRLDGSGPTDVYFVWRKHAKNQSVETFI
>42_Plesiomonas sp.
MNNRQLTAFCAVVESDSLTAAAARLFCVPSNITKKIKELESEFDVMLFTRDRNRLLLTP
EGRAFYDKARHALQLEEQGRQ
LFARHPIRASLHVGALDVALSQWLPEKVAHFRHLHPDVQLQLTPGYSLDLEYQLLKGE
RDLIFSDGPVQHPLLHSQLAFT

EELVLVGAEPEPAQLNQRDLYVFTRQCHYRHLIDHWIAQHPFTPRAVLEIESYPVIFACI
RAGLGFAFVPKTLAEAEQLP
WSAKLEGIRSDIYAIWRCTHA
>43_Gluconobacter japonicus
LRLAQLEMFCAVVETGTVVAASRMLNCVASNVTARLRELEQLLGQDLFTREKNRLFLT
PEGRLFHQRAKAVVDQARGLTD
LFSEGSPKGVLTVGALDVALKTYLPDRVPRFLASAPGVEIRILIRPTDTLERMLMDGDID
LALTDGPIKHPMMSSAFAFR
ERLAIVLPKNQDILLSGQAVFLFNTDCLYRRYFEAWLEQGGVGGGLIHTIESYDVILACV
EAGLGISCIPQSVLSALQAK
RCEQVNVLYPEDLQGSDIYAVWREPGLGPLGQMFVNSLQA
>44_Gluconobacter thailandicus
LRLAQLEMFCAVVETGTVVAASRKLNCVASNVTARLRELEQLLGQELFAREKNRLLLT
PEGRLFYQQAKAVVDQARQLTD
LFSEGTPKGVLTVGALDVALKKYLPDRVPRFLASAPGVEIRVLIRPTDTLERMLVDGDID
LALTDGPIEHPMMGSAFAFR
ERLAIVLPKGQDALRSGQAVFLFNTDCLYRRYFEAWLEEGGVGGGLIHTIESYDVILAC
VEAGLGISCIPQSVLSVLKAE
RHADVNVLYPDDLQGSDIYAVWREPGLGPLGQILVDSL
>45_Acetobacter orleanensis
KLSHLRFFCAIVEAGSLAAAARRMHCVASNLTMRLKELEADLGQDLFVRERKKLLITPA
GRLFYKEAKDIVSRADRLPQL
WAQTLPRGILNIGALDVAFLRLLPQCVPRFIRDYPGIQLNILQKPSFSLERMLVSDEIDLA
LTDGPVENTLLEGTFAFSE
TLHLVVPEHVRKITPDVIEHSDIYLFDKDCFYRRCAENWLESKNYHPRSIVTLESYALIM
ACLKTGAGLSFMPESVLRQF
RKEGENLRTFRLDGLGPTDVYFVWRKHAKSQIIETFIE
>46_Acetobacter sp. DsW_063
LKLSQLQFFCAIMEEGSIVAASRRMHCVASNITARLKELEQILGQELFLREKGRLIVTPA
ARLLYREVTPLVAGLGRLPH
LFDADKPRGILKVGALDVALRHFLPERLPAFAAAYPGVELTLLTRPSYTLERMLSDGEL
DLVLTDGPIQHPLLDSRFAFS
ESLSLVTPRSIKAVDDIDWLTTTVFLFNTDCFYRNRFEGWLNARGLETPAIQTIESYDVL
RACVAAGLGVSCFPDSMLHE
RSENDGGNVRVFRPEDLDPGPVYFVWRREALS
>48_Rosenbergiella australiborealis
LKLPQLKIFCALVESGSVVQAAKKMHCVPSNISTRIRELEESLTVNLVNRDKQRLTLTPE
GRAFYPQAQKLLRQSEECLN
FFKPESLHGHLRLGALDVALNQQLEKLMITFMKQNPNVTVSLSCFSSLPLMDRLLAGE
VDMILVDGPIDHPVLESQLFAP
ERLRVVTHCATLDEFSSKVAALTLYTFGEHCFYHVLIEDWLASQQLQARQQCDIESYS
MILAAIHQQLGFTVMPESFIND
NPQVNGLYCYPLTDISSCDIYLVWRRYTASPVEKTFL
>49_Rosenbergiella collisarenosi

LKLPQLRIFCALVETGSVVQAAKKMHCVPSNVSTRIRELEESLAVNLVNRDKQRLTLTP
EGRAFYPQAQKLLQQSEECLN
FFKPDSLHGHLRLGALDVAINDHLQRVMVDFMEVHPDVTVSLSCHSSLPLMDRLLAGE
VDMVLVDGPVTHPVLASQLFAP
ESLSLVTHCATMAEFAERVAQLSLYTFGEHCFYHVLIEDWLAHQQLQARQRSDIESYP
LILAAIHQGLGFTVMPQSFIDN
HPQIAGLQCFPLTDISACDIYLVWRRYTTSPVEKAFLSINA
>50_Gluconobacter cerinus
LRLAQLEMFCAVVETGTVVAASRKLNCVASNVTARLRELEQLLGQELFAREKNRLFLT
PEGRLFHQQAKTVVEEARHLTD
LFSEGVPRGILTVGALDVALKEYLPSRVPQFLAAAPGVEIRILIRPTDTLERMLVDGEIDL
ALTDGPIEHPMMGSAFAFR
ERLALILPEGQKTLQSGQAVFLFNTDCLYRRYFEAWLEQKGFGGGLIHTVESYDVIMAC
VEAGLGISCIPQSVLSTLGAV
RKTDIAVQYPTDLQGSDIYSVWREPRVSVLARLFMDSLGS
>51_Iodobacter arcticus
SQLKALCAVIEQGGVSAAANTLFCVPSNITKKIKELESEFGLCLFHRERNRLILTAEGRT
FYQKAQEYLALHAQGRSLFY
DEPIVGELRIGALDIALSHHLPEPIAHYRIAHPEIKLNIVHGHSLELERQLQAGELDLIFSD
GPIEHPLMGGVLAFTEQL
VLVGAEPVEIPKQTLYAFPSTCHYRHLIEASLNMQGIRPQTLLEIESYPIIFALIQAGKGC
AFIPRSLAEAQGVPHYSEP
IQSNIYALWRKGAICHLGSEFIEKV
>52_Acetobacter fabarum
LKLAQLRFFCAVMEAGSVVAASRRLNCVASNVTARLRELEQSVGQPLFLREKGRLVET
PSGRAFYAQAREVVQKASALEH
FFKPDVPRGVLRLGALDVALQHFLPQCLPSFMQTFPDVELTVLKRASYTLEQMLMERA
VDLALTDGPIQHPLLESRLVFT
ECLRLVAPAQIASPGDIDWANTDVLLFNSDCFYRSVFEAWMQKNGVVPRRVQTIESYH
VIFACVRAGLGVCCCPDAMLPA
AHAHGLHAMTPPDLPEAPVYSVWRRDGVTPLLMRFVDYLG
>53_Pseudomonas putida
MDGELDLILTDGPIEHPLLESRLAFRERLLRVVPKAFSSPTPEDLAGLELYVFGRTCHY
RQQVDNWLERSGIQPRAILEI
ESYPSIFACIAQEIGFACVPESYIDRFASQADTFQADHVPSLDSSDIYFVWRKNQQSPLI
KNFIDAI
>54_[Acetobacter pomorum
LRLAQLQIFCAVMEEGSVVAASRRMHCVASNVTARLKELEQLLGQELFAREKGRLVPT
PEGRLFYHEAREVVEKAHNLAS
FFQPDVPRGILRIGALDVALAHFLPQYMPGFMATHPDVEIRLLRRASYTLEHMLEDQEL
DLAITDGPIAHPLLESRFAFT
EELVLVAPPNIQNLHDIHWPGVTVFLFHTDCFYRSAFERWMAEQELTPRAIQTVESYD
VIRACVRDGIGISCFPKSIFAQ
QTPHDVRVLAAPGLRAAPVYCVWRRNGAKPLLHRFVNTI

>55_Plesiomonas sp.
TKKIKELESEFEVMLFTRDRNRLLVTPEGRIFYQNAKQALQLEEQGRQLFTRDQLRGIL
HIGALDIALSHYLPTQIAAFR
AQHQDIQLNITPGYSLDLEYQLLKGERDIIFSDGPIQHPLLQSQLAFAEELVLIGADFEPA
LLSQRDLYVFTRQCHYRHL
IDRWIKQHAVMPRAILEIESYSVIFACVRAGLGFALVPKSLAESEHLAYLVQPDRINCDIY
TIWRSTHASPLLQDFIHSI
TA
>56_Acetobacter cibinongensis
KLSQLRFFCAIVEMGSIAAAADKMHCVASNITMRLKELENDIGHPLFTRENKSLIITPAG
RKFYQDVKPLVEKCDELPSL
WAQNKQRAIFKIGALDVAFLTCIYQKLPQFIKNYPDIDLSLIKGSSFALERMVLEQAIDLA
VTDGPVQTTQLQSVFAFHQ
PLYLVMPQGMKKERADIIKKSNFYLFNRDCFYRQRVEVWLEQKGYQPRSIQTIESYDVI
TACLDLGNGFSCMPESIIRQL
RKQGHVLNVRPMPEIGPTDVYFVWRKTAASEIREKFI
>57_Acetobacter syzygii
LRLAQLRFFCAVMEEGSVVAASRRLNCVASNVTARLRELELIVQQPLFLREKGRLIETP
AGRAFYDQAREVVHKASALEH
FFQPDVPRGVIRLGALDVALQHFLPQRLPAFMHDCPDVVLHVLKRASYTLEHMLVEREI
DLALTDGPIEHPLLESRLVFT
EHLRLVVPDYVQTLQAIDWARTDVLVFNTDCYYRSVFESWMQNNGVAPRRIQTIESYP
VIFACVRAGLGVCCCPDVMLPQ
NPAPGLQVMTPPDLPVAPVYSVWRRAGADPLLMRFVEHLGA
>58_Acetobacter musti
LRLSQLQFFCAIIEEGSVVSAARRMNCVASNITARLKELEQILGQQLFEREKGRLVVTPA
GRLFYREAAEVVTRARRLSQ
MFEPDVPAGVLAVGALDVAMRSFLPERVAAFLKAHPRMELNLLIRPSYTLERMLIDGEL
DLVLTDGPIVHSLLDSRFAFD
EALHLVVPLHIATVDDIDWEKTTIFLFNTDCFYRRHFEAWLARRGVLNPVVQTIESYDVI
RACVQAGLGISCFPHCMLRE
NRSGIKVLSPADLEPGPVCFVWRRNGVSPSLTRFVASM
>59_Acetobacter pasteurianus
LRLAQLQIFCAVMEEGSVVAASRRMHCVASNITARLKELEQLLGQDLFAREKGRLIPTP
EGRLFYHEAKEVVEKAHSLAN
FFQPDVPRGILRIGALDVALAHFLPQYMPAFMAAHPDVEIRLLRRASYTLEHMLDDQEL
DLAITDGPIVHPLLESRFVFT
EELVLVAPQNIRSLQDIHWPSTTVLLFHTDCFYRSAFERWMAEQGFVPRAIQTVESYD
VIRACVRDGIGISCFPQSIFAH
KAPDDVRVLAVQGLRPAPVYCVWRRNGAKPLLHRFINTI
>60_Gluconobacter kondonii
LRLAQLEMFCAVVEEGTVVAASRRLNCVASNVTARLRELELLVGQDLFLREKNRLILTP
EGRLFYRQALPVVSGAHSLTD

LFSEGSPRGILNIGALDVALEAYLPQIVPEFLKVCPGVELKMFLRPSYTLERMLVDGEID
LALTDGPIVHPLMESRFAFH
ERLALAMPVGQDGARDGQHVFLFNTDCFYRRYFETWLERHGVSNAIIHTVESYDVILA
CVQAGLGISCVPHSIILQMQSR
CDEGITFTYPEDLKGSDVYFIWRDPGLNALGRRFLDHAFAN
>61_Endobacter medicaginis
LKLAQLRFFCAVVEEGSVAAAARRLNCVASNVTTRLRELEQSLGEELFQRERGRLLLS
PTGRIFYREALPVVASAARLAR
LFDDAGQPRGLLRIGALDVALDGFLTRHLPGFIAAHPGIEFSLLRRPSYVLERMLADGEI
DLALTDGPIAHPLLDGRAAF
AEDLCIVTAAAIGDLSAVDWSSAQVFLFDTDCFYRDLFTRWLNTHELRPAGIQTIESYDL
IAACVRAGLGVSCVPATMIP
ARNAREGLAIHRPDDLGPSEVHFVWRSE
>62_Acetobacter sp.
LKLAQLRFFCAVMEEGSVVGASRRLNCVASNVTARLRELEQIVGQPLFLREKGRLIETP
AGRAFYTQARDVVQKAVGLEH
FFAPDVPRGVLRLGALDVALQHFLPQRLPSFMQDCPDVDLCVLKRASYTLEQMLMAR
EIDLALTDGPIVHPLLDSSLVFT
QRLRLVTPAHVAVPEQIDWARTDVLLFNTDCFYRSVFESWMQKNSVVPRRIQTIESYQ
VIFASVRAGLGVCCCPDVMLPA
AHSFGLHALTPPDLPEAPVYSVWRRDGATPLLKRFVAHLTA
>63_Acetobacter ghanensis
LKLAQLRFFCAVMEEGSIVAASRRLNCVASNVTARLRELEQVVNQPLFLREKGRLIETP
AGRAFYDEAREVVRQASALEH
FFEPDVPRGLLRLGALDVALRHFLPQKLPSFMQACPDVELHLLKRASYTLEQMLMARE
IDLALTDGPIEHPLLESRLAFT
EHLRLVAPAHVADFAQINWAETDVLLFNTDCYYRSVFENWMRRNGIVPRRVQTIESYH
VIFSCVQAGLGVCCCPDVMLPV
AYAQGLHVMTPPDLPEAPVYSVWRRDGTMPLLMRLVEHLSA
>64_Rosenbergiella epipactidis
LKLPQLRIFCALVETGSVVQAAKKMHCVPSNISTRIRELEESLGVALVNRDKQRLTLTPE
GRAFYPQAQQLLKQSEECLH
FFKPESLKGHLRLGALDVALNSRLQQIMIGFMQQYPEVTVSLSSHSSLPLMDRLLAGEV
DMVVVDGPVEHPVLESRLFGP
ESLCLVTHCASLETFIEQVSQLTLYTFGEHCFYHVLIEDWLAAQQLEARQQCDIESYPLI
LGAIRQKLGFTVMPRSFIEN
NDEVKGLYCYPLDTLSSCDIYLVWHRYSASPVEKAFLSMV
>65_Acetobacter sp. LMG 32666
LKLAQLRFFCAVMEAGSIVAASRRLNCVASNVTARLRELEQSVGHPLFLREKGRLIETP
AGRAFYAQAHEVVQKASALEH
FFEPDVPRGVLRLGALDVALHHFLPQRLPSFMQAFADVELQLLKRASYTLERMLMERE
VDLALTDGPIEHPLLESCLVFT
QHLRLVAPVHVASVAAIDWASTDVLLFNTDCFYRSVFESWMKKNRVTPRRIQTIESYQ
VIFACVRAGLGVCCSPDVMLPA

PHAQGLHVMTPPDLPEAPVYCVWRRDGATPLLMRLVDHL
>66_Gluconobacter oxydans
LRLAQIEMFCAVVEEGTVIAASRRLNCVASNVTARLKELELLVGQELFLREKGRLILTPE
GRLFYQQATPVVAGARSLTD
MFSEGTPRGVLNIGALDVALEGYLPRIVPEFLKACPGVELKMFLRPTYTLERMLADGEI
DLALTDGPIVHPLMESRFAFR
EKLALAMPEGQKGARDGQHVFLFNTDCFYRRYFETWLERHGVSNAIIHTVESYDVILA
CVQAGLGISCVPRSIISQVKRR
RREGISFTYPEDLKGSEVYFIWREPGLNDLGRRFLD
>67_Escherichia coli
YVFGHTCHYRRQVDRWLAESGIQPRATLEIESYPSLFACIEAGLGFACVPESFVARMP
STRRGFHAEPVAGLDSSDIHYV
WRKQQASPLIQGFIDSIGAD
>68_Gluconobacter morbifer
LRLAQLEMFCAVVEEGTVVGASRRLNCVASNVTARLRELELLLERDLFVRTKGRLNLT
PEGRLFYRQAVPLIRGAQDLAG
LFADKTPRGILTVGALDVALQGYLPSVVPQFLENCPGVELRMLMRPTYTIERMLMDGEI
DLALTDGPIVHPMMDSRFAFR
EDLALVLPQGQEVPQEGQHVFLFNTDCFYRRYFEIWLRKQGVTSSIIHTVESYDVILAC
VRAGLGISCFPQSILAGVAGE
RRQGMEILYPADLQGSEVYFVWREPGVTDLARQFVD
>69_Brytella acorum
LKLSQLQFFVAVLEEGSVVGASRRMNCVASNVTARLKELEQLLGQELFLREKGRLVITP
AARLFYREAKPVVLNAARLLH
LFEADAPRGILTVGALDVALRRFLPERLPAFATAYPGVELTLLMRPSYTLERMLADGDL
DLALTDGPIQHPLLNSRFAFA
ESLSLVVPSHITAIKEINWPTTTVFLFNTDCFYRSCFERWLKKRGIETPVIQTIESYDALH
ACVAAGLGVSCFPDSMTSP
QDCDDLNGVRFLKPDDLASAPVYFVWRRDS
>70_Gluconobacter kanchanaburiensis
LRLAQIEMFCAVVEEGTVIAASRRLNCVASNVTARIRELELLLGQDLFFREKGRLILTPE
GRLFYQKAGPIVSGARDLTD
LFEEGRPRGILTIGALDVALEGYLPGIVPHFLADCPEVELRVLLRPTYTLERMLAEGEIDL
AVTDGPIVHPLMASRLTFH
EKLALAMPGGQTRLRAGPKDRQHVFLFNTDCLYRRFFETWLESHGVTNAVIHTVESY
DVILACVQSGLGISCIPRSILLQ
VRQDRRDGIDFTYPEDLQGSDVYFIWREPGLNDLGRRFLDHIPA
>71_Tatumella citrea
LKLHQLGFFCALVEEGSVAAAADKMCCVPSNISTRIKELESSLGVALVNRDKQRLTLTP
EGRAFYPKAQRLLEQSQQCLT
FFQHSLLQGQLQIGVLDVALKGPLQQAVVEFMHQNPQVQINICCHSSLTLMEKLLQGE
LDMILVDGPIEHPALKAEFFAP
ERLSLVTHLADRQQFRDHASGLTLFSFGEKCFYHLLVRQWLSQQNLRYARQSDIESYD
LILSAVHRRLGFTVMPQTFIDN

NPLLSGLHLFPLDNIPSCDIYLVSQSVEQSALAEAF
>72_Gluconobacter wancherniae
LRLAQLQMFCAVVEEGSVVAASRRLNCVASNVTTRLRELEQLLGQELFLRDKGRLSLT
PEGRLFYQKASPLVISAQGLTD
FFSDASPKGVLVVAALDVALQGYLARRVPSFIAAYPGVELKLLLRPTYTVERMLTDGEI
DLGLTDGPISHPMIGSLPAFQ
ERLALVVPKGVTERGLIGLHVFLFNTDCFYRRYFETWLEGQGLSAPLIHTVESYDVILG
CVRAGLGMSCFPQSIVEGLSA
EQRQGVEVLYPEDLRGSEVCFIWREPGLSDLGRRFVQHIDED
>73_Plesiomonas sp.
MNNSQLAAFCKVVESESVTAAAAQLFCVPSNITKKIKELESEFEVMLFTRDRNRLLVTP
EGRIFYQNAKQALQLEEQGRQ
LFTRDQLRGILHIGALDIALSHYLPTQIAAFRAQHQDIQLNITPGYSLDLEYQLLKGERDII
FSDGPIQHPLLQSQLAFA
EELVLIGADFEPALLSQRDLYVFTRQCHYRHLIDRWIKQHAVMP
>74_Serratia sp. CY81593
GVTLFDRERNRLSLTPEGRALYHKAKRFLAIEERARAQLEQDPVQGELKLGALDIALSH
HLPQKLAHYRAQNRRVQLSVV
QGYSLDLEFQLQKGTLDVVFSDGPIEHPQLQSRRAFQERLVFIGGAYQPALLAKSDLY
VYSRQCHYRHLVDAWLQKQQVI
PAALLEVECYDVIFACVSAGLGYAFVPESLARQREVAYTECGNEMRSDIYAIWRKASD
SRIVQAFIEAV
>75_Halomonas caseinilytica
IKLNQLRHFRAVVEQGSVAQAAKTLFCVPSNVTARIRELEEVLGTVLFHREGKRMVLTP
NGRRFYEEAVSIIQRVDHAAG
LFDEKQAQWEMYLGALDVSLFGFLPTRLPKFRVQYPQVKLDICCRPSSELEAAVLADE
MDLVITDGPIEHPLLESCLAYR
ERMCLITPLNRGGPGDAMTDLDIFGFGTNCSYRVRMDEWVRQQGVGSGRSIRVESY
HVIFQYVKAGQGLSWFPLSMLDQI
DPGNAVERYEMGDSELFFVWRRGREQPEISTFMN
>76_Streptomyces sp. NPDC048430
LKRAQLVTFCMVVESESLTAAAERLFCVPSNVTKKLKELEMACGVTLFDRERNRLSLT
PEGRALYHKAKRFLAIEERARA
QLEQDPVQGELKLGALDIALSHHLPQKLAHYRAQNRRVQLSVVQGYSLDLEFQLQKGT
LDVVFSDGPIEHPQLQSRRAFQ
ERLVFIGGAYQPALLAKSDLYVYSRQCHYRHLVDAWLQKQQVIPAALLEVE
>77_Alcanivorax jadensis
IKLNQLRHLQAVVDRGSVAGAAKVLFCVPSNVTARIRELEEALGTQLFHREGKKLLLTP
NGRRFYREAVVILKRVDQAAR
LFDETQAQWELSLGALDVSLFQFLPARLPAFRAAYPQAQLDISCRPSSELEAAVLAGE
RDLIITDGPIEHPLLESRWAYR
EKMCLVTPAGRGGEGDTLSDLDMLGFGRNCSYRVRMEEWANRQGVGSGRGQRIES
YHVILEYVKAGLGFSWFPLSILDQL
DPERRVERYDVGALGDSNLFFVWRRGREQP

>79_Janthinobacterium lividum
LKLSALEMFCAVIEEGSMAAAARRVNCVPSNIVARMKELESDLGEPLFHREKGKISATP
FARIFHRETRDLLDRADQLTQ
YFQQDATLGLLKVGALDVAMLDYLPARLPRFLKDHARSEISLQCQPSFALERMLGASEI
DVALTDGPITHPLFESCPAYT
EDLYLVANQHASRITDELLRAPVFLFDEDCFFRPHFKSWLANRGVAQTTIQTIESYETIT
ACVAAGLGISCFPGSVAARI
DEKKQGVRFLKPDDLAPSGVHFVWRRNGMTHLLDRFI
>80_Pantoea sp. At-9b
MQLSQLRAFCAVAQTGSISGAARVLNRVPSGITVRIQQLEQDLGCELFLRDRQGMSLS
QNGRLLLEHAQRILDLTDSTRL
LMREEDLGGKLVIGALDVVLVDFMPALIGTFRKRYSGIGLDIRHEASEVLVQHVADGTL
DIALTDGPVQSKALESRLAFV
DEMLLITELNHPAVTTPSDLQCSELYGFRHDCSFRFRMDRWLAEERINHLPVTEIESYH
TMLACVTAGMGAAWVLRSVLQ
TLPGHQLVRSHSLGKVGYTEIHYLWRSGHLSP
>81_Enterobacterales bacterium CwR94
MDMTQLKMFCVVAETGSVSQAAEQLHRVPSNISTRIKQLETELGTPLFIREKLRLRLSP
DGHLFHQYAREILAKIDEAVS
ALSGRQPRGRFALGALESTAAVRIPPLLARYHQRFPEVELEFTTGPSDELLQALLEGRL
DAVFVDGPIETDVLTSIAVWQ
EEMALVAAKDHPPIQTAQDVAGAPLYAFRRTCSYRRQFELWFARQGVSPGRIYEMES
YHGMLACVSAGAGLALAPRSLLA
HLPSAQNLSIWPLDGTLSAVPVSLVWRKESHS
>82_Mangrovibacter sp. MFB070
MQLSHLRAFCAVAQTGSVAAAAKVLNRVPSGITVRIQQLEQDLGCELFLRDSKGMSLS
PNGRLLLEHAQHILDLTDSTRL
LMREEDLGGRLVIGALDVVLVDFMPALIGTFRKRYQGVGLDIRHEASEVLVQHVADGS
LDIALTDGPVHSKSLESKLAFV
DEMLLITERNHPPVKSPADLQCSELYGFRHDCSFRFRMDRWLAEAGISHLPVTEIESY
HTMLACVTAGMGAAWVLRSVLQ
TLPGHQQVQSHTLGPVGYTEIHYLWRSGQLSPNARKLIET
>83_Acetobacter oeni
LRLSQLQFFCAIIEEGSIVAAARRMNCVASNITARLKELELLLGQQLFEREKGRLAVTPS
GRLFYREARELVAQAHRLSH
MFEPDVPRGILAVGALDVAMRSFLPERVPAFLRAHPRMELNLLIRPSYTLERMLADGEI
DLALTDGPVVHSLLDSCFAFD
EPLRLVVPPHIGALEDIDWASTRVFLFNTDCFYRRRFEAWLRDRGIHDPVVQTVESYD
VIRACVHAGLGISCFPHCMLRD
DTGGIRVLAPDDLRPGPVYFVWRRDGVSEPLTRFVASM
>85_Acetobacter tropicalis
IGQDLFVRDRKKLLITPAGRLFYREAREIVSRAERLPSLWSKPKSRGILKVGALDVAFLT
CLPKHVPQFIMDHPDVELNV

LQHSSFVLERMLSEEEIDLAITDGPVENTLLEGSFAFHQSLYLVMPQHIKKITPEIMRSS
DIYLFNRDCFYRRCAELWMD
QKGYQPRSVLTMESYDLILSCLDVGIGLACMPESVVAQFRTHGRKLNILRMDGIGPTDV
YFVWRKHAKNNIIQAFLDYM
>86_Novosphingobium profundi
MQIPQLRAFCAVAQSGSISTAARVLNKVPSSITVRIQQLEEDLKQQLFLRERKRMSLSP
AGRQLLEHAQRILDLTDGTRA
LMRDEEISGSLTVGALDVVLVDYMPRVIGYMRQRHRGIHLNVRHQASEDLVAHIADGS
LDIALTEGPISSKALRSRIAFV
DEMLLVTDPGHRDVVTPTDLECAELYGFRHDCSFRFRMDRWLASGGRSDLPVIEIESY
HTMLACVSAGMGAAWMLRSILK
TLPGHNQVKAHSLGDHGRTEIHFVWRDGHLSRNAELFMDA
>88_Tatumella saanichensis
LKLHQVSFFCALVEEGSIAAAADKMCCVPSNVSTRIKELEASLGISLVNRDKQRLTLTPE
GRAFYPKAQQLLEQSRMCRS
FFSHDQLQGTLQLGVLDVALKGRLQAPVIEFMQQNTQVQINISCDSSLPLMEKMLQGE
LDMILVDGPVQHPALKTDFFAP
ESLSLVTHLADRQTFAKQAASLTLFSFGEKCFYHQLARQWLSHEHLSFARQSDIESYD
MILSAVRKGLGFTVMPESFMRD
NPALDGLYHFPLENIAPCDIYLAAPALEQSALANAFYQQL
>89_Neorhizobium sp. NCHU2750
MQIPQLRAFCAVAQTGSVSAAAKVLNKVPSGITVRIKQLEDDLGRELFLRDRQRMSLS
PAGRQLLEHAQRILDLADGTRA
LMRDEEISGPLVIGALDVVLVDYMPALIGRLRQRHRGITLEVRHQASEDLVAHVADGSL
DIALTEGPIVTKSLQSRVAFV
DEMLLVTEIGHPDVTSPEELDCTELYGFRHDCSFRFRMDRWLAAGGRAHMPVLEIESY
HTMLACVSAGMGAAWMLRSVFK
TLPGHHQVKAHSLGDAGRTEIHFVWRDGHLS
>90_Sphingobium sp.
MQIPQLRAFCAVAQTGSITAAARVLNKVPSSITVRIRQLEEDLGRELFLRESKRMFLSPA
GRQLLEHAQRILDLADGTRA
LMRDEESSGPLVVGALDVVLVDYMPRLIGRMRQRHRGIHLNVRHQASEDLVAHVADG
SLDVALTEGPVISKALKSKVAFV
DEMLLVTAPGHRDVTTPADLDCTELYGFRHDCSFRFRMDRWLSCGNRADMPVIEIES
YHTMLACVSAGMGAAWMLRSILK
TLPGHHQVKTHSLGEAGRTEIHFVWRDGHLS
>91_Tatumella terrea
LKLHQLSFFCALVEEGSIAAAADRMCCVPSNVSTRIKELEASLEVPLVNRDKQRLTLTP
EGRAFYPQAQQLLEQSRQCRT
FFRHALLQGTLRLGVLDVALKGRLQAPVIRFMQQNPQVQINICCDSSLPLMERLLRGDL
DMILVDGPVQHPALKTHFFAP
ESLSLVTHLADQQQFKAQAAGLTLFSFGESCFYYLLARQWLSQQELSYSRQSDIESYD
LILNAVRQGLGFTVMPQTFIAG
NPALDGLYHFPLDDLPPCDIYLAAPANEQSALAQTFYRHLLSD

>92_Luteibacter anthropi
MELSQLRAFCAVAETGSVIGASRALNKVPSSITVRIQQLEQDLGCPLFLRERQRLSLSP
DGRRLLEHAQRIVGLADSTRD
LMRNSEVGGRLVVGAVDVALMAFMPALIGRYRSRHRSVELDVRRGASDVLIEQVTDG
ALDIALGEGPVSATGLASRLAYT
DDVVLVTEFGHPPVDSPRDLRCNELYGFRHDSSFRYRVDAWLEASGRQLLPVVEVGS
YHTMLACVTAGMGAAWIPRSVLA
TLPGHPQVRAHALGEAGRSEIHFVWRKTHLS
>93_Serratia
MQLSQLRAFCAVVSTGSITAAAKVLNRVPSGVTIRIQQLEDDLQCQLFVREKKRLTLSP
AGRQLLEYAPQLIEQADKLRS
LMRNEQAQGCLTIGAIDLGLIAFMPTLIGRFRERYRGVNMDIRCEPSERLVEQVLDGTL
DIALTDGPVQSPSLESDYAFS
DQLVLITEKNHPIVTNASELCCQEVYGFRQNCSYRLRLDSWLLKTPQAHLKVVEMESY
HTMFSCVSAGVGAAWIPQTVLD
GLPGRDSVKVHTLGPSAQSDLYFIWRK
>94_Pseudomonas sp. LP_7_YM
MQLAQLRAFCAVARTGSVTAAAKELHRVPSGVTLRIQQLEEDLDCKLFVREKQRLTLS
PFGRALLARAQQIIELSDGAKA
LIREEQVGGHLTLGALDVGLVAFMPQLVGRFRQRHPQVSMDIRSDVSEVLMGQVAEG
LLDIALTDGPVQSPALDSRYAFS
DHLVLITEQAHGPVHAASDLRCREIYGFRQNCSFRLRLDRWLQASGGEHLQMIEMESY
HTMLACVTAGAGAAWVPRAMLQ
TLPGRAAVRVHSLGAAGRSDLYFVWR
>95_Sphingobium yanoikuyae
MQISQLRAFCAVAETGSVTGAARILNKVPSSITVRIRQLEEDLDRILFLRERKRMSLSPA
GRQLLEHAQRILDLADGTRA
LMRDEGISGPLLVGALDVVLVDYMPLLIGRMRQRHRGMTLNVRHQASEDLVAHVADG
SLDIALTEGPVRSKALRSKVAFV
DEMLLVTDPGHPDIGGPGDLQCAELYGFRHDCSFRFRMDRWLAAGGRGDMPVIEIES
YHTMLACVSAGMGAAWMLRSILK
TLPGHLQVKAHSLGEAGRTEIHFVWRDGHLSP
>96_Alcanivorax sp.
IKLNQLRHLQAVVDRGSVAGAAKVLFCVPSNVTARIRELEEALGTQLFHREGKKLLLTP
NGRRFYREAVVILKRVDQAAR
LFDETQAQWELSLGALDVSLFQFLPARLPAFRAAYPQAQLDISCRPSSELEAAVLAGE
RDLIITDGPIEHPLLESRWAYR
EKMCLVTPAGRGGEGDTLSDLDMLGFGRNCSYRVRMEEWANRQGVGSGRGQRIES
YHVILEYVKAGLGFS
>98_Kluyvera cryocrescens
MNSTALNLFKTVAETGSISAAARQLHCVSSNVTTRLKQLESSLDVSLFIREKNRLHITPE
GELLLRYANKIMALLEEAQL
SMQGTKAAGPLRIGAMETTAAIRLPPFLARFHREMPAVELNLKTAPTELLMQQVNNNE
LDIAFVADNPRLHDENSSLQSQ

VFCQETLVLVTAADQRAVTCAADLQITRPLVFVKGCNYRSRLEQWLESLGVEHATPLE
FGSFQAILGCVSAGMGIALLPE
SVVRQYEDNFSIRSHAISPQMGRVNTLIVWHKNREISPVISGFISFMAAE
>99_Erwinia sp. 9145 PtrR
MDITQLEVFRAIAEEGSVTAAAEKLHRVPSNISTRLRQLEEELGTLLFNREKLRLHITDA
GRTLLDYATRILDLTAEAKQ
RIAGGQPAGVFTLGAIESTAAVRLPPLIARYHQQWPEVELSLSTGPSGAMTDGLLAGE
FSAVFIDGPPMHPQLDGIAAFP
EEMVLISALNHPAITGAHALSGATLYAFRTNCSYRRLFENWFARANAMPGKIFEMESY
HGILACVSAGAGLALIPRSMLE
TMPGRDSVQAWPLADKAGHLDIWLVWRKN
>100_Clostridium perfringens
VVESESLTAAAERLFCVPSNVTKKLKELEMACGVTLFDRERNRLSLTPEGRALYHKAK
RFLAIEERARAQLEQDPVQGEL
KLGALDIALSHHLPQKLAHYRAQNRRVQLSVVQGYSLDLEFQLQKGTLDVVFSDGPIE
HPQLQSRRAFQERLVFIGGAYQ
PALLAKSDLYVYSRQCHYRHLVDAWL
>101_Serratia sp. EWG9
MAQAERSKQLFAGEHQHGLLNLGALDFSLVSHLPARIARLRHLQPHLQINVLSRDSLVL
ERMLIDSTLDLAITDGPIEHP
LLASQKAFDERLVMLMPADAGQPDAATLAALEFYTFSRECSFRLKVDHWLASRGLKP
RMTLEMESYAAMAACVQAGCGVA
CVPGSLLPQILPAPGLKVVEMGEEGVSDLYFVWRRHQLSDELQTIL
>103_Caulobacter sp. (uncultured)
MQLSQLRAFHAVAQTGSISAAAKMLNRAPSGVTVRIQQLEQDLRCELFLRDRQGMSL
SPSGRLLLEHAQRLLDMADGTRA
LMRDEDPGGKLVIGALDVVLVDYMPALIGQFRQRFRNVKLDVRHEASEDLVQHVSDG
ALDVALTDGPVRSNALESRLAFV
DEMLLITERSHPEVRRPSDLRCAELYGFRHDCSFRFRMDRWLADGGVRDMPITEIESY
HTMLACVTAGMGAAWVLRSVLQ
TLPGHQQVQTHSLGEAGHTEIHFLWR
>104_Brevundimonas sp.
MKIDQLRAFCAVAQTGSVSGAARVLNRVPSGVTVRIQQLEQDLGCELFLRDRQGMSL
SPAGRQLLDHARRILDLTESTRS
LMKDEDVGGRLTLGALDVVLVDYMPALIGRFRSRYRGVDLDIRHQASEDLVQNVGEG
ALDLALTDGPVRSNALESRLAFI
DEMLLVTEQAHPPVGGPAELECGELYGFRHDCSFRFRMDRWLTEGKATDLPVVEIES
YHTMLACVTAGMGAAWMLRSVLE
TLPGHQQVRVHSLGEAGRTEIHFLWRS
>105_Erwinia psidii PtrR
MDLTQLEVFRAVAEQGSVTAAAERLHRVPSNISTRLRQLEEELGTELFTREKLRLHITD
AGRTLLDYACRILALTEEARQ
RVSGQQPAGIFTLGAIESTAAVHLPPLIARYHQAWPEVELDLSTGPSGDMIEGLLAGSF
SAVFIDGPPRHPLLEGVAVFP

EEMVIISSLNHPPVTSAKNIAGATVYAFRANCSYRRLFENWFADSASAPGKIFEMESYH
GIVACVSAGAGLAMIPLSMLE
NMPGRDSVQAWPIAEQRGKIAIWLVWQRGAASANLRAMVN
>106_Kluyvera sp. 142053
MNSTALNLFKTVAETGSISAAARQLHCVSSNVTTRLKQLETSLDVSLFIREKNRLHITPE
GQLLLRYANKIMALLEEAQL
SMQGTKAAGPLRIGAMETTAAIRLPPLLASFHREMPAVELNLKTSPTELLMQQVSNNEL
DVAFVADHQKLHDENSSLQSQ
ILCQETLVLVTAAEHPAVTQGTDLQVTKPLVFVKGCNYRSRLEQWLESLGVEHSTPLE
FGSFQAILGCVSAGMGIALLPE
SVVRQYEGNFSIRSHAISPQIGNVNTMVVWRKNREISPVINCFISFMTAE
>107_Escherichia coli
LELYVFGNTCHYRRQVDRWLAESGIQPRVTLENESYPSLFACIEAGLGFACVPESFVA
RMPSTRRGFHAEPVAGLDSSDI
HFVWRKQQA
>108_Pantoea osteomyelitidis PtrR
MDLVQLRMFCSVAESGSLARAAEQLHRVPSNLTTRLRQLEEELGVDLFIREKQRLRLS
PMGHNFLCYAQRILALSDEALS
MTRAGEPAGNFALGSMESTAATRLPGLLAAYHQRYPAVSLSLITGTSGEIIDRVREGTL
AAALVDGPVLHDELNGCIAFR
EQMVLITSLDHVPIRSARDAAGDTLFAFRNSCSYRLKLEAWYRDSNTQPDSVMEIQSY
HAMMACVAAGAGLAKIPASVLA
QMPDRARVQQHALPAQYRDTATWLIWRRDAFTPNVEAL
>109_Rouxiella chamberiensis PtrR
MDLTQLEVFVAVADEGSVTAAADRLHRVPSNISTRIKQLEEELGTPLFHREKLRLHISDP
GKTFLGYAKRILALADEAKQ
SVSGQKPNGIFTLGAIESCAAVHIPPVLARYHQAYPDVELDLSTGPSGDMIDGILAGDY
SAAFVDGPPRHPELEGVPAFD
EEMVLISALNHPAVTRAQDISGATIYAFRANCSYRRLFENWFAQDNAVPGKIYEMESYH
GIVACVTAGAGLALVPASMLA
SMPGKRGVKVYPLSEEMGKIKIWLMWRKGAAS
>110_Pseudomonas sp. PtrR
MEFSQLRIFQAVAEEGSITRAAERLNRVPSNLSTRLKQFEAQLGTELFIRERQRLSLSP
AGKVLLDYTARLMALHDEAEA
AIKGGEPSGNFVLGTMYSTAAIHLPAMLVRYHRAYPAVNLQVQAATSGDLIEGLLNGRL
DMALVDGPPEFSALDGIPLFE
ERMVVISDETQGPIHSAEDIQGLSVFTFRTGCSYRRRMESWCSSQQVAMGRVMEIES
YQSMLACVIAGAGVALIPQSMLE
TLPGRENVLVHELASPFDRATTWLMWRRGRVGANLNAWID
>111_Erwinia sp. unclassified PtrR
MDLTQLEIFRAIAEEGSVTAAAERLHRVPSNISTRLRQLEDELGVELFSREKLRLHITDA
GRTLLDYAGRILALTEEARE
RVSSQTPSGVFTLGAIESTAAVRLPPLIARYHQAWPGVELDLSTGPSGEMTEGLLSGE
YNAVFIDGPPKHPQLEGIPVFA

EEMVLISSLSHRPVTRAADINGATIYAFRANCSYRRLFENWFAQDEAAPGKIFEMESYH
GIVACVSAGAGLAMIPESMLQ
NMPGRDSVLAWPLAEQRGHLNIWLVWRKGRSSANLRAMVE
>112_Pantoea sp. PtrR
MDLVQLRMFCSVAESGSLARAAEQLHRVPSNLTTRLRQLEDEIGVDLFIREKQRIRLSP
MGHNFLNYAQRILALSDEALN
MARAGEPGGNFALGSMESTAATRLPALLAAYHQRFPAVELSLITNTSGEIIDKVREGTL
AAALVDGPVAHDDLNGCIAFN
EQMVLITSPEHAPIASARDVQGDTLFAFRNSCSYRTKLEAWYRESQTAPSSVMEIQSY
HAMVACVAGGAGVAMIPASVLA
QMPARARVQEHALPPAYRDTATWLMWRRDAFTPNVEAL
>113_Pseudomonas brassicae PtrR
MELSQLRIFQAVAEEGSVTRAAERLHRVPSNLSTRLRQLEEQLGVELFRRERQRLQLS
PAGKVLLDYTARMLALHDEAFA
AVQGGEPAGAFVMGSMYSTAAIHLPSLLARYHRRYPAVNLQVQTAPSGELLEGLLSG
RLDAALVDGPLALAGLDGVALCE
ERLVLITEADHLPVRSALDVAGRAVFTFRQGCSYRMRLEAWYAHDHAAMGRVMEIES
YPSMLACVIAGAGVALMSASMLD
SLPGRESVAAHPLQAPFDQATTWLMWRRGMCGANLSAWI
>114_Erwinia sp. HDF1-3R PtrR
MDLTQLEVFRAVAEEGSVTAAAERLHRVPSNISTRLRQLEEDLGTPLFSREKLRLHITD
AGRTLLDYAIRILALTEEARQ
RVTYQQPAGIFTLGAIESTAAVRLPQLIARYHQQWPDVELDLSTGPSGDMTDGLFSGR
FSAVFIDGPPKHPQLDGVAAFA
EEMVLISSLNHPPVPSAASVSGSTIYAFRANCSYRRRFENWFAAEHAAPGKIFEMESY
HGILACVSAGAGLAMVPRSMLE
SMPGRESVAAWPLAEDAGLLDIWLVWRKGYASGNLRAMV
>115_Chimaeribacter coloradensis PtrR
MDLTQLRMFCCVAETGSLAKAAEQLHRVPSNLTTRLRQLEQELGTDLFIREKQRLRLS
AMGHNFLGYAERILALSDEAMS
LTHAGEPAGQFALGSMESTAATRLPALLAAYHQQHPAVALSLVTGTSGHLIDGVRGGR
LAAALVDGPINHDDLNACIAFR
ERLVVISPADRPLAAHPVETPLTVFAFGTTCSYRHKLHAWLGQAGWVTGTTLEIHSYH
AMLACVASGAGIAVLPQSVLAQ
LPGHERVTANPLPDEVADTATWLIWRREAFSPNVRALKEII
>116_Pantoea sp. BAV 3049 PtrR
MDLTQLEIFRAIAEEGSVTAAAERLHRVPSNISTRLRQLENELGVALFSREKLRLHITDA
GRTLLDYALRILALTEEARE
RVSSQKPAGIFTLGSIESTAAVRLPPLIARYHQTWPEVELDLSTGPSGDMTDGLLAGQF
SAVFTEGPPRHPQLEGIPVFA
EEMVLISSLNHPPVSSAADIAGSTIYAFRANCSYRRLFENWFARDNAAPGKIFEMESYH
GIVACVSAGAGLAMIPQSMLQ
SMPGRDSVLAWPLAGKRGHLHIWLVWRKDSHSANLQAMVKLLKSD
>117_Pseudomonas aeruginosa strain

MQLSQLRAFCAVAHTGSINAAAKALNRVPSSLSVRIRQLEEDLGCQLFLREHQRLRLS
PDGRRLLEHARRILDLSDSTRA
MMRNEESGGRLVVGALDVVLVAFMPCLIGRFRQRHRAIELDIRSEASEALVQQVSDGV
LDLALSDGPVRSQTLESSLAFV
DEMVLVTELDHPPITSPRDLRCAELYGFRHDCSFRFRMDRWLEEADCLQQLPVLEIES
YHTMLACVSA
>118_Cernens ardua
MDINQLRTFQAVAHTGSVTAAAKQLHRAPSSITTRIHQLEEAFEQPLFLRIRNRLTLSPA
GERLLHQVDQLIAMFDRIQR
MMHESKQSDHISLGALDVALDTYLPTAIGKVREHFPKSRLFVRQAPSEILRDELVNKRL
DVVLNDGPINADGIISQYAYS
ERLVLVTDTHQPTITRSSELEGLDIYGFQRNCSYRLKLDEWLREGGLVSPNVIEMESYH
VILACVSGGTGAAWVPETYLN
TIKDRSAIKVHDLGETGNTDLYFSWRKNEDSPLLQALIENVA
>119_Erwinia sp. P6884 PtrR
MDLTQLEVFRAIAQEGSVTAAAGKLHRVPSNISTRLRQLEDELGVELFRREKLRLHITD
AGRTLLDYATRLLTLAEEAKE
RVSGQHPAGVFTLGAIESTAAVRLPPLIARYHQTWPAVELDLSTGPSGEMIEGLLEGKF
SAIFVDGPPRHPQLEGVAAFS
EELVLISSLNHSPVTQAAQISGATIYAFRANCAYRRRFENWFAREDAAPGKIFEMESYH
GILACVSAGAGLAMIPRSMLE
TMPGRESVQAWPLADGKGKLAIWLIWRKNGETANLRAMVSLLDAQ
>120_Erwinia endophytica PtrR
MELTQLEVFRAIAEEGSVTAAAERLHRVPSNISTRLRQLEEELGVALFSREKLRLHITDA
GRTLLDYAARILALTEEARE
RVSSQLPAGIFTLGAIESTAAVRLPPLIARYHQTWPEVELDLSTGPSGEMIDGLLAGRFS
AVFIDGPPKHPQLEGMAVFA
EEMVLITSLNHPPVRSAADIAGATIYAFRANCSYRRLFENWFAQDNAAPGKIFEMESYH
GIVACVSAGAGLAMIPASMLQ
NMPGRDSVKAWPLARPWGELHIWLVWRKGNLSANLQAIVNMLNAE
>121_Erwinia billingiae PtrR
MDLTQLEVFRAIAEEGSVTAAAERLHRVPSNISTRLRQLEDELGVMLFSREKLRLHITD
AGRTLLDYAIRILALTEEARE
RVSSQHPAGIFTLGAIESTAAVRLPPLIARYHQAWPGVELDLSTGPSGDMTDGLLSGR
FSAVFIDGPPKHPQLEGVAAFA
EEMVLISSLNHPSVSRAADINGSTIYAFRANCSYRRLFENWFALDNAAPGKIFEMESYH
GIVACVSAGAGLAMIPKSMLE
NMPGRDSVMAWPVADDRGLLDIWLIWRKGNGSANLKAMASMLEID
>122_Erwiniaceae bacterium CAU 1747
MDLTQLDVFRAIAEEGSITAAADRLHRVPSNVSTRLRQLEEELGCTLFHREKLRLYITDS
GRTLLDYARRILALADEAKL
SVSGQHPSGIFTLGAIESTAAVRLPPLIARYHQRWPQVELDLSTGPSGDMIDGIFSGRH
SAAFVDGPPRHPQLEGVPAFS

EELVLIGALNHPPVPDAASISGSTIYAFRANCSYRRLFENWFARENAVPGKIFEMESYH
GILACVSAGAGLALIPASMLE
SMPGRNSVQAWPLGGGMGNIDIWLVWRRGTHSTSLQAMI
>124_Sphingobium sp. AP50
MQISQLRAFCAVAQTGSITTAARVLNKVPSSITVRIRQLEEDLGRELFLRENKRMSLSPA
GRQLLDHAQQILDLTDGTRA
LMRDEETSGPLLVGALDVVLVDYMPHLIGMMRQRHRGINLTVRHQASEDLVAHVADG
SLDIALTEGPISSKALRSKVAFV
DEMLLVTELSHRDVHQPADLECGELYGFRHDCSFRFRMDRWLSAGDRLDMPVIEIES
YHTMLACVSAGMGAAWMLRSILK
TLPGHHQVKAHSLGEHGRTEIHFVWRDGHLS
>125_Yersiniaceae sp. PtrR
MDLTQLRMFCCVAETGSLAKAAEQLHRVPSNLTTRLRQLEQELGTDLFIREKQRLRLS
AMGHNFLGYAERILALSDEAMS
LTHAGEPAGQFALGSMESTAATRLPALLAAYHQQHPKVALSLVTGTSGDIIDSVRAGTL
AAALVDGPIDHDDLNACIAFR
ERLVVISPANRPLAAHGAGASLTLFVFGPSCSYRRKFYAWLKQAGWAVGASLEIHSYH
AMLACVASGAGIAILPESVLAQ
LP
>126_Pseudomonas sp. RIT-PI-AD PtrR
MEFSQLRIFQAVAEEGSIAKAALRLSRVPSNLSTRLRQLEAQLGVELFHRERQRLQLTP
AGRVLQRYAERLFALQGEAEA
ALKGDEPGGVFVLGSMYSTAAIHLPALLARYHHAYPGVNLQVRTCPSGELLDGLIGGR
LDAALLNGSGDYPELEGMPLFD
EDMLLITELGHPPVRSARDVAGRAVFTFRPRCSYRRLLEAWFAEARVAMGPLMEIESY
HSMLACVIAGSGVALMPASMLD
SLPGREAVARHRLAAPFDRATTWLMWRKGMLGANLRAWID
>127_Erwinia pyri PtrR
MDLTQLEVFRAIAEQGSVTAAAERLHRVPSNISTRLRQLEEELGVALFTREKLRLHITDA
GRTLLDYAGRILALTEEARQ
RVSSQQPAGVFTLGAIESTAAVRLPPLIARYHQAWPEVELDLSTGPSGDMIDGLLAGR
FSAVFVDGPPKQPQLEGIPAFA
EEMVLISSLNHPPVARAADIAGATIYAFRANCSYRRLFENWFARDNAAPGKIFEMESYH
GIVACVSAGAGLAMIPESMLQ
NMPGRESVLAWPLAEQRGHLNIWLVWRKGNASANLRAMMNMLEAQ
>128_Erwinia mallotivora PtrR
MDLTQLTVFRAIAEQGSVTAAAVQLHRVPSNISTRLRQLEDELGTELFTREKLRLHITDA
GRTLLDYACRILALVEEAQQ
QVSGSQHAGIFTLGAIESTAAVHLPPLIARYHQAWPQVELDLSTGPSGEMIDGLLAGRF
SAVFIDRPPRHPELEGVAVFP
EEMVVISSLNHPPVTSADNIAGATIYAFRANCSYRRLFENWFSRAGSAPGKIFEMESYH
GIVACVSAGAGLAMIPLSMLK
SMPGRDSVRTWPVADGMGEIAIWLVWRKGSAPANLRAMLD
>129_Rouxiella aceris PtrR

MDLTQLEVFATVAAEGSVTAAAEKLHRVPSNISTRIRQLEEELGTLLFHREKLRLHISDQ
GKTFLSYALRILALAEEAKQ
SVSGQRPMGIFTLGSIESTAAVRIPPVLARYHQAYPSVELDLSTGPSGDMIEGMLSGRF
SAAFVDGPPRHPELDGVAVFD
EEMVLIAALNHAPIQRAQDISGATIYAFRANCAYRRLFENWFAEDQAVPGKIYEMESYH
GILACVTAGAGLALIPASMLE
NMPGKTGVHAYPLANEMGKLQIWLMWRKGQTSANLQAMVS
>130_Pseudomonas sp. dw_358
MQLSQLRAFCAVARTGSVTVAAKELHRVPSGVTLRIQQLEADLDCQLFVREKQRLTLA
PAGRALLERALQILELSDGAKA
LVREEEIGGHLTLGALDVGLVAFMPQLVGRFRQRHPQVSMDIRSDVSEVLMSQVAEG
LLDIALTDGPVQSPALDSRYAFS
DDLLLITERAHGPVRSAGDLQCREVYGFRQNCSFRLRLDRWLQATGGERLPLIEMESY
HTMLACVGAGAGAAWVPRAMLD
TLPGRAAVRVHSLGAAGRTDLYFVWRS
>131_Pelagibius sp. Alg239-R121
MDLSDLKIFTAVVEEGGITRAAERLHRVQSNVTTRIRQLEDELGVPLFIRQGKRLHLAPA
GQTLLDYAKRLLLLAEEARE
AVHGDVPRGLFRLGSMESTAAVRLPGPLAAYNDLYPKVELELRTGNPTQLASALLSGE
LDAALMAEPIADAKFESVLAFE
EELVIVTAQDHPTIDKPDAVPRTILVFEHGCPHRRQLERWYAARDETPQRIIELRSYHA
MLGCVLAGMGAALLPRSVLGT
FPESKRLRIHDLRAGECDLNTLLVWRKDVRSPKVKALAD
>132_Pseudomonas alkylphenolica
MELSQLRIFQAVAEAGSITRAAERMHRVPSNLSTRLRQLEEQLGVELFRRERQRLQLS
PAGKVLLDYAGRMLALRDEAVA
AVQGGQPAGDFALGTMYSTAAIHLPALLARYHKTYPAVNLQVLPAPSGELLEGLLSGR
LDAALVDGPLALAGLDGVPLCE
ETLVLITEPDHPPVHSARDVAGRAVFTFRQGCAYRMRLEAWYAQDHAAMGRVMEIES
YPSMLACVIAGSGVALMAQSMLD
SLPGRERVAAHRLQSPFDLATTWLMWRKGMRGANLSAWID
>133_Pantoea sp. IMH
MDLTQLKMFCLVAETGSLVRAAEVLHRVPSNLTTRLRQLEHELGTDLFIREKQRLRLSP
MGHNFLCYAQRILALSDEAMR
MAQSGEPGGNFALGSMESTAATHLPGLLAAYHQRYPDVALSLVTGTSTEIIKDVREGK
LAAALADGPATADELNSCVAFR
EKLILISSRDHLPIAEPADAAGDTLFTFRSGCAWRQRLESWFRSGGAQPGPFMEIQSY
HAMLACVASGAGVALMPESVLN
QLPASSILRHALPESIADAATWLVWRRDGFTPNVEAL
>134_Bordetella genomosp. 9
MDLVQLEIFCAVAQHQSIAAAAEQIHRVPSNLTTRLKQLEADLGVDLFIRENNRLRLSAT
GRNFLGYAKNILSLVDQARS
AVSGEEPAGSFPLGSLESTAAVRIPGILARFNARHPDVELELSTGPSGSMIDGVLEGRL
VAAFVDGPIKHPAIDGIPVFD

EEMVIIAPAQHAPVRRGRDAAGDTIYVFRDNCSYRHHFEKWFAEDKAIPGKVREIESYH
TMLACVSAGGGLAVIPRAMLE
SMPGYLSVAAWPMHGRFSVLQTWLVWRRDTASRALDAF
>135_Pseudomonas sp. PA15(2017)
LSLNKIKFFCGVVENETIAAAAKIFHTVPSNLSLRIQDLERDLGVDLFYRENKKLLLTPQG
RLFYKKVKPLLSELSELEN
SYKAGSVFSHLRLGVPDFALMSIISSQVESIRRDITGIKIEIQSQDSYCLEQMLMSGDLDL
ILMEGPVVHPLLDAKFFCD
QDFFLITPECFAGSELHALSELDYYSFNGSCSISALTLSWMENNGITPSYTLQTDNYAT
RVNSVRGGEGFSFLPRSIIES
NIYDLSKIHVRDLHGQIRQSLYFAWRKK
>136_Pseudomonas sp. PtrR
MEFSQLRIFQAVAEEGSITRAAQRLNRVPSNLSTRLKQLEEQLGTPLFVRERQRLQLSP
AGKVLLDYTLRLMALRDEAEA
AIKGGEPAGDFVLGTMYSTAAIHLPAMLVRYHRAYPAVNLQVQAGTNGELIDGLLNGR
LDMALVNGPPEFSALDGIPLFE
ERMVVISDAARGPIKSAEDIEGLSVFTFRSGCAYRKRLETWYASQHVAMGRVMEIESY
QSMLACVIAGAGVALIPQSMLD
SLPGRENVLVHELDSPFDRATTWLMWRRGRVGANLNAWID
>137_Erwinia unclassified sp.
MDLTQLKMFCSVAETGSLARAAEQLHRVPSNLTTRLRQLEQELGADLFIREKQRLRLS
PVGHNFLCYAQRILALSEEAMS
MAHNGEPGGNFALGSMESTAATRLPVLLAAYHQRYPGVALSLITGTSGEVISRVREGT
LAAALADGPITADELNSCIVFH
EKMVLISGLGHTPIHHPADAVGETLFAFRPSCSYRLRFENWFRSSGIQPGSFMEIQSYH
AMLACVASGAGLALLPESVLG
QLPARAGVQRHSLPADVSETDTWLVWRRDAFTPNVKAL
>138_Erwinia sp. PtrR
MDLTQLEVFRAIAEEGSVTAAAGKLHRVPSNISTRLRQLEDELGTVLFSREKLRLHITDA
GRTLLDYANRILALTEEARL
SVSSQHPAGVFTLGAIESTAAVRLPPLIARYHQTWPEVELDLSTGPSGAMTDGLLAGE
FSAVFIDGPPKHPQLEGMPVFN
EEMVLISSLNHPPIARAREISGATIYAFRANCSYRRLFENWFARDNASPGKIFEMESYH
GIVACVSAGAGLALIPKSMLA
SMPGKESVRAWPLAGDQGKLAIWLVWRKGGGSANLRAMVKMLEA
>139_Erwinia sp. S38
MDLTQLDVFRAIAEEGSISAAADRLHRVPSNISTRLRQLEEELGCELFRREKLRLHITDS
GRTLLDYARRILALADEAKL
SISGQHPSGVFTLGAIESTAEVRLPPLIARYHQQWPQVELDLSTGPSGDMIDGIFSGRY
SAAFVDGPPKHPQLEGVPAFS
EELVLIGALNHPPVPDAASISGSTIYAFRANCSYRRLFENWFARENAVPGKIFEMESYH
GILACVSAGAGLALIPASMLE
SMPGRDSVQAWPLGGGMGNIEIWLVWRKGTHSANLKAMTE
>140_Pseudomonas aeruginosa PtrR

MEFGQLRIFQAVAEEGSIARAAERLHRVPSNLSTRLRQLEEQLGVDLFLRERQRLQLS
PAGKVLLDYAARLFALQEEARA
AVQGGEPVGDFALGSMYSTAAIHLPPRLAEYHRRYPAVNLQLQTAPSGELVESLLGGR
LDAVLVDGPLDCDGLEGLPMFE
ERMVLVTENGHPPVRGPEDVAGSAVIAFRPRCSYRLLLESWFASARVSMGRVMEIES
YHSMLACVVAGGGVALMPVSMLQ
SLPGRESVAVHALAEPFARASTWLVWRKGMVGANLKAWI
>141_Rouxiella sp. Mn2063
MDLTQLEVFKAVAEEGSITAAADKLHRVPSNISTRIKQLEDELGVNLFQREKLRLYISDA
GRTFLEYSRRILDLAEEAKQ
SVSGQQPSGIFTLGSIESNAAVRLPSLLARYHQRFPQVELDLSTGPSGTMVEGLLSGQ
YNAAFFDGPLRHPEFDGIPVYD
EQLVLIASLNHPPVTRAKAISGSTIYAFRTNCSYRRIFENWFREDNAAPGKIFEMESYHG
IVACVTAGAGLALMPISMLE
SMPGQTGVKAYPLSGTMGQVKIWLVWRKGMATANLNAMV
>142_Pseudomonas sp. AL 58
MELSQLRIFQAVAETGSVTRAAEQLHRVPSNLSTRLRQLEEQLGVELFRRERQRLQLS
PAGKVLLDYAGRLLALHDEAFA
AVQGGQPAGSFVLGTMYSTAAIHLPALLARYHRAYPAVNLQVQAAPSGELLEGLLSGR
LDAALVDGPLSLAGLDGVPLCD
ETLVLITEADHPPVRSARDVAGRAVFTFRQGCSYRMRLEAWYAHDHAAMGRVMEIES
YPSMLACVIGGAGVALMSQSMLE
SLPGRESVAVHRLQSPFDEATTWLMWRK
>143_Pseudomonas fluorescens PtrR
MEFSQLRIFQAVAEEGSITRAAERLHRVPSNLSTRLKQLEEQLGVELFVRERQRLQLSP
AGKVLLDYTAKLFALRDQASA
AVMGGQPAGDFVLGTMYSTAAIHLPELLARYHQQYPAVNLQVQAAPSGELLEGLLTG
RLDAALVDGPLELAGLDGVPLCD
ERLVLITEPEHPPVRTALDVAGRAVFTFRQGCSYRMRLEAWFAHYHAAMGRPMEIES
YQGMLACVIAGSGVALMAESMLA
SLPGRERVAMHPLAEPFVGATTWLMWRKGMVGANLNAWID
>144_Metapseudomonas otitidis PtrR
MEFSQLRIFQAVAESGSIARAAERLNRVPSNLSTRLRQLEEQLGVDLFLRERQRLQLSA
SGKVLLGYAERLFALQAEAEA
AIKVGEPAGDFLIGSMYSTAAIHLPPMLARYHGAWPAVNLQVQTQPSGELVEGLVAGR
LDAALVDGPLEYADLDGIPLFD
EALVLITALGHGPVTSAADVAGAAVYTFRSRCSYRKRLERWFAESRVAMGRVMEIESY
HSMLACVVAGGGVALMPRSMLQ
SMPGREAVTIHPLAEPFATATTWLMWRKGMLGANLKAWIE
>145_Erwinia typographi PtrR
MDLTQLEVFRAIAQEGSVTAAAARLHRVPSNISTRLRQLEDELGVALFNREKLRLHITD
AGRTLQDYAGRILALTEEARE
RVSGQHPTGIFTLGAIESTAAVRLPPLIARYHQAWPEVELDLSTGPSGEMTEGLLAGRF
SAVFIDGPPKHPQLEGVPAFA

EEMVLISSLNHPAVSCAADINGATIYAFRANCSYRRLFENWFAVDNAAPGKIFEMESYH
GIVACVSAGAGLAMIPKSMLQ
SMPGRDSVLAWPLAEARGHLHIWLVWRK
>146_Pantoea sp. BAV 3049 PtrR
MDLTQLKMFCSVAETGSLARAAELLHRVPSNLTTRLRQLEQELGTDLFIREKQRLRLSP
VGHNFLCYAQRILALSEEALS
MTHSGEPGGNFALGSMESTAATRLPALLSAYHQQHPTVTLSLITGTSGEIIERVREGTL
SAALADGPVNVNELNSCKAFD
ERMVLISSLSHAPIMQPSDAAGDTLFAFKPSCSYRLRFENWFRSAGVQPGTFMEIQSY
HAMLACVASGAGLALLPESVLA
QLPAAERIQRHKIPAEVSETATFLVWRRDAFTPNIKAL
>147_Pseudomonas unclassified sp. PtrR
MDLVQLEIFCAVATHKSIAAAAQAMHRVPSNLTTRIKQLEADLGADLFIRENNRLRLSLA
GHDFLVYATRILGLVEEARV
VVSGKEPTGRFPIGSLESTAAVRIPRLLARYHQQYPKVELDLSTGPSGEMIDGVIAGELI
AAFVDGPIRHPALEGVPVFD
EEMLVVAPSHHAPIQRGQDADGEMIYVFRDNCSYRHHFERWFAADGAAPGAIRELES
YHSMLACVSAGGGLAVMPRSMLS
NMPGSASVSAWPMAGDFARLNTWLIWRSDSRSRSLELF
>148_Pseudomonas mangiferae PtrR
MEFSQLRIFQAVAEEGSIAKAALRLNRVPSNLSTRLRQLEERLGVALFLRERQRLQLSP
AGKVLQDYASRLFALQDEAEA
ALKGGEPAGTFVIGSMHSTAAIHLPAMLAHYHRRYPAVNLQVHTLPSGELVDGLLAGR
LDAALVDGPEVSSELEGIALFD
EELLLITEAGHAPVRSARDVAGQPLFTFRPRCSYRRRLENWFAAQRATLGPVMEIESY
HSMLACVIAGGGVALMPRSMLD
SLPARDAVSVHPLAAPFGQATTWLMWRKGMAGANLRAWSDGL
>149_Paraburkholderia tropica PtrR
MELAQLQIFREIADAGSVQAAAQRLHRVPSNLTTRLRQLEEELGAELFIREKMRLRLSA
TGAAFLEYARRILDLVDEAKL
SVAGDEPRGALPLGSLESTAAVHIPPVLAAFHRRYADVTLDLSTGPSGELIDRVIEGELA
ATFVDGPVAHPALTGVPVFD
DELILISAANRAPIKRARDVNGANLYAFRSNCSYRRTFERWFEEDGAAPGRIFELESYH
GMLACVSAGAGLALMPASLLE
SLPNRSAVKAHRLPAAYRKVKIYLVWRRDLRSPAVQRLIEMLEA
>150_Enterobacter sp. Ap-1006 PtrR
MDLTQLEMFNAVAETGSISAAAQQVHRVPSNLTTRIKQLEADLGVALFIRENQRLKLSP
SGHSFLAYSKRILALVDEARM
VVSGDEPQGPLALGSLESTAAVRIPGVLANFNQRYPKIHLSLTTGPSGDQIDGVLEGRL
AAAFVDGPVLHPSLEGVAVYR
EEMTIVAPADHAPILSGADVNGESIYAFRANCSYRRHFESWFHDSQAMPGKIHEMESY
HGMLACVIAGAGLALMPRSMLE
SMPGSHQVSTWPLPEDKRYLDTWLLWRRDAKTRQLDAFI
>151_Duffyella sp. PtrR

MQLSQLEMFRAVAATGSISAAAEAVHRVPSNVTTRIKQLEAELGVALFIRENQRLRLSP
AGRHFLDYNNRILDLVDEARL
AVSGDRPAGLFALGSLESTAAVRIPPLLARYHHHYPQVELALSTGPSGDMLDRVLEGS
LEAAFVDGPILHPVLEGVPVFR
EEMVLVATQQHPAIACAEDVNGENLFAFRDNCSYRRHFESWFRDGNAMPGKIYPVES
YHGMLACVTAGSGIALMPRSMLE
SMPGSGTVSAWPLAENYRYLDTWLVWRRGPRSINLNEFI
>152_Zestomonas insulae PtrR
MELSQLRIFQAVAEEGSIARAATRLNRVPSNLSTRLRQLEEALGVDLFLRERQRLQLSA
AGKTLLGYAERIFALQAEAEA
AVKGDEPVGDFVLGSMYSTAAIHLPAMLVRYHQTYPAVNLQVQTQISGELMDGLLAGR
LDAALVDGPLQHSELDGLPLFE
EELQLITPIGHAPVHSAQDVAGEAVFTFRQRCSYRRRLEAWFAASRVATGPVMEIESY
HSMLACVVAGGGVALMSRSMLD
SLPGRESVAAYTLAPPFNSTTTWLMWRKGRFGANLRAWID
>153_Erwinia sp. B116 PtrR
MDLTQLEVFRTVAEEGSITAAAGKLHRVPSNVSTRLRQLEEELGCELFRREKLRLHITD
SGRTLLDYARRLQALADEARA
CVSGQQPGGIFTLGSIESTAAVRLPPLISRYHQRWPQVELDLTTGPSGAMIDGVLSGR
YSAAFVDGPPRHPQLEGVAVFA
EELVLISALHHPPVPDAASISGATVYAFRANCSYRRRFENWFARDGAVPGKIFEMESY
HGIVACVSAGAGLALVPASMLA
SMPGRDSVRAWPLADGQGAAEIWLVWLKGSQS
>154_Pantoea sp. MBD-2R PtrR
MDLTQLEVFRAIAEEGSVTAAAGKLHRVPSNISTRLRQLEEELGTQLFSREKLRLHITDA
GRTLLDYAHRILALTEEARL
QVSSQHPTGVFTLGAIESTAAVRLPPLIARYHQTWPDVELDLSTGPSGEMIDGLLAGEY
SAVFIDGPPKHPQLEGMPVFA
EEMVVISSLNHPPVTSATTISGATIYAFRANCSYRRLFENWFARENAAPGKIFEMESYH
GILACVSAGAGLAMVPRSMLE
NMPGKDSVSAWPLAGQMGLINIWLVWRKGSRSANLRAMTD
>155_Burkholderia sp. PtrR
MDLIQLEMFQAVARHGSIAAAAQAVHRVPSNLTTRIKQLEEELGVDLFTRERNRLQLSQ
PGRVFLDYASRILGLVSEARA
VTAGQQPAGRFALGALESTAAVRIPGILAAYNQRFPQVALELATGPSGQIIDALLGGDLI
AAFVDGPLDHPELTGFPVFD
EQMVVVAPSNHAPIRRGRDADGDALYVFRKNCSYRRHLERWLAHDGAVHGEIRELES
YHGMLACVSAGGGLAIMPLSMLE
SMPGSQTVRAWPMGKEFGLLRTWLVWRKETISKALEAFTQLVKAS
>156_Carnimonas bestiolae
MDINQIRAFQAVAQSGSVTAAAQQLHRAPSSITTRIHQLEEEFDQQLFLRIKNRLVISSA
GRRLLQYTDQLLPLFDRAQR
AMKEYNRGHHLSLGALDVSLETYLPQAIGSVRERISEARIYVRQAPSEVLSDELLNGTL
DAVLNDGPVDAPGIGNEYVFS

EQLVLVTDKDHPKVTSSMDLEGVDIYGFQKNCSYRLRFDQWLQQGGLSSPNVIEMES
YEVMLACISGGSGAAWVPAAYLD
RVHHSSSVKIHDLGSIGITDLYFSWREEEHSDVLRQLIEEV
>157_Mixta sp. PtrR
MDLVQLRMFCSVAETGSVARAAEQLHRVPSNLTTRLRQLEQEIGVDLFIREKQRLRLS
PMGHNFLCYAQRILALSEEALS
MARAGEPGGNFALGSMESTAATRLPNLLAAYHQRYPTVALSLITGTSGEIIDRVREGTL
AAALVDGPANYDELNGCIAFR
ERMTLISSVDHTPIRSAADASGDTVFAFRTSCSYRLLLEEWYRDSGVQPHNVMEIQSY
HAMMACVASGAGVALIPESVLA
QLPARERVQAHRLPDRFRDTATWLIWRRDAFTPNVEAL
>158_Minicystis sp.
MELSDIEVFQTVVAEGGVIAAARRLHRVPSNVTTRVRKLEEELGVPLFLREKNRLRLAP
AGRLLLPYAERLLALSEEARQ
ALRDPTPRGPLRLGSMESTAAVRLPRPLAAYHKRFPEVAVELRTGPSQKLVAAVLAGE
LDAALVADPGTDPRLTRLPVFR
EELVLVTEASRRDPVTARDLAGATLLAFADGCAYRRRLEAWLADASVVPERTAEIASY
HAMLGCVVAGMGVALVPRSVLQ
TLPARSRTRAHPLPKRWRWAETILIWRREAGAARIEALAEVMRSE
>159_Shimwellia pseudoproteus PtrR
MDLTQLEMFTAVAETGSISAAARQVHRVPSNLTTRIRQLEEELGKDLFIRENQRLRLSP
DGHNFLIYSRQILGLVDEARR
VISGDETRGTLALGALESTAAVRIPSILAAFNQRYDGIQLDLSTGASGTMLDGILEGTLS
AAFVDGPVLHPVLEGVPVFD
EEMMIIAPISHQPITRARDVSGASLYAFRANCSYRRHFEGWFLADQAMPGKIHEMESY
HGMLACVIAGAGLAMMPRSMLE
SMPGSHQISAWPLAEPWRWLTTWLVWRRGARTPQLDAFID
>160_Erwinia endophytica PtrR
MDISQLKVFCSVAREGSISQAAKVLHRVPSNISTRLKQLENELGTDLFIREKLRLHLSEQ
GKIFLEYAEKVLMTLDEARE
AVVSSEPKGSFSLGAILSTAAVRAPDILAAYHQRYPHVELDFSTGASDDMVNGLLEGRL
TAAFIDGPIPHPLLEGMPLWT
EQMMVVAAKNHPPITCAKDVAGAQLYAFRRVCSYRRYFESWFEAENVRPGKISEMES
YHGMLGCVSAGAGLALAPQSILA
TIPGAQHNLSIWPLKGNFQSTQIWLWWKKNTRSANLQAL
>161_Ewingella americana PtrR
MDLTQLRMFCSVAETGSLARAAEQLHRVPSNLTTRLRQLEIELGCDLFIREKQRIRLSA
MGHNFLNYAQRILALSDEAMS
ITHAGEPAGGFALGSMESTAATRLPSLLAAYHQLYPQVSLSLNTATSGETIDGVREGKL
AAALVDGPIDYDDINGCIAFR
ERLVIISPQGESPFDTPPGRSGHTVFAFRATCSYRTRLQSWIKHTGLAVSNIMEIQSYH
AMVACVASGAGIAMVPRSVLD
QLPGHERVQAHEIEDGFANTATWLIWRRDAFTPNVRALKELI
>162_Citrobacter amalonaticus PtrR

MDLTQLEMFNAVAETGSITQAAAKVHRVPSNLTTRIRQLEADLGVDLFIRENQRLRLSP
AGHNFLRYSQQILALVDEARM
VVAGDEPQGLFSLGALESTAAVRIPATLARYNQRYPKIQFSLATGPSGNMLDGVLEGH
LNAAFVDGPITHPGLEGMPVYQ
EEMMVVTPHGHAPVTRACEVNGSNIYAFRANCSYRRHFENWFHADRAMPGTIHEME
SYHGMLACVIAGAGIALIPRSMLE
SMPGHQQVDAWPLSGNWRWLTTWLVWRRGAKMRQLEAFIE
>163_Rhizobiaceae bacterium
LDLSTLEIFRQVAREGSISSAAKNLNRVQSNVSTRIKQIEDQIGVSLFERGRRGMALTDS
GHTLLGYANELLTLSEQAVE
AVGIVRPAGVLRIGAMESTAASRLPVILTRFHDENPNVEVHVRTDTAGAIIGHLMSGQID
VAFVAEPVELRNIIKQPVFR
ENLILITPKSFPGSSIRDLTGKTMIAFEEGCAYRRYLNQWLLENGIVPEKTISIGSYLGMF
ACISAGTGFGVVPESVL
>164_Shimwellia blattae PtrR
MDFTQLEMFTAVAETGSISAAARQVHRVPSNLTTRIRQLEEELGKDLFIRESQRLRLSP
DGHNFLVYSRQILALAAEARR
VLSGDETKGALTLGSLESTAAVRIPSILARFNQRYSGIQLDLATGSSGTMLEGVLEGTL
GAAFVDGPVLHPALEGVPVFD
EEMMLIAPASHAPVSRARDVSGASLYAFRSNCSYRRHFESWFLADQALPGKIHEMES
YHGMLACVIAGAGLAMMPRSMLE
SMPGSHQVSAWPLAEPWRWLTTWLIWRRGARTPQLDALIE
>166_Dryocola clanedunensis PtrR
MDLTQLQMFNAVAETGSIQAAAQLVHRVPSNLTTRIKQLEADLGVQLFIRESQRLKLAP
AGYSFLEYSRRILALVDEARM
VVAGDEPQGVLALGSLESTAAVRIPSLLAAFNQRYPKIQLSLSTGPSGDQIDGVLEGRL
SAAFVDGPLLHPSLEGQAVYK
EEMVIVAPVDHPPIETPRQVNGANIYAFRANCSYRRHFENWFHQDGARPGKIHEMESY
HGMLACVVAGAGLALMPRSMLE
SMPASHQASAWPLSSDWRYLNTWLIWRRGARTRQLDAFM
>167_Pseudomonas sp. JV551A1 PtrR
MDLTQLEIFRAVAEEGSVTAAAQRLHRVPSNVTTRLKQLESELGVELFIRQRLRLYLSP
TGAVFLDYSRKILDLVSEARG
VIKGGEPHGMFAIGSLESTAAVRMPAVLASFHQHYPAVQLDLTTGSSAEMIIGVSEGRL
AAAFIDGPVGHLALDGQKVFE
EDMLVIAPRKHAPIRHPRDVAGETVYAFRSNCSYRRHFENWFASARVRPARVFEMES
YHGMLACVSAGAGIAMFPRSMLS
AMQGAEMVESFPLSGSFSKVEIWLAWRKGFMHPNVNALLQLLSPD
>168_Citrobacter freundii PtrR
MDLTQLEMFNAVAETGSITQAAAKVHRVPSNLTTRIRQLEADLGVDLFIRENQRLRLSP
SGHNFLRYSQRILALVEEARM
VVAGDEPQGLFSLGSLESTAAVRIPATLAEYNQRYPKIQFALATGPSGTMLDGVLDGTL
NAAFVDGPITHPGLEGMPVYR

EEMMIVTPNGHSPIQRASDVNGCSIYAFRANCSYRRHFESWFHADRAMPGTIHEMES
YHGMLACVIAGAGIALIPRSMLE
SMPGHHQVNAWPLSENWRWLNTWLVWRRGAMTRQLEAFIEVLN
>169_Bradyrhizobium sp. JYMT SZCCT0180
MDLSDLKIFSAVVREGGVTRAAEHLHRVQSNVTTRIRQLEDDIGVALFIREGKRLHLAP
AGQVLLDYADRLLALADEARN
AVQDPRPRGIFRLGAMESTAAVRLPGPVSEYHRLYPEVEFELRTGNPPTLSKAILAGEL
DAALVTLPIADALFEKVAVFE
EEPVIVSAAGMPAIGQGKNKGKTEYFPRTIIAFEHGCPHRKRLEDWYALRDQMPERTIE
LGSYHAMLGCVAAGMGVALLP
KSVLTTFPESKRLRVHRLPPGENRADTYLIWRKGAGSPKIQALRDVLGA
>170_Paramixta manurensis PtrR
MDLIQLRMFCAVAATGSIVRAADQLHRVPSNLTTRLRQLEQELGTDLFIREKQRLRLSP
MGHNFLGYAQRILSLSEEALN
MGRSGDPAGNFALGSMESTAATRLPALLATYHQRFPAVALSLITGTSGEIIERVREGTL
AAALVDGPINYDELNGCIAFR
EQMVLISSPDRAPIANATDASGDTLFAFRSSCSYRLRIEAWFRENGAQPGNVMEIQSY
HAMLACVASGAGLAMIPQSVLA
QMPDRMRVQVHPLPPAFRDTATWLIWRRDAFTPNIEAL
>171_Pseudomonas gingeri PtrR
MEFSQLRIFQAVAEEGSITRAAERLHRVPSNLSTRLKQLEEQLGVELFLRERQRLQLSP
AGKVLLDYCSKLFLLHDEAQA
AVQGGQPGGDFVLGTMYSTAAIHLPSLLSRYHRAYPAVNLQVQSSISSELFEGVLSGR
LDAALVDGPLQLAGMEGIPLFE
ESLVMIGAADHPPILSARDVEGRPVFTFRPGCSYRVRLEAWYAHYQAAMGRAIEIESY
PGMLACVTAGSGVAMMSRSMLA
SLPGRENVSVHPMAAPFDSATTWLMWRKGMKGANLTAWIE
>172_Pseudomonas akapageensis PtrR
MEFSQLRIFQAVAQEGSITKAAERLHRVPSNLSTRLKQLEDQLGVELFLRERQRLQLSP
AGKVLLDYAVRLFALHDEAHA
AVQGGQPAGDFVLGTMYSTAAIHLPALLARYHKAYPAVNLQVQAGPSGELLEGLLSGR
LDAALVDGPLQLAGLDGVPLCN
EELVLISEADHPPVRSARDVEGRAVFTFRQGCAYRMRLEAWYAHDHAAMGRVMEIES
YPSMLACVIAGSGVALMSASMLA
SLPGRESVAVHPLQPPFDQATTWLMWRKSMVGANLNAWIE
>173_Pantoea alhagi PtrR
MDLVQLRMFCLVAETGSVARAAEQLHRVPSNLTTRLRQLEQEIGVDLFIREKQRLRLS
PMGHNFLSYAQKILTLSEEALS
MARTGEPGGNFALGSMESTAATRLPNLLAAYHQRYPAVALSLITGTSGEIIDRVREGTL
AAALVDGPVNYEELNGCIAFR
EQMILISSAGHGPIRSAVDASGDTVFAFRSSCSYRLRLEEWFREDGVQPHNVMEIQSY
HAMMACVASGAGVALIPQSVLA
QMPARERVQAHVLPPKFSDTATWLIWRRDAFTPNVEAL
>174_Pseudomonas piscis PtrR

MEFSQLRIFQAVAEEGSITRAAERLHRVPSNLSTRLKQLEDQLGVELFLRERQRLQLSP
AGKVLLDYAARLFALHDEAQA
AVLGGQPAGDFVLGTMYSTAAIHLPGLLARYHRTYPAVNLHVQSGPTGELLEGLLAGR
LDAALVDGPMELAGLDGVPLCR
ENLVLISEVDHPPVHTALDVEGRAVFTFRQGCSYRMRLESWFAHYHAAMGRAMEIES
YQSMLACVIAGSGIALMTRSMLD
SLPGRESVAVHPLAEPFAHSTTWLMWRKGMVGANLNAWIE
>175_Pantoea sp. B65 PtrR
MNLSQLEMFSAVAETGTISQAARKVHRVPSNVTTRVRQLEADLGVELFIRENQRLRLS
PAGHSLLQYSQQILSLVAEARQ
VVTGQQPQGIFTLGSLESTAAVRIPGILGQYHQRYPMIQLDLSTGPSGNMLNAVLAGEL
SAAFIDGPVMHPVIEGMPVFR
EEMVIIVAAGYPEIKRAQAVSGHNIYAFRENCSYRRHFENWFLDDQATPGKIYEMESYH
GMLACVIAGAGLAMLPRSMLE
SMPGHEQVRVSIPDENWRWLNTWLIWRRGAKTPQLEALIRLLPAD
>176_Gammaproteobacteria bacterium
MNLDDLTIFRTVVHEGGITRAAAKLHRVQSNITTRVRQLEAELGVDLFVREGRRLTLAP
AGRVLLNYTERLLALADEARL
AVAGGSPQGSLRLGSMESTAAARLPRVLADYHARYLDVQLELRTGPSTPLVADVIDGR
LDAALVSAPIDDPRLHCLPVFE
EELLLITPATQPAVGDPRDLQVRTLLTFAPGCAYRRHLEAWLAQAGVVPERVVELSSY
HAMFGCAAAGMGIALAPRSLVE
LMPVDGVSLHSLP
>177_Sodalis (in: enterobacteria) unclassified
MNSASLNVFKAVADTGSIAAAARRLHCVSSNVTTRLKQLEAGLEVSLFIREKNRLYITPE
GEHLLRYANRILALIDEAQT
SLRDKNPVGPLRIGAMETTAATRLPPLLAQFHRHMPSVELHLNTSASGPLTAQVLNSE
LDVAFVAENDGIRHDLLNSAVL
CRETLVLVTAAGHPDVLTGADLQVTNPLAFRQGCNYRLRLEHWLWEQGISEAAPQEF
GSFQAILGCVAAGMGIALLPENV
VCQYEQGFSIRSHGITPAIGEVDTLMIWLKSRETSRVLRCFKDFV
>178_Pantoea cypripedii PtrR
MDLVQLRMFCSVAETGSLVRAAEQLHRVPSNLTTRLRQLEEELGVDLFIREKQRIRLSP
MGHNFLNYAQRILALSDEALS
MARAGEPGGNFALGSMESTAATRLPGLLAAYHQRFPSVALSLITGTSGEIIDKVREGTL
AAALVDGPVTHDDLNGCIAFR
EEMVLITSLEHKPIINARDTQGDTLFAFRNSCSYRVKLESWYRDSQTAPDNVMEIQSYH
AMMACVAGGAGVAMIPASVLN
QLPGKVRVQEHALPPAYRDTATWLMWRRDAFTPNVEAL
>179_Marinobacterium aestuarii
MDISDLQVFKAVVDSGGISRAAEQLHRVPSNVTARIQKLEQALQQPLFLREKNRLKITP
AGKRLLVYAEQMLRLRQQAID
DLTDPEPGGTLRIGSMESTSAAHLPPILSAYHREFSLVDIELRTGASGTLVDLVLQGELD
VALAADPVPDDRLCMHPCFE

EQLLIVKPAALSHSCGAAELPEPLSVVTFTQSCSYRNRIQRWLDGDNRRPDRTIEIPSH
HTMLACVLAGMGIALVPRAVL
ALHPQFSELATEDPGADISLATTWLIWRKDSTLPSITAFRDTI
>180_Winslowiella iniecta PtrR
MNLSQLEMFTAVAETGSISAAAKQVHRVPSNVTTRIRQLETDLGVELFIRESHRLRLSP
AGHSLLQYSKQILSLVSEARQ
AVTGEAPQGTFTLGALESTAAVRIPGLLAEYNRNYPRILLDLSTGPSGTMLDGVLEGKL
SAAFIDGPVTHPAIDGMPVFD
EEMVIIGPAKQKPFTRAREVSGVNIYAFRANCSYRRHFENWFHADQATPGKIYEMESY
HGMLACVIAGAGLAMLPRSMLN
SMPGNEQVVVCVPEEPWRWLSTWLIWRRGVKTPQLEVMIE
>181_Pantoea vagans PtrR
MDLVQLRMFCSVAETGSLARAAEQLHRVPSNLTTRLRQLEEELGVDLFIREKQRIRLSP
MGHNFLNYAQRILALSEEAMS
MTRTGEPAGNFALGSMESTAATRLPGLLAAYHQRFPKVALSLITGTSGEIIDRVREGTL
AAALVDGPVSHDDLNGCIAFR
EEMVLITGLDHAPITTARDVQDDTLFAFRNSCSYRVKLESWYRDSQTAPDNVMEIQSY
HAMLACVAGGAGVAMIPASVLK
QMADRSRVQAHTLPAAYRDTATWLMWRRDAFTPNVEAL
>182_Prodigiosinella aquatilis
MNSVSLNLFKAVADTGSVSAAAKRLHCVSSNVTTRLKQLEASLDVSLFIREKNRLYITP
EGEHLLTYANKILGLIEEAAS
SLRDKKPVGQLRIGAMETTAATRLPSLLSAFSRLMPFVELSLKTSPTLFLVKQVLESELD
IGFVADNGMLNHEQLNSAIL
CRETLVLVTSADHPNVMSGADLRVHKPLAFHQGCNYRSRLVLWLQEQGIDNAVPQEF
GSFQAILGCVAAGMGIAMLPENV
VKQYEQSLDIRCHSITPAIGEVDTLMIWLKSREASLVVRRFKDFVMQD
>183_Pantoea sp. PtrR
MDLVQLRMFCSVAETGSLARAAEQLHRVPSNLTTRLRHLEEEIGVDLFIREKQRIRLSP
MGHNFLNYAQRILTLSDEALS
MARAGEPAGNFALGSMESTAATRLPGLLAAYHQRFPKVALSLITSTSGEIIDKVREGTL
AAALVDGPAPYDDLNGCIAFR
ETLTLITSLDHAPVESARDAQNDTLFAFRNSCSYRVKLEAWYRDSGVAPGSVMEIQSY
HAMMACVAGGAGVAMIPESVLA
QMPERARVQAHELPPAYRDTATWLMWRRDAFTPNVEAL
>184_Pseudomonas sp. PtrR
MDLTQLELFKAVAEEGSISAAAQRLHRVPSNLTTRIKQLEQELGVDLFIREKLRLRLSAT
GWTFLDYARRILDLVEEAKQ
AASGSEPQGGFTLGSLESTAAVRIPSLLAAFHQRYPKVELDFSTGPSGDMIEGVLSGR
LSAAFTDGPLRHPNLEGIEVFE
EEMVVMTAAGHPPVTRPQDVNGKTIYAFRANCSYRRHFEEWFMSDQASPGKIFEIES
YHAMLACVTAGAGVAMIPRSMLE
SMPGSTNVSVYAIEEKFRHLTTWLVWRNGVRSPNLNAFIE
>185_Pseudomonas citronellolis GltR

MDLVQLEIFRAVAEHRSVAAAAQVMHRVPSNLTTRIKQLEADLGADLFIRENNRLRLSP
AGHEFLGYATRILELVGEARA
AVSGDQPIGRFPLGSLESTAAVRIPQILARFHRHYPQIELDLSTGPSGEMVDGVLDGRL
VAAFVDGPIEHPALEGVPVFD
EEMLVVAASHHGPIGRGRDVQGEMIYVFRQNCSYRRHFERWFAEDGATPGQVRELE
SYHGMLACVSAGGGLAVMPRSMLQ
SMPG
>186_Pseudomonas citronellolis PtrR
MDLTQLEFFRAVAQDGSITAAAGRLHRVPSNLTTRIKQLESELGAELFIREKSRLRLSPT
GRSFLDYAERILDLVEEARQ
VASGAEPHGSFSLGSMESTAAVRIPELLARYHQRFPKVQLDLSTGPSGEMIDGVLSGR
FIAAFADGPANHPQLDGRPVFQ
EELVLMSAQGHPPVRDARDVAGETVFAFRDTCSYRKRLEGWFAEQRVVPGKIMEME
SYHGMLACIAAGGGVAIIPKAMFD
SLAGGRNVTIHHLALSHAEATTWLFWRRGTDTPALRAFI
>187_Pseudomonas cavernae PtrR
MELSQLRIFQAVAEEGSIAKAAVRMSRVPSNLSTRLRQLEESLGVELFLRERQRLQLSA
AGKVLQSYAERLFALQAEAEA
ALKGGEPAGDFTIGSMYSTAAIHLPAMLACYHRAYPAVNLQVQTQPSGELVEGLIGGR
LDAALIDGPMEYAELDGLPLFE
EELVLITAAGHGPVASAADVAGESVYTFRPRCSYRRRLEAWFAASRVAMGQIMEIESY
HSMLACVVAGGGVALVPSSMLA
SLPGRESVDAHALAAPFDRTTTWLMWRKGMLGANLRAWIE
>188_Pseudomonas sp. RIT623 PtrR
MEFSQLRIFQAVAEEGSVTRAAERLHRVPSNLSTRLRQLEEQLGVELFLRERQRLQLS
PAGKVLLDYANRLAALRDEAMA
AVQGGQPAGDFLLGTMYSSAATHLPALLARYHQAYPAVNLQVRAAPSGELLEGLLGH
TLDAALVDGPPSLAGLDGVPLCD
EVLVLITSPEHPPVQTARDVAGKAVFTFRQGCSYRMRLEAWYAHAHTPMGRVMEIES
YQSMLACVIAGAGVALMAQSMLD
SLPGHERVRVHRLQAPFDQAQTWLMWRQGMCGANLQAWID
>189_Streptomyces acidiscabies
MEFAQLEMFVAVAEHGGITAAAEKLLRAPSNVSTRIRHLEKELGMDLLLRDKRQVTLSA
EGEAFREYARRLIDIADEAKS
LANGERPHGRFRIGALESTSAVRIPSLLAEFHLSYPEVDLELEAGASGMLFDHVLKGQV
SAVFTDGPPASPVLTGMHAFT
ENLVILTPEPVKTIDDDFRQSNPTVFLFGTVCSYRQRFEDWLEESGIKPGRMVEISSYY
SMLACVAAGAGICMMPRCLYE
SLPGSNRVTCHPIKGDLGRAETWLTWRRDARSANLRAFVRQVRA
>190_Erwinia piriflorinigrans PtrR
MDLTQLEIFCAIAEQGSVSAAAEKLHRVPSNISTRLKQLEEELGCELFRREKLRLHITDS
GRTFLAYSQRILALSEEAKQ
SVSSQRPGGIFTLGAIESTAAVRLPPLIARYHQSWPQVGLDLSTGPSGDMIDGVLAGR
YSAAFVDGPPKHPQLEGMPVFA

EELVLIGALNHPPVSDASGISGSTIYAFRANCSYRRLFENWFARTDAVPGKIFEMESYH
GILACVSAGAGLALIPSSMLE
SMPGRDSVNAWPLSEGTGQISIWLVWRKGVHSANLQAMKTLLTAD
>191_Pseudomonas sp. 14P_8.1_Bac3 PtrR
MDLTQLRLFQTVAEEGSITAAAERLHRVPSNLTTRIKQLEQELGVDLFIREKLRLRLSAT
GWTFLGYTRRILELVEEARQ
ATSGTEPQGIFTLGSLESTAAVRIPSLLADFHQCYPKVELDLSTGPSGDMIEGVLSGRL
NAAFVDGPLRHPELEGLKVFE
EEMVLITPSSHAPVSRAQDVSGKSVYAFRQNCSYRRHFEAWFKADLAAPGKIFEMES
YHGMLACVTAGAGVAMMPRSMLD
SMPGNRRVNAYPLVEAFRYLNIWLTWRRGIRSPALNAFIE
>192_Luteibacter sp.
MQLSQLRAFCAVAETGSVIGASRVLNRVPSSITVRIQQLEEDLGCELFVRERQKMSLS
HEGRKLLEHAQRLVGLADGTRA
FMREEDAEGHLVLGALGVTQVGFLPRLIARYRSRHRAVTLDVRSEASEVLVEQVADGV
LDIALVDGPVSTAALESRLAYT
DDMLLVTEFGHPVVESPRDLRGMQLFGSRHDGSFRYRIDAWLEAAGRQLLPVVEVES
YHAMLSRVVAGMGAAWIPRSVLP
TLAAHAHVRAHALGDVGRTEIHFLWRIGQ
>193_Pseudomonas bharatica PtrR
MELSQLRIFQAVAETGSVTRAAEQLHRVPSNLSTRLRQLEEQLGVELFHRERQRLQLS
PAGKVLLDYAGRMLTLHDEALA
AVQGGQPAGEFVLGCMYSTAAIHLPDLLARYHKAYPAVNLQVQTAPSGELLEGLLAGR
LDAALLDGPLSLSGLDGVPLCE
ETLVLLTEPDHPPVRSARDVAGRAVFTFRKGCSYRMRLEAWYAHDHAAMGRAMEIES
YPSMLACVVAGSGVALMSQSMLD
SLPGRENVAVHRLQTPFDQATTWLVWRRGMRGANLDAWVN
>194_Erwinia psidii PtrR
MDLTQLKMFCTVAETGSLARAAELLHRVPSNLTTRLRQLEHELGMDLFIREKQRLRLS
PVGHNFLGYAKRILALSEEAMN
MTHNGEPGGNFALGSMESTAAARLPMLLSAYHQRYPTVALSLLTGTSAEIIERVRAGT
LSAALASGPILFNELNSCLAFT
EQMILISSLDHRPVYHPKDAAGKTLFAFGPGCSYRSRFENWFHNEGVQPGTFMAIQSY
HAMLACVASGGGLALLPESVLA
QLPAAGQVRRHTLPEAIACTTTYLVWRRDAFTPNIKAL
>195_Enterobacter sp. Bisph1 PtrR
MDLTQLEMFNAVAQTGSITQAAQKVHRVPSNLTTRIRQLESELGVDLFIRENQRLRLSP
AGHSFLRYSQQILTLVDEARM
VVAGDEPQGLFSLGALESTAAVRIPATLARYNQRYPRIQFDLATGPSGTMIDGVIEGQL
SAAFVDGPIMHPGLEGLPVYR
EEMMIVAPVSHETISRAAQVNGASIYAFRANCSYRRHFESWFHADNATPGKIHEMESY
HGMLACVIAGAGLALMPRSMLE
SMPGHHQVQAWPLAENWRWLTTWLVWRRGAKTRHLEAFIELLNAD
>196_Rahnella sp. PD12R PtrR

MNLSQLEMFRAVAETGSISAAAQRVHRVPSNLTTRIKQLESELGVELFIRENQRLRLSP
AGRNFLSYNNRILDLVEEARV
SVSGTEPQGIFSLGALESTAAVRIPALLARYHQEFGKVELALSTGPSGELLDKLVEGEL
EATFVDGPVLHPVLDGIPVYQ
EEMVIVAPLNHAAVTRGKEVNGEIIYAFRSNCSYRRHFESWFADDGAAPGKIYEMESY
HGMLACVIAGGGLALMPRSMLQ
SMPGS
>197_Rhodospirillales bacterium PtrR
MDLSDLRIFSTVVRQGGVTRAAERLHRVQSNVTTRIRQLEQDLGVALFIREGKRLHLAP
AGQVLLDYSDRVLALADEARL
AVQDPRPRGVFRLGAMDSTAAVRLPGPLAEYHRFYPDVVVELRTGNPTQLATAILAGE
LDAALVAEPIPDEPFEKIFAFE
EEPVIVAAADHPPVGKKGAIPRTILCFEHGCPHRKRLEDWYSRRGAMPERTIELGSYP
ALLGCVVAGLGVALLPRSVLST
FPESKRLSVHALPRGENRAETVLIWRKGAGSPNIQAL
>198_Escherichia sp. unclassified PtrR
MDLTQLEMFNAVAEAGSITQAAAKVHRVPSNLTTRIRQLETELGVELFIRENQRLRLSP
AGHNFLRYSQQILALVDEARN
VVAGDEPQGLFSLGSLESTAAVRIPATLAEFNRRYPKIQFSLSTGPSGTMLDGVLEGKL
NAAFIDGPINHTAIDGMPVYR
EELMIVTPQGHTPVTRASQVNGCNIYAFRANCSYRRHFESWFHADGAAPGTIHEMES
YHGMLACVIAGAGIALIPRSMLE
SMPGHHQVEAWPLGEQWRWLTTWLVWRRGAKTRQLEAFIE
